# Unveiling novel conserved HIV-1 open reading frames encoding T cell antigens using ribosome profiling

**DOI:** 10.1101/2024.11.12.623167

**Authors:** Lisa Bertrand, Annika Nelde, Bertha Cecilia Ramirez, Isabelle Hatin, Hugo Arbes, Pauline François, Stephane Demais, Emmanuel Labaronne, Didier Decimo, Laura Guiguettaz, Sylvie Grégoire, Anne Bet, Guillaume Beauclair, Antoine Gross, Maja C. Ziegler, Mathias Pereira, Raphaël Jeger-Madiot, Yann Verdier, Joelle Vinh, Sylvain Cardinaud, Stéphanie Graff-Dubois, Audrey Esclatine, Cécile Gouttefangeas, Marcus Altfeld, Laurent Hocqueloux, Assia Samri, Brigitte Autran, Olivier Lambotte, Hans-Georg Rammensee, Emiliano P. Ricci, Juliane Walz, Olivier Namy, Arnaud Moris

## Abstract

The development of ribosomal profiling (Riboseq) revealed the immense coding capacity of human and viral genomes. Here, we used Riboseq to delineate the first translatome of HIV-1 in infected CD4^+^ T cells. In addition to canonical viral protein coding sequences (CDSs), we identify 98 alternative open reading frames (ARFs), corresponding to small Open Reading Frames (sORFs) that are distributed across the HIV genome including the UTR regions. Using a database of HIV genomes, we observed that most ARF amino-acid sequences are likely conserved among clade B and C of HIV-1, with 8 ARF-encoded amino-acid sequences being more conserved than the overlapping CDSs. Using T cell-based assays and mass spectrometry-based immunopeptidomics, we demonstrate that ARFs encode viral polypeptides. In the blood of HIV-infected individuals, ARF-derived peptides elicit potent poly-functional T cell responses mediated by both CD4^+^ and CD8^+^ T cells. Our discovery expands the list of conserved viral polypeptides that are targets for vaccination strategies and might reveal the existence of viral microproteins or pseudogenes.

## Introduction

Open reading frames (ORFs) of the human genome have been long annotated using restricted criteria including the presence of canonical start (AUG) and stop codons, and the potential to encode protein longer than 100 amino-acids (aa) ^1^. Recent advances in detection methods challenged this definition revealing that thousands of small ORFs (sORFs) encode polypeptides or microproteins shorter than 100 aa ^2^. These sORFs are widely spread along the human genome, some overlap classical ORF, on a different frame (so-called alternative reading frames (ARFs)), but the majority is found within 5’ untranslated regions (UTR) of known genes ^3^. Although some microproteins encoded by sORFs have been shown to play fundamental roles, for instance, in DNA repair, mitochondrial functions, RNA regulation, *etc.* ^4^, the biological functions of most remain enigmatic. Nonetheless, the thorough characterization of sORFs greatly diversified the translation landscape in healthy and malignant cells expanding in particular the sources of cancer antigens.

To date the characterization of sORFs mostly relies on the ribosomal profiling (Riboseq) that allows unbiased assessment of actively translated mRNA sequences. However, only a limited number of studies correlated Riboseq data with the generation of peptides ^5^. Combining Riboseq and mass spectrometry (MS)-based immunopeptidomics several studies revealed that a fraction of peptides presented by major histocompatibility complex (MHC) molecules (known as ligandome) is derived from sORFs encoded polypeptides that are specific or overrepresented in tumour cells ^6–9^. Other studies demonstrated that the tumour immune microenvironment is likely to favour the expression of human endogenous retroviruses (hERVs) that are normally silent in healthy tissues and of out-of-frame polypeptides, both constituting antigen sources for T cell immunity ^10,11^. Remarkably, these findings and others ^12^ highlighted that noncanonical translation events of sORF located in 5’UTR or out-of-frame are likely favoured in the context of stress.

Viral infections represent a major stress to the cell whose translation machinery is mobilized for the translation of viral RNAs. Viruses acquired specific features to control viral mRNA translation such as sequences favouring frame-shifting or internal entry sites of ribosomes ^13^. Thereafter, not surprisingly during the last decades, peptides derived from alternative translation events were described in cells infected with influenza virus ^14–16^, murine leukemia virus (MLV) ^17^ and human immunodeficiency virus-1 (here referred to as HIV) ^18–20^.

The expression of these non-canonical viral peptides often referred to as cryptic epitopes (CE), was mainly studied in the context of peptide presentation by MHC class I (MHC-I) molecules to cytotoxic CD8^+^ T cells (CTLs). To date, in the context of viral infections, only a limited number of studies have combined Riboseq and immunopeptidomic approaches to characterize the landscape and origin of viral peptides presented by MHC-I molecules. These studies confirmed that MHC-I molecules present peptides from unannotated ORFs within ARFs of the human cytomegalovirus (HCMV) ^21^ and of the severe acute respiratory syndrome coronavirus 2 (SARS-CoV-2) ^22^.

In the context of HIV infection, the existence of peptides derived from ARFs (ARFP) was, so far, revealed using T cell-based assays ^20,23–26^. CTLs specific to peptides derived from ARFs were indeed detected during the acute and chronic phases of infection ^18–20,24^. ARFP-specific T cells seem preferentially abundant in people living with HIV (PLWH) with favourable clinical outcome expressing protective Human Leukocytes Antigen (HLA) alleles ^23^. Further underlining the contribution of ARFP-specific CTLs in the control of viral replication, several labs have shown that HIV adapts to and escapes from ARFP-specific T cell responses by introducing mutations within ARF sequences ^18–20,24^. Remarkably, in the macaque model of simian immunodeficiency virus (SIV) infection, a single mutation within an ARF-derived epitope was strongly associated with viral rebound ^27^. Bioinformatics approaches analysing the association between HLA polymorphisms and HIV sequence variations (HLA footprint) revealed that the virus might produce peptides from ARFs buried within sense (5’ to 3’) or anti-sense (3’ to 5’) frames of the viral mRNA ^18,19^.

In the present study, we used Riboseq to delineate the first translatome of HIV in infected CD4^+^ T cells. In addition to canonical viral protein coding sequences (CDSs), we identify 98 ARFs, corresponding to sORFs that are distributed across the HIV genome including the 5’UTR region. Using a database of complete and unique HIV genomes, we show that most ARF aasequences are likely conserved among HIV-1 clade B and C strains, with 8 ARF aa sequences being more conserved than the overlapping aa CDS sequences. Using T cell-based assays and mass spectrometry-based immunopeptidomics, we demonstrate that 43 ARFs encode viral polypeptides. In peripheral blood mononuclear cells (PBMCs) of HIV-infected individuals, ARF-derived peptides elicit potent and poly-functional T cell responses mediated by both CD4^+^ and CD8^+^ T cells. Our findings broaden the spectrum of HIV immunogenic antigens that might help in the design of efficacious vaccines and might reveal the existence of viral microproteins or pseudogenes.

## Results

### HIV Ribosome Profiling reveals 98 Alternative Reading Frames

To date, the existence of ARFs in the HIV genome has been highlighted using indirect approaches such as T cell-based assays and HLA footprint analyses. We thus intended to provide an unbiased assessment of actively translated viral mRNA sequences in HIV-infected cells. To this end, we infected the SupT1 CD4^+^ T cell line with HIV_NL4-3_, in three biological replicates, and analysed viral mRNA translation using Riboseq (Supplementary Fig. 1a). Twenty-four hours post-infection, the viability of the cells and the infection rates were analysed using a viability dye and a combination of anti-HIV-Gag and anti-CD4 antibodies. Combining Gag and CD4 staining allowed the direct detection of cells expressing HIV-Gag protein and the indirect assessment of cells expressing viral proteins that down modulate CD4 expression such as HIV-Nef, -Vpu or -Env ^28^. Among the three biological replicates, the viability of the cells ranged from 44 to 71% and the rate of infected Gag^+^CD4^-^ and Gag^+^CD4^+^ cells from 54 to 73 % (Supplementary Fig. 1b). The cells were harvested, cellular and viral polysomes extracted. The ribosome protected fragments (RPFs) were then isolated and sequenced (Supplementary Fig. 1a). Data were analysed using the RiboDoc tool ^29^ and inhouse bioinformatics pipelines. The length of isolated RPFs ranged between 25 and 35 nucleotides (nts). We analysed the periodicity of the reads by aligning RPFs to annotated start codons of human CDSs. The reads, in particular the 28- and 29-mers, uniformly aligned along the human CDSs with a periodicity of three nucleotides corresponding to codon triplets, thus strongly suggesting that the RPFs correspond to *bona fide* translation events (Supplementary Fig. 1c). The data from the three biological replicates were then combined and viral RPFs aligned to the HIV_NL4-3_ genome (Fig. 1a and Supplementary Fig. 2a, b). The ribosome footprints aligned only on the forward strand throughout the viral genome. Note that on Fig. 1, each reading frame is highlighted by a specific colour: green, red and blue for Frame 0 (F0), 1 (F1) and 2 (F2), respectively (Fig. 1a). For clarity, the RPF occupancy for the different reading frames is also presented in Supplementary Fig. 2b in 3 separate graphs. The number and density of RPFs were particularly high in the UTR regions of HIV RNAs (Fig. 1a). As expected, we observed patches of RPFs in frame and covering the CDSs of known viral proteins such as HIV-Gag and -Env (Fig. 1a, in green and red, respectively). The decrease in density and number of RPFs covering the CDS of HIV-Pol also highlighted the specific feature of the HIV-GagPol poly-protein expression, which is controlled by a -1 ribosomal frameshifting during translation elongation ^30^. Based on the number of RPFs and the length of the CDSs, we calculated the relative expression of each CDS (Fig. 1b). The relative expression of HIV-GagPol polyprotein was about 6% of HIV-Gag, which is consistent with the literature ^30^. Thereafter, our ribosome profiling of HIV translation identifies all known HIV CDSs and confirms specific features of the regulation of HIV translation such as ribosome frameshifting.

**Fig. 1.**
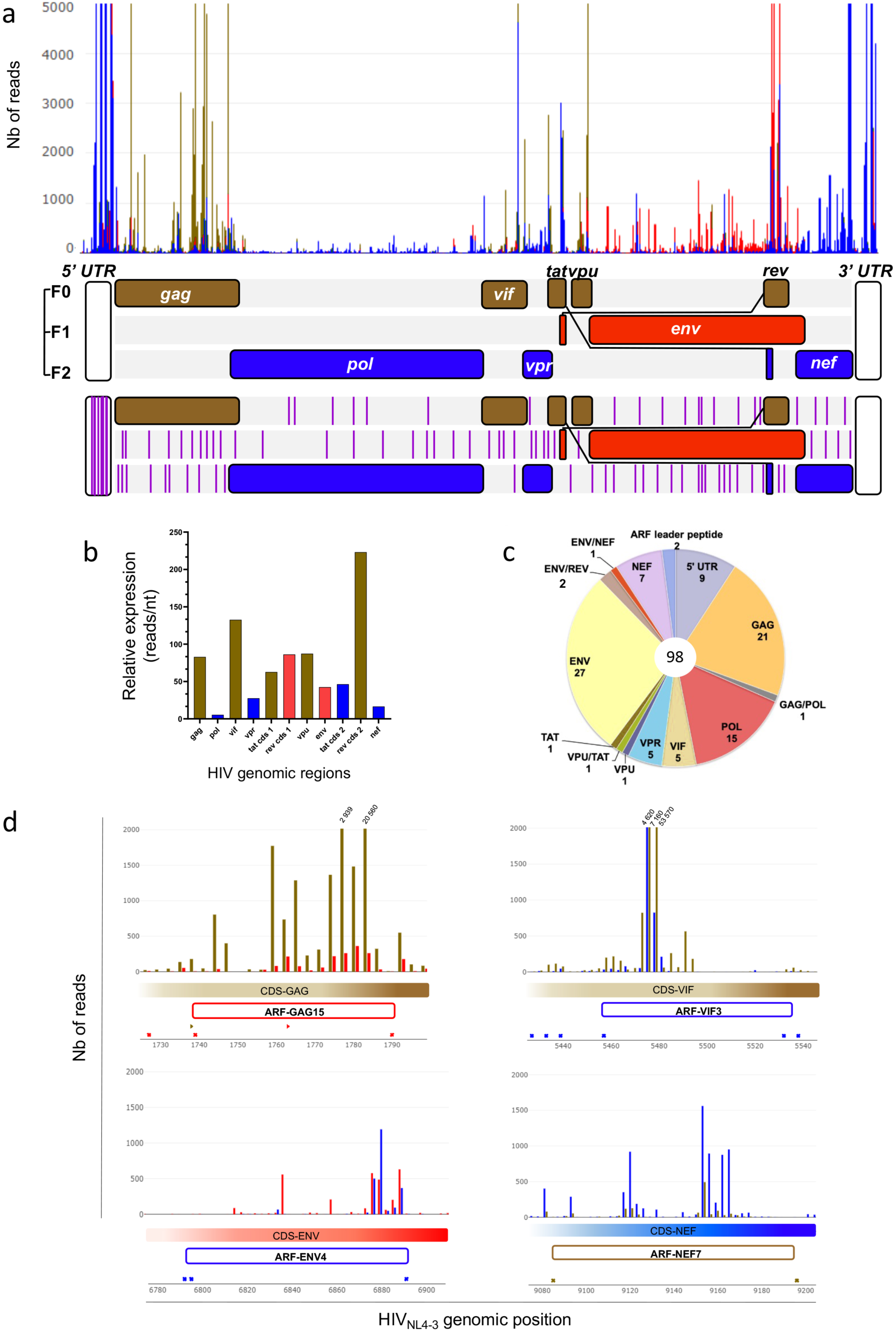
The translational landscape of HIV-1: identification of ARFs. **(a) Distribution of ribosome protected fragments (RPFs) across HIV-1 genome obtained from HIV_NL4-3_-infected SupT1 cells.** The number of RPFs (Nb of reads) are represented according to the position in HIV_NL4-3_ genome. For clarity, the numbers of RPFs are truncated above 5000 reads. To illustrate the translation signal in the UTR duplicated regions, multimapped RPF are shown. RPFs translated in frames 0, +1, and +2 are represented in green, red, and blue, respectively. HIV genomic RNA is represented below; CDSs are indicated and colored accordingly to their reading frames from the first position of the genome. Note that the classical reference strain, HIV_HxB2,_ owns an extra T (position 5772 within the *vpr* gene) as compared to most clade B strains. As a direct consequence in the HIV_NL4-3_ genome, env CDS is not in the same reading frame as pol and vpr CDSs. The CDSs from tat, rev, vpu and nef frames are also in a different frame than in the HIV_HxB2_ genome. The positioning of ARFs (purple lines) are indicated in the lower RNA genome representation. The selection criteria were: 1) not corresponding to CDS sequences; 2) aa length ≥ 10; 3) translated between 2 stops codons and 4) reads per codon ≥ 8. (b) Relative expression of HIV CDSs. Relative expressions of HIV_NL4-3_ CDSs were obtained by dividing the total read counts of each RPF by its length establishing a read/nucleotide value. As in (A), the color codes indicate the frames of in which the ARFs are localized (green, 0; red, +1; blue, +2). (c) Ribosome densities reveal novel viral coding regions. The number of RPFs (Nb of reads) are represented (y-axis) according to the position in HIV_NL4-3_ genome (x-axis). The color codes indicate the frames of the ARF and the relative CDS (green, 0; red, +1; blue, +2). Filled and open rectangles indicate the relative CDS and the ARF, respectively. ARFs within gag (top left), vif (top right), env (bottom left), and nef (bottom right) CDSs. Putative start and stop codons are labelled according to their reading frames with triangles and crosses, respectively. (d) Pie chart of the genomic distribution of the 98 identified ARFs according to the overlapping CDS. ARFs that overlap two CDS sequences are group into segments labelled with the name of the 2 CDSs. The number of ARFs is indicated below the name.

Strikingly, we also observed patches of RPFs overlapping known CDSs but aligning on other reading frames (e.g. patches of RPFs in frame-3 (red) overlapping HIV-Gag CDS or RPFs in frame-2 (blue) overlapping HIV-Vif CDS) suggesting translation of ARFs in the HIV genome (Fig. 1a and Supplementary Fig. 2). In our study having no information on potential start codons, we defined an ARF as a sequence between two stop codons with minimal size of 10 codons (shorter sequences being unlikely to be processed to produce MHC-ligands). In order to discriminate between background whole RNA sequencing and *bona fide* translation events, we then determined the number of reads in frame within each potential ARF, which allowed calculating a mean of RPF/codon for each ARF. We observed that 75% of all ARFs had less than 8 RPF/codon and decided to use this value as the minimum threshold to define *bona fide* translation of ARFs. Based on these criteria, we identified 98 HIV ARF sequences distributed across the viral genome (Fig. 1a bottom panel purple lines, 1c, 1d and Table 1). Examples of potential ARFs that failed or succeeded to pass our selection criteria are presented in Supplementary Fig. 2c. Note that ARFs were named based on the CDS that they overlap plus a specific number (e.g. HIV-GAG15). In Fig. 1c four examples of selected ARFs overlapping HIV-Gag, -Vif, -Env and -Nef CDSs are highlighted, named ARF-GAG15, ARF-VIF3, ARF-ENV4 and ARF-NEF7, respectively. Note that we also analysed the presence of these 98 ARFs in each dataset of the biological replicates: 86 ARFs were common to at least 2 replicates and 12 were found in only 1 of the replicates. Nonetheless, since these 12 ARFs exhibited a very high coverage, we decided to maintain them in our study. Interestingly, 16 out of 98 ARFs were previously described in the literature using indirect approaches (Table 1) ^18,19,25,26^. We also identified 9 ARFs in the 5’ UTR region of the HIV genome, commonly referred as uORF, which is consistent with previous studies describing translation events in the 5’ UTR regions of different viral mRNA ^31–34^. Note that we observed a large number of RPFs on the 5’ UTR region upstream the major splice site (SD1) (Fig. 1, Supplementary Fig. 2 and 3).

**Table 1:**
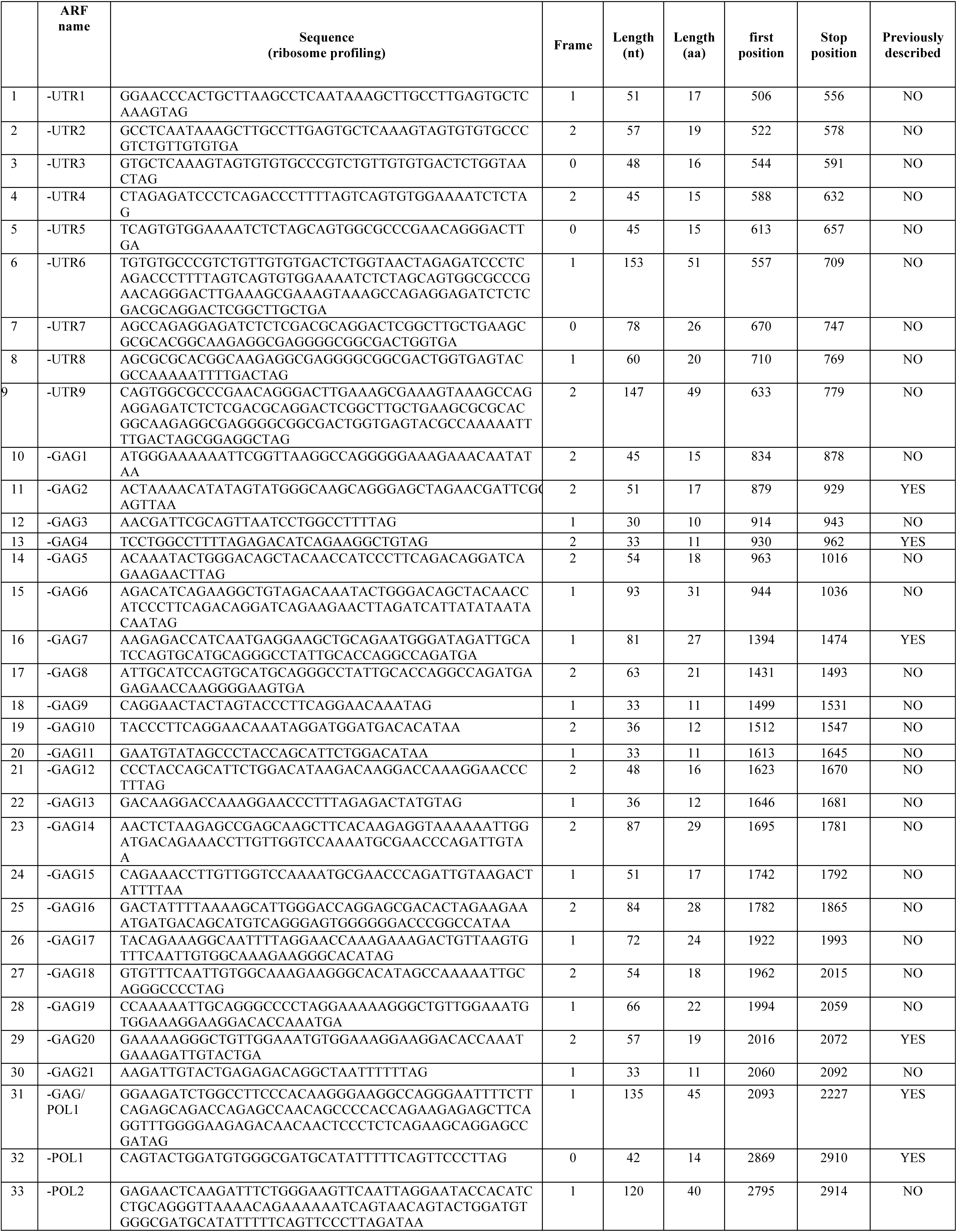
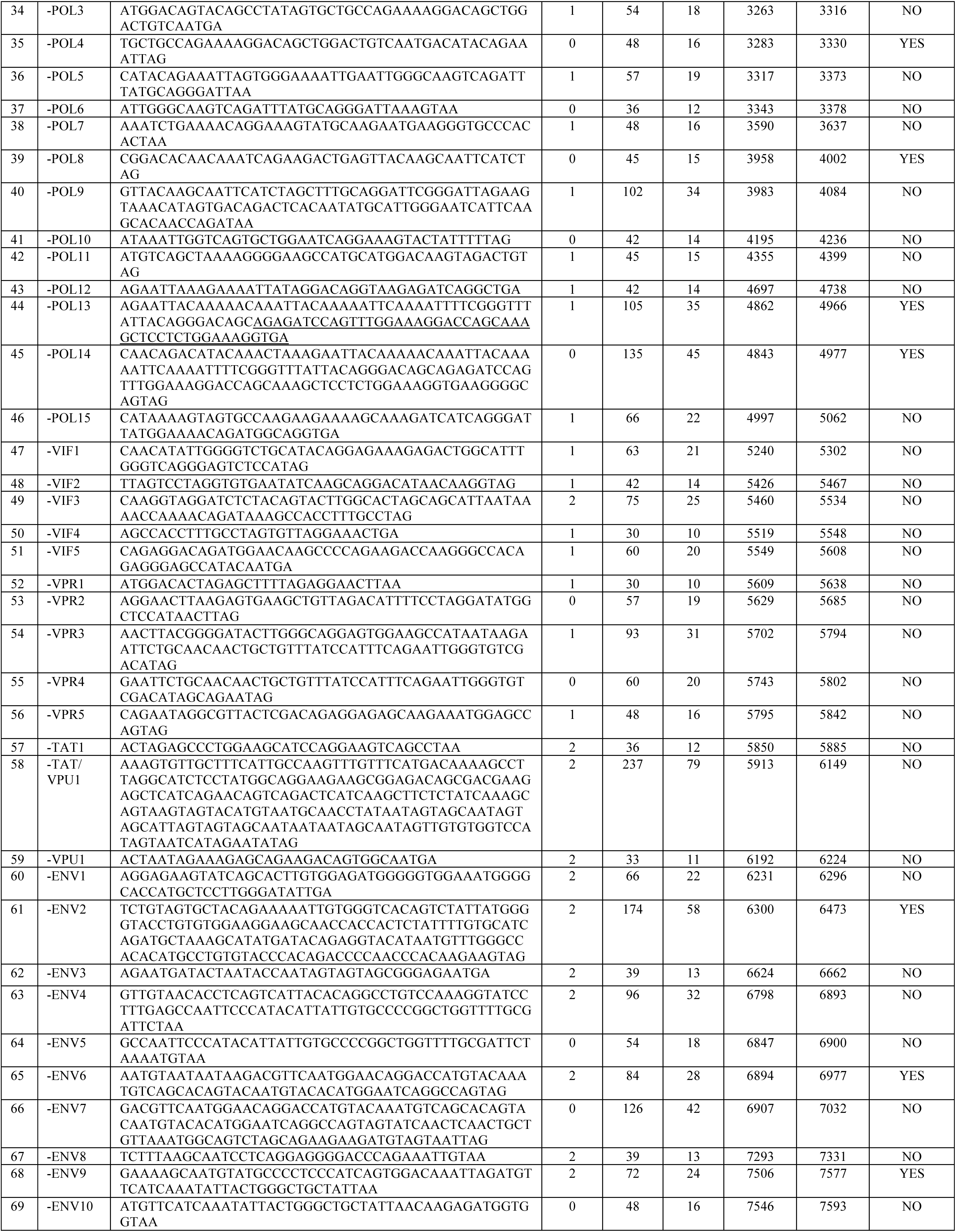
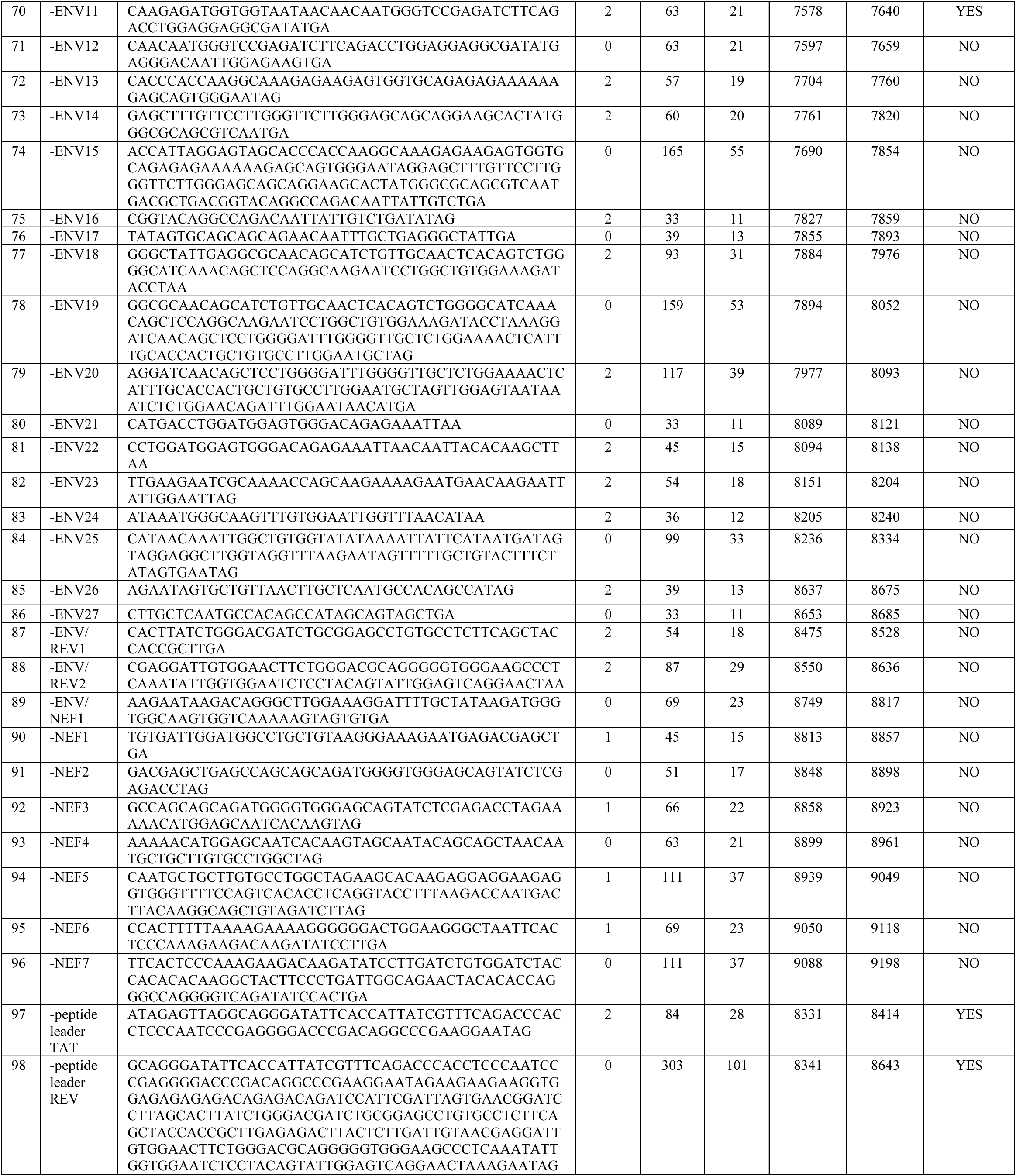
List of HIV ARFs identified by Ribosome profiling. Each of the 98 identified ARFs is listed according to their name, frame, length, and position in the HIV_NL4-3_ genome. Their description in previous studies is also informed.

In a second set of Riboseq experiments, using lactimidomycin (LTM), a drug that stops the initiation step of translation and thus leads to the accumulation of ribosomes at starts sites, we then define the start codons of each ARF. To further enriched ribosomes at start codons, LTM was combined with puromycin (PMY) to dissociates elongating ribosomes from mRNAs. As previously, SupT1 CD4^+^ T cells were infected with HIV_NL4-3_, in three biological replicates, cell viability and infection rates were analysed (not shown). Cells were treated with LTM and PMY prior harvest, RPFs isolation and sequencing. The RPFs of cellular mRNAs were analyzed for periodicity and metagene. Cellular RPFs were highly enriched at start codons, thus strongly suggesting that the combined treatment of LTM and PMY allowed the accumulation of ribosomes at start codons in treated cells (Supplementary Fig. 4). Viral RPFs were then aligned to the viral genome and RPFs in frame with the newly identified ARF analysed. We obtained heterogenous results with 71 ARFs where multiple RPF occupancy could be observed and 27 ARFs where it was not possible to distinguish them from background (not shown). Taking into account the three replicates, we observed that the mean intensity of RPF for all ARF was 62 reads and we decided to use this value as minimum threshold. We also excluded RPFs that accumulated 15 nucleotides downstream of stop codons. Based on these restrictive criteria, we listed the potential initiation sites that are shared by the three biological replicates (Table 2). For 26 ARFs, a single start position was identified while for the others one or two patches of RPF were observed (Table 2). Note that in the same ARF previously defined from stop to stop, translation might start at different initiating sites. Interestingly, among all potential initiation codons, 18 % were closely related to the classical ATG methionine start codon (near-cognate).

**Table 2:**
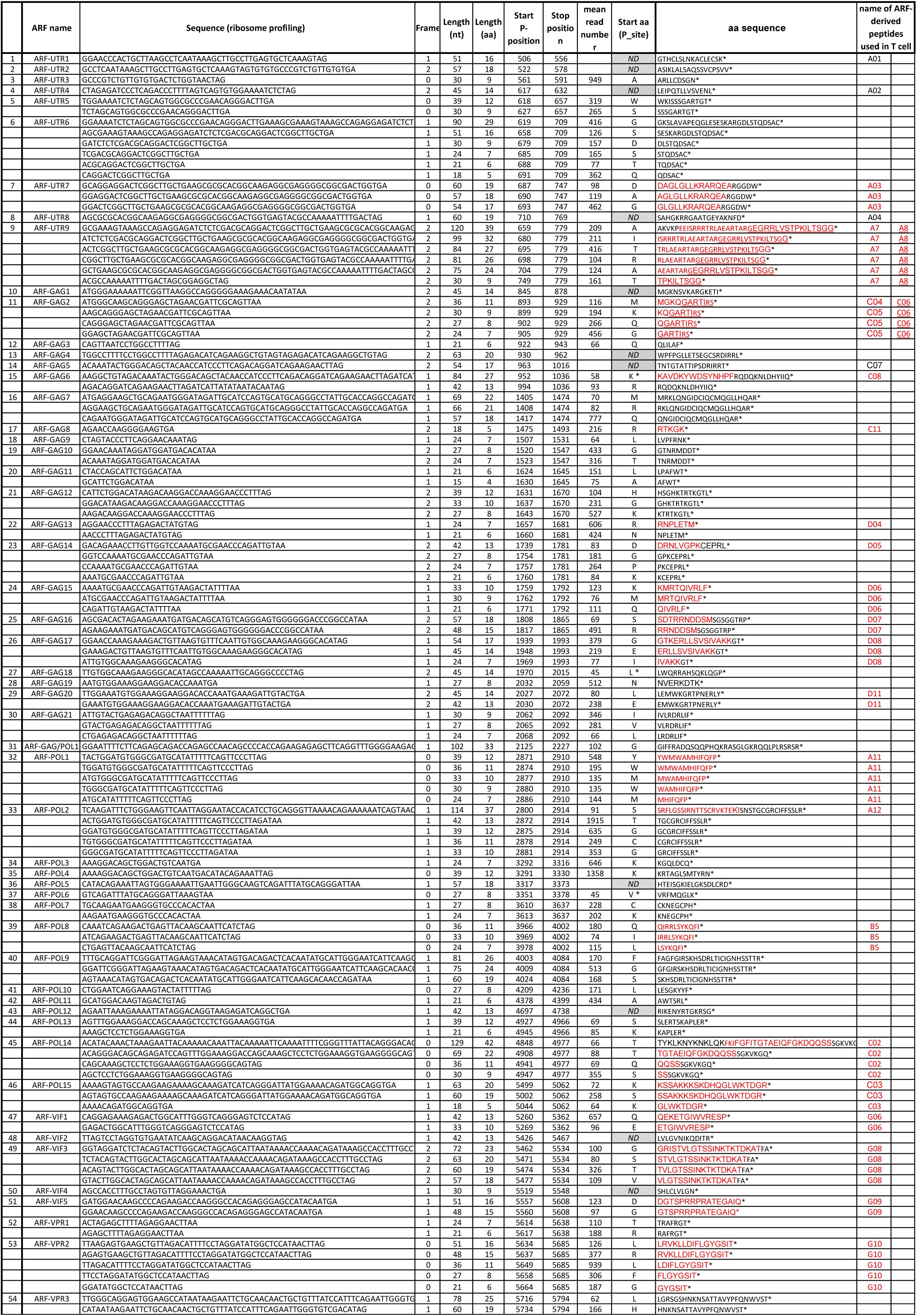
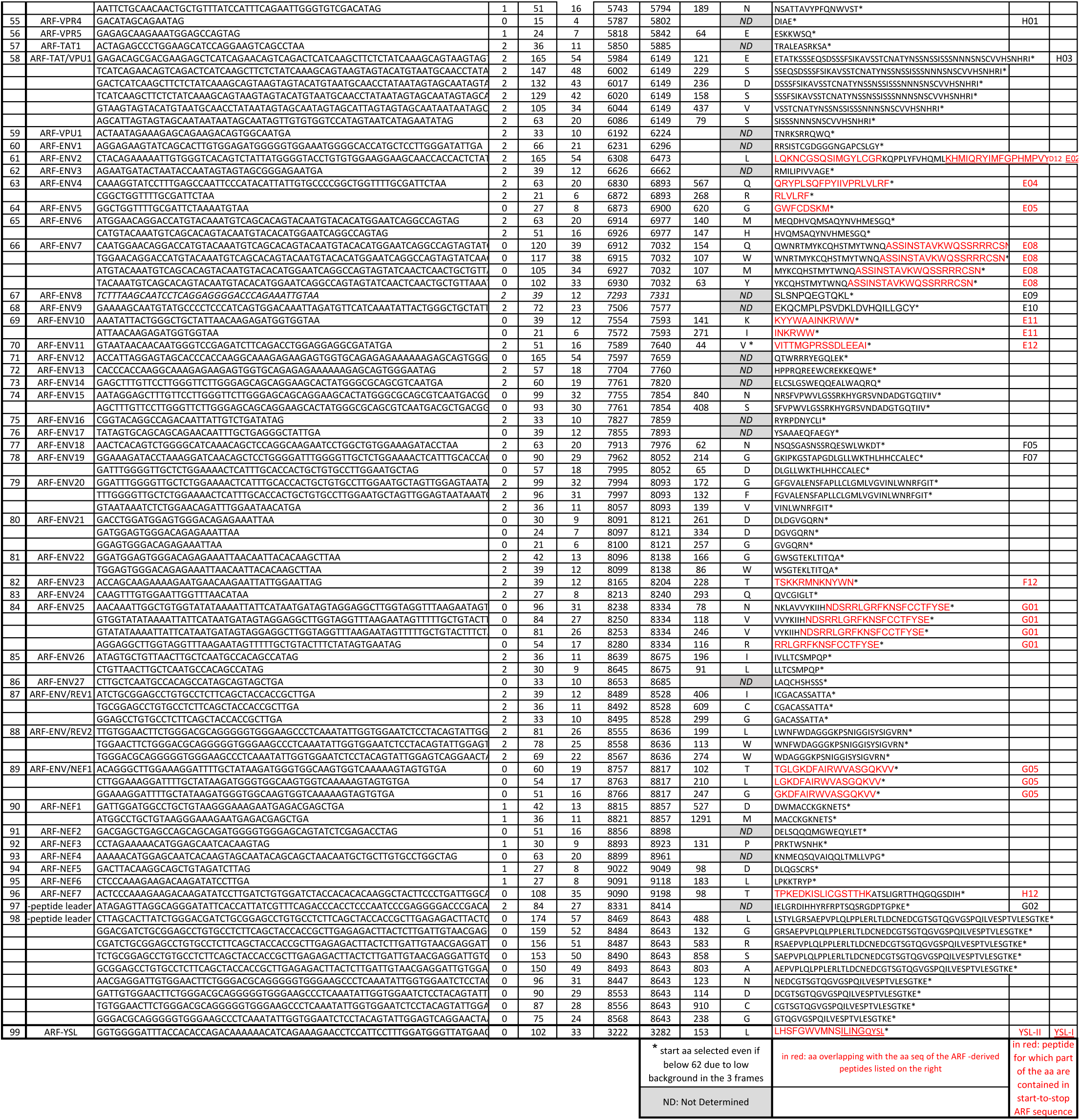
List ARF aa start positions. The table summarizes the data set of the Riboseq experiments performed with LTM+PMY-treated HIV_NL4-3_-infected SupT1 CD4^+^ T cells. As selection criteria, we used the mean intensity (62) of RPF across the 3 biological replicates. We also excluded RPF that accumulated 15 nucleotides downstream of stop codons. For each ARF, the name, the frame and the length of the nucleotide and aa sequences from the putative start to the stop codons are given. The position of the start and stop codons within HIV_NL4-3_ genome as well as the name of the start aa are presented. The number of RPFs at start position (P-site) of the is also indicated. The name of ARF-derived peptides used in T cell assays is also shown and the aa contained within the start-to-stop ARF sequence highlighted in red.

Using Riboseq, we therefore reveal the existence of 98 ARFs distributed across the HIV genome that are actively translated in infected CD4^+^ T cells. We confirm and extend previous work showing that other regions than annotated HIV CDSs are indeed translated.

### Identified ARFs are conserved among HIV clade B and C isolates

Assuming that these ARF-encoded peptides might be a reservoir of genetic novelty and a source of T cell antigens, we analysed their aa sequence conservation among clade B and C HIV isolates. Using Los Alamos HIV Sequence Database, we created in-house databases of clade B and C HIV sequences containing complete HIV genome without aberrant mutations within known CDSs and isolated from different individuals. We obtained 1609 and 411 clade B and C sequences, respectively. Using UGENE software, we then aligned the ARFs that we identified in the HIV_NL4-3_ genome to the databases (Fig. 2). Owing to the diversity of potential translation initiation sites for most ARFs, we decided to perform this analysis using the ARF aa sequence from stop-to-stop. The ARF-encoded peptide aa sequence conservation ranged from 48% to 98% with a median of 88% among the clade B (Fig. 2a) and from 42% to 98% with a median of 89% among the clade C (Fig. 2b). We did not notice a particular trend in aa sequence conservation when comparing the reading frames in which the ARFs are located in relation to the overlapping CDS (Supplementary Fig. 5a). In contrast, statistically significant differences in aa sequence conservation could be observed depending on the overlapping CDS (Supplementary Fig. 5b). The most significant difference in aa sequence conservation was found between ARFs overlapping *pol* and *env* genomic regions, which is consistent with the fact that *env* CDS is the most variable region. There was no difference in ARF-encoded peptide aa sequence conservation between clades B and C regardless of the reading frame from which the ARFs are translated (Supplementary Fig. 5c). Finally, we did not observe a correlation between ARF-encoded peptide aa sequence conservation and ARF relative expression (Supplementary Fig. 5d).

**Fig. 2:**
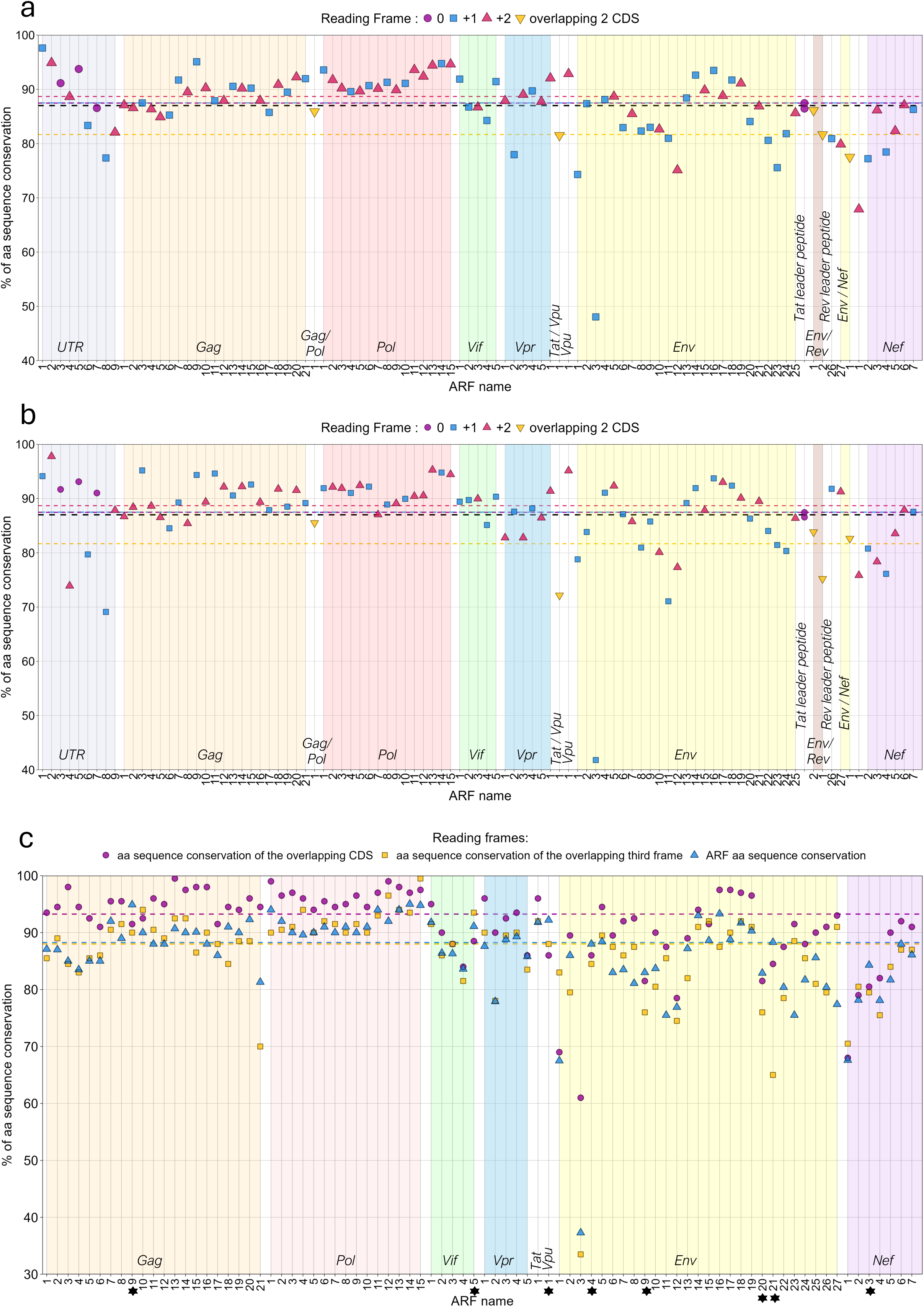
Amino-acid conservation of ARF-encoded peptide sequences among HIV-1 clade B and C strains. Percentage of amino-acid (aa) sequence conservation of each identified ARFs among HIV-1 clade B (a) and clade C (b). ARFs are represented according to the reading frame of their overlapping CDSs (top legend). ARFs expressed in +1 or +2 are represented by a blue square and a red triangle, respectively. ARFs overlapping two CDSs are represented by a yellow reverted triangle and those that correspond to peptide leaders, that are expressed in the first reading frame or in the 5’UTR having no CDS, are represented by a purple circle. The horizontal lines indicate: the global median conservation in aa of ARF-encoded peptides (black dotted line), the median for ARF-encoded peptides in 0 (purple dotted line), +1 (blue dotted line) and +2 (red dotted line) frames and the median for ARF-encoded peptides overlapping 2 CDSs (yellow dotted line). Genomic regions are indicated by background colors, with ARF names on the x-axis simplified for readability (e.g., ’1’ for ARF-GAG1), ranging from the 5’UTR (left) to the *nef* region (right). **(c) Percentage of ARF-encoded peptide amino-acid conservation and of overlapping genomic regions, among HIV-1 clade B.** Genomic regions are indicated by background colors, with ARF names on the x-axis simplified for readability (e.g., ’1’ for ARF-GAG1), ranging from the *gag* (left) to the *nef* region (right). The aa conservation of each encoded product of ARF (blue triangle), overlapping CDSs (purple circle), and third frame (yellow square) is presented. The colored horizontal lines indicate the corresponding median of the set. Black stars indicate ARF-encoded peptides that are more conserved than their overlapping CDS.

We then investigated whether it is the conservation of the CDS that imposes the conservation of the ARF. To this end, we analysed the percentage of aa sequence conservation of the overlapping CDS and of the third frame (*i.e*, corresponding neither to the CDS nor to the ARF sequences). To simplify the analysis, ARFs that overlap two ORFs were excluded. We observed a significantly higher conservation of CDS regions compared to the ARF and third frame, with a median of 93%, 88% and 88% aa conservation, respectively (Fig. 2c and Supplementary Fig. 5e). Interestingly, the most variable ARF among clades B and C, ARF-ENV3, overlaps the highly variable V1-loop of HIV-Env glycoprotein (Fig. 2) ^36^. These observations further highlight that ORF encoding the known viral proteins required for viral replication are under positive selection. In addition, there is no statistical difference in aa sequence conservation between the ARF and the third frames (median of 88% aa conservation for both, Supplementary Fig. 5e), which might imply that there is a low or no selection pressure on ARFs. However, it is particularly interesting to note that 8 ARFs sequences are more conserved than their overlapping ORF: ARF-GAG9, -VIF5, -VPU1, -ENV4, -ENV9, -ENV20, -ENV21, and -NEF3 (Fig. 2c, ARF names highlighted with stars).

Overall, our HIV sequence analysis strongly suggests that the 98 non-canonical transcripts identified by ribosome profiling are likely conserved among HIV clade B and C clinical isolates. Although, most ARFs are probably not under selective evolutionary pressure, eight ARFs are more conserved than their overlapping CDSs.

To determine whether all these ARFs readily encode viral polypeptides, we then used two complementary approaches, monitoring immune responses to ARF-derived polypeptides using PBMCs of HIV-infected individuals and using biochemical immunopeptodomics approaches to isolate directly, in infected cells, ARF-derived peptides.

### 42 ARFs encode viral polypeptides eliciting ARF-specific T cell responses

We first selected a set of peptides to be synthetized and then tested their capacity to elicit T cell responses using PBMCs of HIV-infected individuals, using a cultured IFNγ-ELISPOT assay (Supplementary Fig. 6a). ARF-derived peptides (ARFP) were selected based on their aa sequence conservation among HIV isolates and their predicted capacity to bind HLA molecules exhibiting high prevalence in the general population (Table 3). For ARFs with long potential aa sequences, several peptides were synthetized and tested (Table 3). The peptide length varied in order to include as much as possible multiple potential epitopes within the same aa sequences. We then monitored T cell-responses to these ARFP, using blood samples from individuals under anti-retroviral treatment (ART) or individuals who naturally control viral replication without treatment, so-called elite controllers (EC) (Table 4). In total, 96 different ARFP were synthesized and distributed in 9 different peptide pools (Table 3). As control, we included peptides derived from canonical HIV proteins (HIV) and from non-HIV common viral peptides (CEF). The HIV-canonical peptide pool was composed of previously characterized T cell epitopes (Supplementary Table 1) ^37^.

**Table 3:**
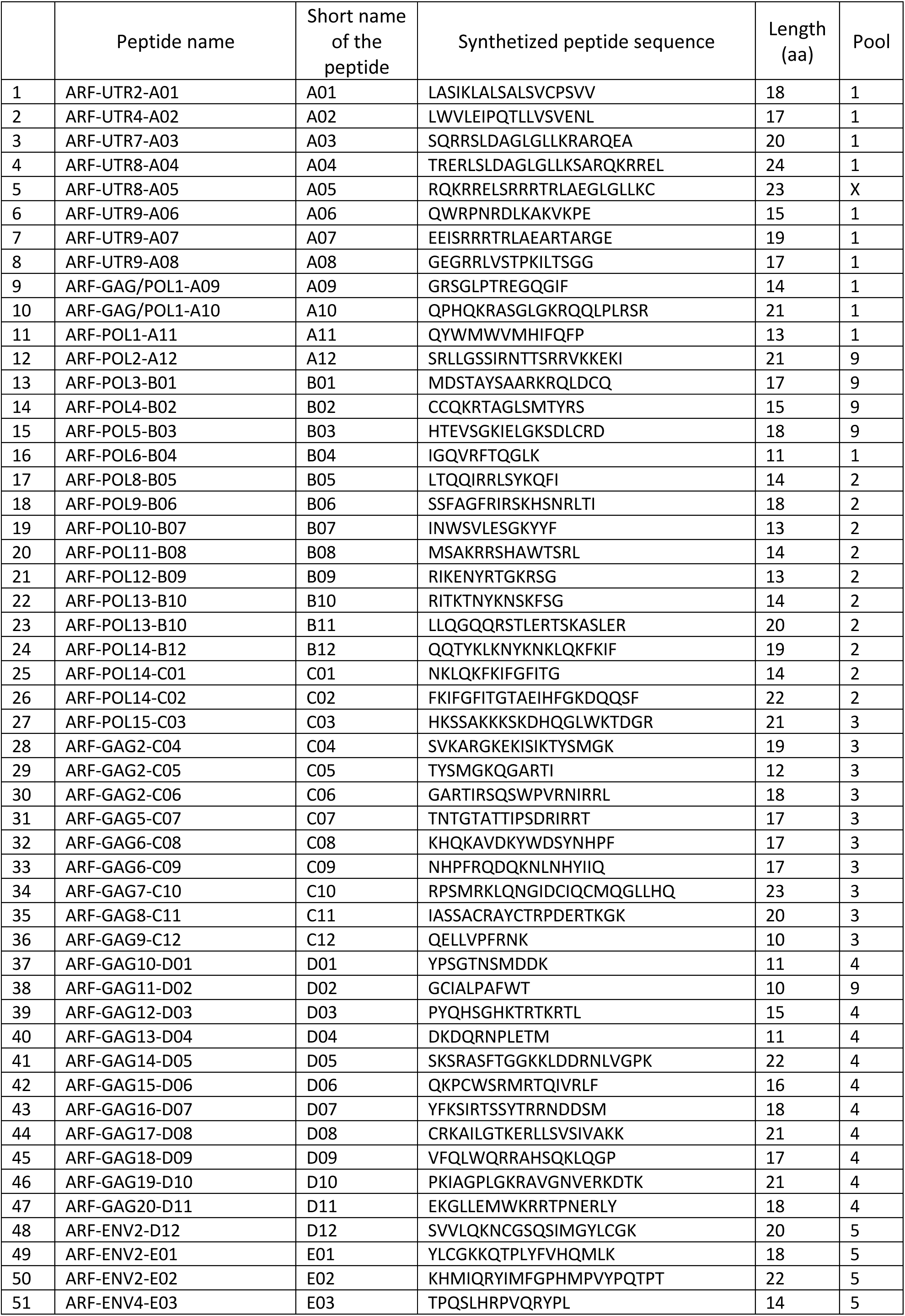
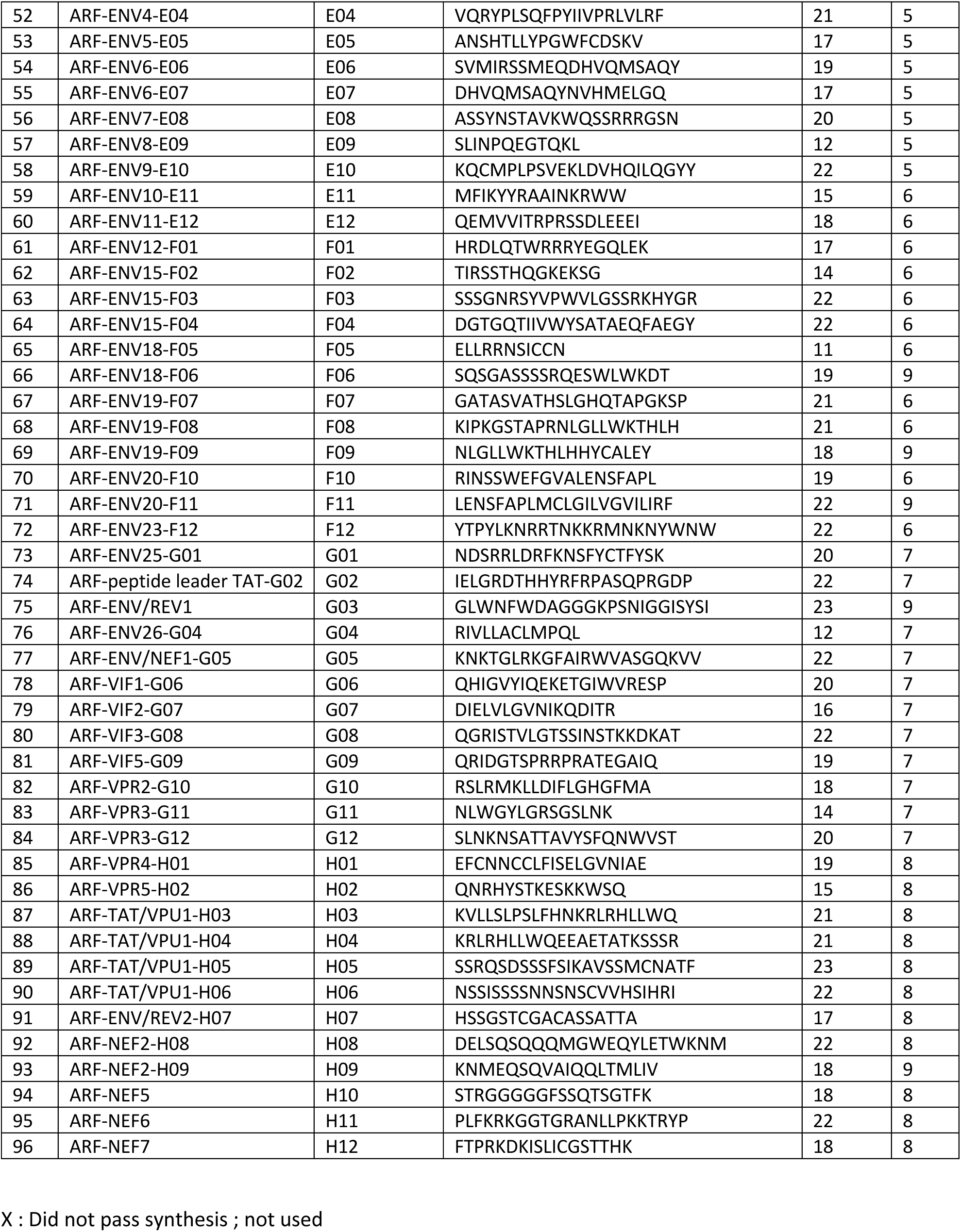
List of synthesized ARF-derived peptides. Peptides were named according to the ARF from which there are translated (column peptide name). The short name of the peptides is the peptide name used in the Figures. Peptide sequences and length are indicated. Pool: the number indicates in which pool the peptides were mixed.

**Table 4:** Clinical characteristics of HIV-infected individuals. Age, sex, years of infection, treatments, CD4^+^ T cell counts, viral loads, and HLA types of HIV-infected individuals are indicated. NA: Not available, F: female, M: male.

| Sample name used in the study | Age | Sex | Number of year of infection | Number of year of treatment | Treatment | Mean CD4+ T cell counts cells/mm3) | Viral load (copies) | HLA-A allele 1 | HLA-A allele 2 | HLA-B allele 1 | HLA-B allele 2 | HLA-C allele 1 | HLA-C allele 2 | HLA-DRβ1 allele 1 | HLA-DRβ1 allele 2 |
| --- | --- | --- | --- | --- | --- | --- | --- | --- | --- | --- | --- | --- | --- | --- | --- |
| EC-1 | 59 | F | 31 |  |  | 1106 | 40 | *03 | *30 | *57 | *15:63 | *14 | *18 | *01:02 | *13:01 |
| EC-4 | 59 | M | 30 |  |  | 647 | 61 | *26 | *33 | *27 | *51 | *02 | *12 | NA | NA |
| EC-2 | NA | NA | NA |  |  | NA | NA | *02 | *11 | *51:51 | *55:56 | *03 | *15 | NA | NA |
| EC-3 | NA | NA | NA |  |  | NA | NA | *1:01 | *68:01 | *35:03 | *37:01 | *04:01 | *06:02 | NA | NA |
| ART-2 | 61 | M | 22 | 22 | edurant + kivexa | 693 | <40 | *02 | *02 | *27 | *57 | *02 | *06 | NA | NA |
| ART-3 | 63 | M | 32 | 21 | atripla | 488 | <40 | *02 | *19 | *27 | *57 | *02 | *02 | NA | NA |
| ART-1 | NA | NA | NA | NA | NA | NA | <40 | *02 | *03 | *07 | *56 | *01 | *07 | *01 | *15 |

PBMCs from HIV-infected individuals were seeded in 24-well plates and loaded with the different peptide pools. Seven days later, a fraction of the cells was loaded with the peptide pool used for the initial culture and T cell reactivity was monitored using an IFNγ-ELISPOT assay (Supplementary Fig. 6a and b). Although with different magnitudes, the PBMCs from all donors reacted in a specific manner to the pool of peptides derived from canonical HIV proteins (HIV+) compared to the negative control (HIV-) (i.e cells from the same T-cell cultures but loaded with DMSO containing medium during the ELISPOT assay) (Supplementary Fig. 6b). Out of the 7 samples from infected individuals 6 reacted to at least one ARF-derived peptide pool (Supplementary Fig. 6b). Cells from donor EC-1 reacted to 8 ARFP pools. In contrast, we did not identify ARFP-specific response for donor ART-3, due to the high IFNγ secretion observed in negative controls (wells without peptide re-stimulation (-)). The response to the pool of classical HIV peptides was also particularly low in this donor (HIV+) (Supplementary Fig. 6b). Overall, several ARFP pools (including Pool-1, -2, -3, -4 and -6) induced very strong and specific IFNγ secretions in PBMCs from multiple HIV-positive donors (Supplementary Fig. 6b). The cells were then allowed to expand further by renewing the cytokine cocktail and tested on day 12, after the initial culture, for their capacity to react to individual peptides of the pools tested positive at day 7 (Fig. 3 and Supplementary Fig. 7). Note that despite the high background of IFNγ secretion observed at day 7, we tested the capacity of the cells from donor ART-3 to react to individual peptides of the pools 4, 6 and 8 because they showed a tendency to induce specific responses at day 7 (see Supplementary Fig.6, ART-3). In addition, T cell responses where the spots appeared visually particularly large at day 7, even if not exceeding the positivity threshold in terms of spot numbers, were also tested at day 12. The reactivity of each HIV-infected donor to the individual ARFP is exhaustively presented in Supplementary Fig. 7. ARFP-induced IFNγ^+^-T cell responses are summarized in Fig. 3 and pictures of the corresponding wells of the IFNγ-ELISPOT plates, are showed in Supplementary Fig. 7.

**Fig. 3:**
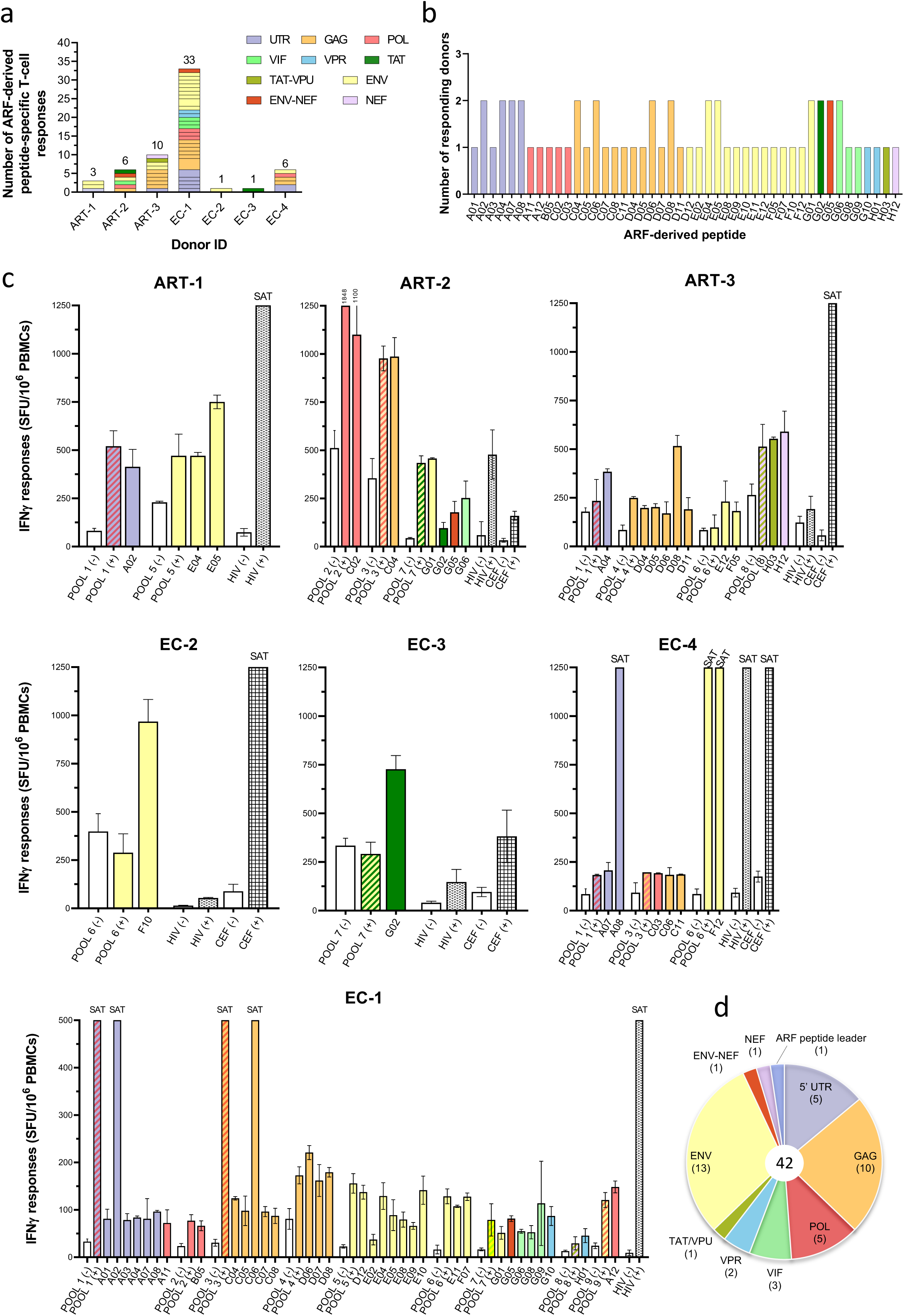
Detection of T cell responses specific to ARF-derived peptides (ARFP) in PBMCs of HIV-infected individuals. **(a) Overview of ARFP-specific T cell responses.** The number of ARFP-specific T cell responses is represented for each donor. The largest rectangle represents different peptides encoded by the same ARF. The color code, on the top right corner, indicates the overlapping CDS. (**b) Number of responding HIV-infected individual to each ARFP stimulation.** As in (a) the color code indicates the overlapping CDS. The rough data for IFNγ-Elispot responses are presented in Supplementary Fig. 7-related to Fig. 3 (**c) IFNγ-ELISPOT deconvolution of positive pools at day 12 and for all tested donors.** Only IFNγ-positive T cell responses upon individual ARFP stimulations are presented, in Spot Forming unit (SFU)/10^6^ PBMCs. Intrinsic negative controls (POOL (-), cells incubated with DMSO-containing medium) are indicated by white bars for each tested pool and the respective peptides. POOL (+), HIV (+), and CEF (+) correspond to PBMCs loaded with the pool of ARF-derived, HIV-classical, and non-HIV common virus-derived peptides, respectively. The color code indicates the overlapping CDS. SAT: saturated well for which the IFNγ signal was too high to be quantified. (**d) Genomic repartition of ARF encoding immunogenic peptides.** The number of ARFs is indicated in brackets. The color code is as in (A).

At day 12, with exception of EC-2, all samples from HIV-infected donors exhibited very strong or saturated responses to classical HIV-peptides. Remarkably, the PBMCs of all donors reacted to at least one individual ARFP (Fig. 3). Among the donors, the median of ARFP recognized was 6. However, responses were very heterogeneous with donor EC-1 reacting to 33 ARFP and EC-2 only to a single one (Fig. 3a). A total of 60 ARFP-specific T cell responses were identified across HIV-infected donor samples (Fig. 3a). Fourteen ARFP elicited T cell responses in samples from two different individuals (Fig. 3b). Altogether, the results using samples from HIV-infected individuals revealed 46 unique ARF-derived peptide specific T cell responses. The magnitude of T cell responses was heterogeneous between and among HIV-infected donor samples, with an intensity ranging from 13 to above 1250 (saturated) SFU/10^6^ PBMCs (Fig. 3c) with a median of 311 SFU/10^6^ PBMCs, but highly significant over background (Supplementary Fig. 8a). For instance, despite exhibiting a broad ARFP-specific T cell response, the magnitude of most T cell responses from donor EC-1 were low as compared with those from other donors (Fig. 3 and Supplementary Fig. 8b). Finally, we did not observe a significant difference in the magnitude of T cell responses between ART and EC individuals (Supplementary Fig. 8b). Remarkably, ARFP-specific-T cell responses target peptides encoded by ARFs distributed all along the HIV genome sequence from the 5’ UTR to *nef* CDS (Fig. 3d). Although the magnitude of uORF-specific T cell responses seemed particularly high (Supplementary Fig. 8c), ARPF-induced T cell responses were not significantly dominated by T cells recognizing ARFP from one specific HIV genomic region, neither UTR nor CDS (Supplementary Fig. 8d).

Using the exact same protocol, the entire set of ARF-derived peptide pools was also tested, at day 7, on PBMCs from non-infected donors (n=3) and on day 12, peptide deconvolution was performed on pools stimulating IFNγ-responses (Supplementary Fig. 9a and b). Cross-reactivity to one peptide pool was observed for EFS 1619 and EFS 5174 samples (Supplementary Fig. 9a) but the peptide cross-reactivity was not confirmed when testing individual peptides of the pool (not shown). Donor EFS 0685 exhibited a high background on all tested peptide pools. Peptide deconvolution showed a cross-reactivity to three peptides that were then excluded from the study (Supplementary Fig. 9b).

Overall, we demonstrate here that T cell responses from HIV-infected donors target 46 ARF-derived peptides encoded by 42 different ARFs. Of note for most ARFP that elicited T cell responses, the corresponding aa sequence could be found at least partially in the aa sequence encoded from the translation initiation sites, identified in our LTM+PMY Riboseq experiments (see Table 2 in red). We thus reveal that at least 42 ARFs identified in our ribosome profiling encode viral polypeptides and elicit T cell responses *in vivo* in the course of natural infection.

#### ARF-specific T cell responses are mediated by CD4^+^ and CD8^+^ T cells with a polyfunctional profile

We then asked whether CD8^+^ and/or CD4^+^ T cells mediate ARFP-specific T cell responses and studied the quality of these responses. The quality of HIV-specific T cell activation, defined as the capacity to produce multiple antiviral cytokines/chemokines, rather than the magnitude of T cell responses has been linked to disease outcome ^38^. Therefore, we characterized the capacity of ARFP-specific T cells to produce MIP-1β, IFNγ, IL2, CD107a, and TNF using Intracellular Cytokine Staining (ICS) (Fig. 4). PBMCs from 2 ART and 4 EC individuals (ART-2 was not tested in ICS due to a lack of sample availability) were seeded in 24-well plates and loaded with the individual ARFPs that induced T cell responses in the previous cultured IFNγ-ELISPOT assay (one ARFP per well). In total, we tested 47 ARF-derived peptides, 45 peptides which induced IFNγ-responses in the cultured IFNγ-ELISPOT assay plus 2 ARFP (one new ARFP (A05) and one (C12) inducing a slight but not significant T cell response in the first ELIPSOT) (Supplementary Table 2). In addition to the HIV+ peptide pool (HIV+), we included immunodominant peptides derived from classical HIV proteins with known HLA binding capacity ^37^. This pool of immunodominant peptides was adapted to the HLA allotype of each individual (Supplementary Table 3). At day 12, peptides inducing an IFNγ secretion were then used to monitor T cell activation using ICS combined with CD4, CD8 and CD3 extracellular staining (Fig. 4).

**Fig. 4:**
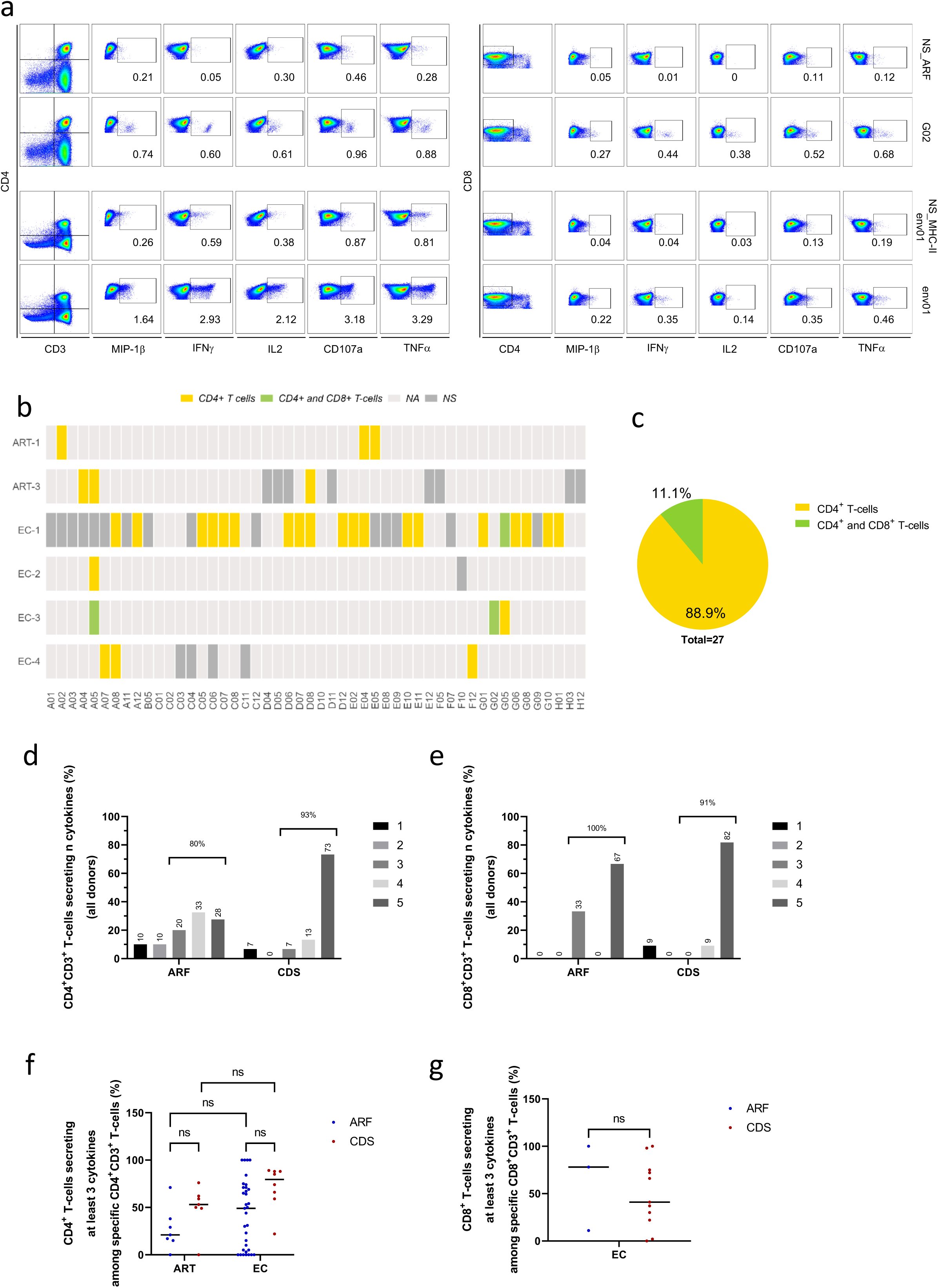
ARF-derived peptide (ARFP)-specific T cell responses are mediated by CD4^+^ and CD8^+^ T cells exhibiting polyfunctional cytokine profiles. **(a) Representative gating scheme for the identification of ARFP-specific T cell responses.** Upper panel, gatings for G02-ARFP-specific CD4^+^ (left) and CD8^+^ (right) T cell responses are shown for the EC-3 HIV-infected individual. Lower panel, gatings for Env01-specific T cell responses as for G02. Env01 epitope is a known immunodominant peptide from HIV-Env. CD8^+^ CD4^-^ T cell populations were pre-gated on CD8^+^ CD3^+^ T-cells, lived cells and doublets were excluded. Gates for each cytokine/chemokine were set based on the negative controls (cells loaded with DMSO containing medium, NS: Not stimulated) NS_ARF and NS_CDS for the assessment of G02-ARFP- and Env01-specific T cell responses, respectively. A figure exemplifying the gating strategy is provided in the Supplementary Fig. 13 **(b) Heatmap of ARFP recognized by CD4^+^ or CD8^+^ T cells from HIV-infected donors summarizing the ICS assays.** NA = Not applicable (i.e, not tested), NS = Not significant. (c) Percentage of CD4^+^ or CD8^+^ T cell responses among ARFP-specific T cell responses after *in vitro* stimulation with ARFP at the cohort level. Proportion of ARFP- and CDS-specific CD4^+^CD3^+^ (d) or CD8^+^CD3^+^ (e) T-cells secreting 1 to 5 cytokines simultaneously. The numbers on the top indicate the percentage of polyfunctional T cells secreting at least 3 cytokines. ARFP-specific T cell responses from all tested donors are combined. **(f) Comparison of the percentage of polyfunctional T cells secreting at least 3 cytokines among ARFP- and CDS-specific CD4^+^CD3^+^ T cells in the EC and ART patient groups. (g) same as (f) among ARFP- and CDS-specific CD8^+^CD3^+^ T cells in EC group.** Each dot represents one peptide-specific T cell response. The lines correspond to the median responses. Mann-Whitney test with Dunn’s comparison was applied. ns: not significant. From (d), data for YSL-I- and YSL-II-specific T cell responses are integrated into the graphs.

As illustrated with the ARF-derived peptide G02 and the HIV-Env derived peptide env01, peptide loading led to high production levels of MIP-1β, IFNγ, IL2, CD107a, and TNF by CD4^+^ T cells as compared to the negative controls (NS_ARF and NS_ORF, respectively) (Fig. 4a). Overall, 27 peptides induced the secretion of at least one cytokine (Fig. 4b). Most T cell responses, 89%, were mediated by CD4^+^ T cells (Fig. 4b and c). The ARF-derived peptides A05, G02 and G05 induced both CD8^+^ and CD4^+^ T cells (Fig. 4b).

CDS-specific CD4^+^ T cell responses were characterized by a significantly higher proportion of cells producing individual cytokines as compared with ARFP-specific CD4^+^ T cell responses (Supplementary Fig. 10a) which was not the case for CD8^+^ T cell responses (Supplementary Fig. 10b). However, although the frequency of responses against ARFP was lower than against CDS-derived peptides, ARFP stimulated polyfunctional responses in both CD4^+^ and CD8^+^ T cells (Fig. 4d and 4e). Since tri-functional T cell responses are often considered polyfunctional and might be associated with protection from disease progression ^39^, we focused our analysis on activated T cells with a polyfunctional profile with at least 3 functional responses. Clustering the data, we observed that 80% of ARFP-specific CD4^+^ T cells secreted at least 3 cytokines simultaneously compared to 93% for canonical HIV-specific CD4^+^ T cells (Fig. 4d). Ninety one percent of CD8^+^ T cell responses directed against canonical HIV peptides and 100% of ARFP-specific CD8^+^ T cell responses harboured a tri-functional phenotype, respectively (Fig. 4e). Profiles and frequencies of cytokines responses induced by each ARF- and CDS-derived peptide for the 6 HIV-infected individuals are presented as heatmaps for CD4^+^ and CD8^+^ T cells (Supplementary Fig. 10c and d). It was interesting to note that upon G02 peptide stimulation, the proportion of activated T cells secreting 5 cytokines simultaneously was higher than for some canonical HIV or even CEF peptide stimulations (see donor EC-3 donor, Supplementary Fig. 10c). In ART and EC individuals, we did not observe a significant difference in the capacity of ARF- and CDS-encoded peptides to induce polyfunctional cytokine productions (at least 3 functions) among neither CD4^+^ nor CD8^+^ T cells (Fig. 4f and g). There is also no significant influence of the clinical status on the polyfunctional profile of ARF- and CDS-specific T cell responses (Fig. 4f).

Finally, since the cytokine combination secreted by T cells might influence disease outcome ^40^, we dissected ARF- and CDS-specific T cell cytokine secretion patterns. To this end, we analyzed the 32 possible cytokine combinations (out of the 5 cytokines detected: MIP-1β, IL-2, INFγ, CD107a, and TNF) secreted by CD4^+^ and CD8^+^ T cells upon ARF- or CDS-derived peptide stimulation. Although the frequencies of CD4^+^ T cells targeting ARFP were lower than that of CDS-derived peptides, activated CD4^+^ T cells exhibited similar cytokine secretion patterns whether targeting ARF- or CDS-derived peptides (Supplementary Fig. 10e). The majority of polyfunctional ARFP-specific CD4^+^ T cells produced TNF, CD107a, IFNγ, and MIP1β simultaneously. Among ARF- and CDS-specific CD4^+^ T cells, the most prevalent monofunctional categories were TNF^+^ and Mip1β^+^ cells, respectively (Supplementary Fig. 10e). ARF- and CDS-specific CD8^+^ T cell responses displayed similar pentafunctional profiles (Supplementary Fig. 10e).

We then sought to analyze *ex vivo* ARF- and CDS-specific T cell responses in PBMCs of PLWH without *in vitro* expansion. To this end, cells from EC-1 and EC-3 were loaded with the individual peptides that induced a potent T cell activation in cultured IFNγ-ELISPOT and/or ICS (Supplementary Fig. 11). Remarkably for EC-1, 4 out of the 6 ARFP tested in ICS induced significant CD4^+^ T cell responses and one peptide stimulated CD8^+^ T cells (Supplementary Fig. 11a, left panel). For donor EC-3, the 4 ARFP induced significant CD4^+^ T cell responses (Supplementary Fig. 11b, left panel). Overall, the frequency of *ex vivo* ARF- and CDS-specific T cells was very low, limiting the analysis of the polyfunctional profile of the cells (Supplementary Fig. 11a and b, pie charts on the right panels). Nonetheless, bivalent and pentafunctional ARFP-specific *ex vivo* T cell responses could be detected for single ARF-derived peptides (Supplementary Fig. 11).

Taken together, our multiparametric study showed that ARF-derived peptides are predominantly recognized by CD4^+^ T cells from ART and EC donors and to a lesser extent by CD8^+^ T cells. Following *in vitro* T cell expansion, the percentage of ARFP-specific T cell activation was lower than CDS-specific T cell responses. Nonetheless, some ARF-derived peptides induced a higher magnitude of T cell activation than CDS-derived peptides. ARF-specific T cells were readily detected *ex vivo* in the PBMCs of HIV-infected individuals. ARF-derived peptide stimulation induced polyfunctional T cell activations.

### Identification of a naturally presented HLA-A*02:01-restricted ARF-derived peptide on infected primary CD4^+^ T cells

MHC ligands derived from sORF that are specific or overrepresented in tumour cells were recently identified combining Riboseq and immunopeptidomic approaches ^6–8^. In the context of viral infection, two studies also highlighted that HLA class I molecules can present peptides from ARFs of HCMV and SARS-CoV-2 ^21,22^. We thus intended to identify ARF-derived peptides presented by HLA class I and II molecules on the surface of HIV-infected cells (Supplementary Fig. 12a). We performed *de novo* experiments and also reanalyzed LC-MS/MS data from previously published immunopeptidomic experiments that used HIV-infected cells. To identify HIV-derived HLA-restricted peptides from all potential ARFs, the reviewed human proteome supplemented with the six-frame translated genome of HIV_NL4-3_, taking into account all potential peptides ≥ 8 aa encompassed between 2 stop codons, was used as reference. For our experiments we used the CD4^+^ T cell lines SupT1 (used in the Riboseq experiment) and C8166, which express HLA class I and both HLA class I and II molecules, respectively. In order to by-pass the down-modulation of HLA molecule expression mediated by HIV-Nef, we used a *nef-*deficient HIV_NL4-3_ isolate (HIV_NL4-3ΔNef_) ^41^. Cells were readily infected, with more than 50% and 20% of HIV-gag^+^ SupT1 and C8166 cells, respectively (not shown and Supplementary Fig. 12b). For both cell lines, although we obtained a large number of HLA class I ligands (5673 for SupT1, 7794 for C8166) and we detected known HIV-derived HLA class I ligands, we could not identify any ARF-derived HLA-restricted peptides. For C8166 cells, we additionally identified 4834 HLA class II-presented peptides from which 13 were derived from the HIV-Gag protein (Supplementary Fig. 12c). To our knowledge, these HIV-derived peptides are the first natural HLA class II ligands identified in infected cells. Nonetheless, we did not detect any ARF-derived peptide. We thus re-analyse the immunopeptidomics data from our previously published work using HIV_NL4-3_-infected primary CD4^+^ T cells ^42^ and we identified 7 HIV-derived peptides (Fig. 5a), one of which, ILINGQYSL, is derived from an ARF overlapping the *pol* gene (Supplementary Fig. 12e). Spectral validation using an isotopic labelled synthetic peptide confirmed the identification (Supplementary Fig. 12d). The ILINGQYSL peptide is annotated as a potent binder of HLA-A*02:01. Note that this ARF did not pass through the selection criteria of our initial Riboseq profiling because of its low coverage per codon (2.5 RPF/codon). However, it was readily identified in our Riboseq performed with LTM+PMY. LTM+PMY treatment led to the accumulation of RFPs at the leucine position 3222, suggesting that it might be the start codon of the ARF from which the ILINGQYSL peptide is derived (Table-2, bottom line). This leucine is highly conserved in strains from the clade B and poorly conserved for clade C (85 versus 15 % conservation) suggesting that other translation initiation events might occur in clade C isolates. Nonetheless, this ARF, is highly conserved (above 90% of aa conservation) among both HIV clades (not shown).

**Fig. 5:**
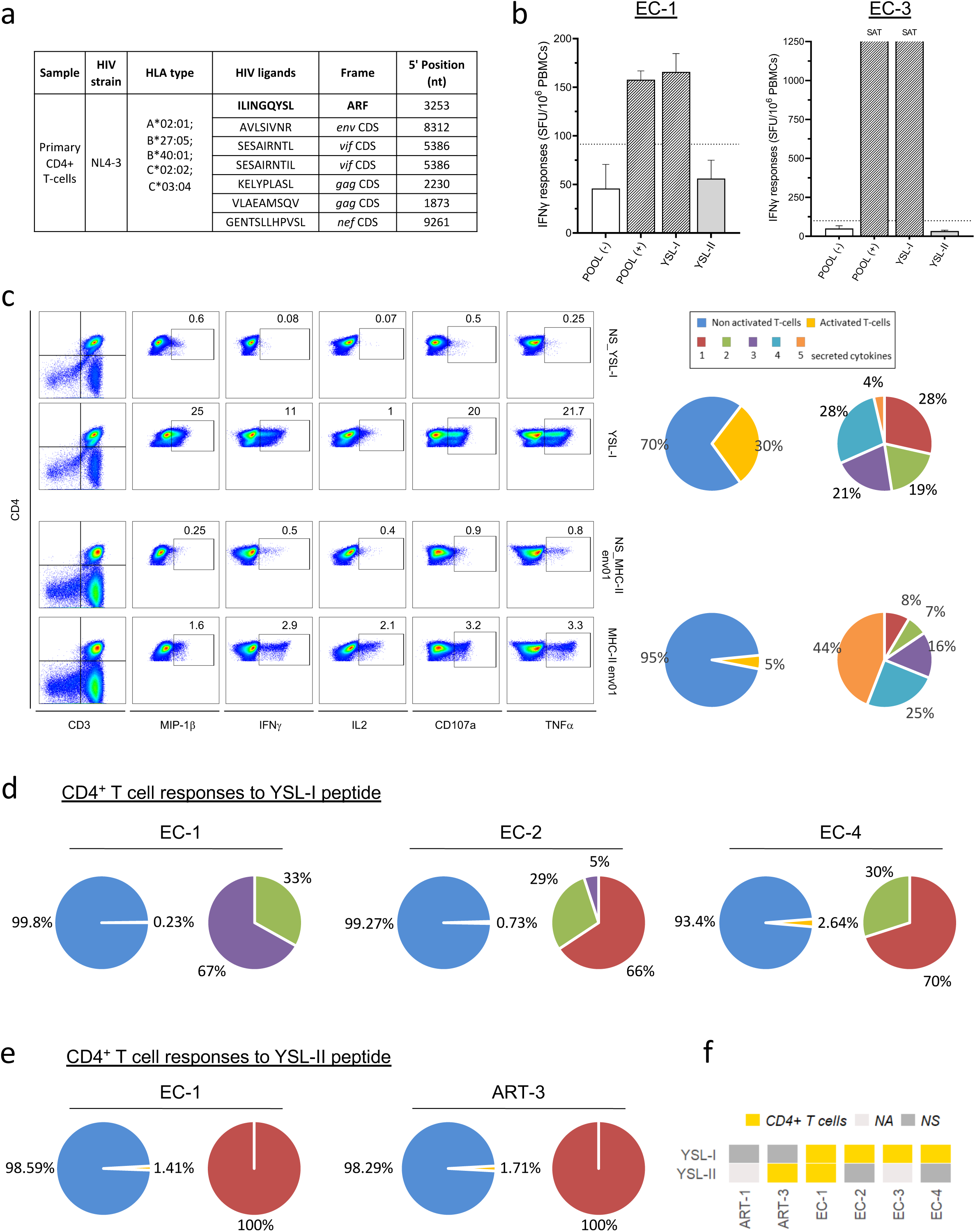
Mass spectrometry-based identification of CDS- and ARF-derived HLA-presented peptides and ARF-derived peptide (ARFP)-specific polyfunctional T-cells responses in HIV-infected donors. **(a) Identification of CDS and ARFP from HIV_NL4-3_-infected primary CD4^+^ T cells.** The sequences of identified HIV-derived HLA class I ligands and their characteristics are presented. **(b) Detection of T cell responses specific to YSL ARF-derived peptide in PBMCs of HIV-1 infected donors.** The IFNγ responses of donors EC-1 and EC3 who responded to the YSL-peptide stimulation, on day 12, are presented. Cells loaded with DMSO-containing medium, used as negative controls, (POOL (-)) are indicated by a white bar for each donor. POOL (+), YSL-I, and YSL-II correspond to cells loaded with a pool of peptides containing YSL-I and YSL-II peptides, or only with YSL-I and YSL-II peptides, respectively. A response was considered positive when the number of rough IFNγ^+^ spots per well was >20 and 2 times higher than the negative control (POOL (-) white bar, cells loaded with DMSO-containing medium). SAT: saturated well for which the IFNγ signal was so high that it could not be quantified. **(c) Functional characteristics of YSL-specific CD4^+^ T cell response from EC-3 donor**. Following a 12-day *in vitro* expansion, intracellular staining was performed for MIP-1β, IFNγ, IL-2, CD107a, and TNF after stimulation with YSL-I (top dotplot panel) or an Env-derived peptide (**MHC-II_env01,** bottom dotplot panel) used as positive control. Cells were gated on the CD4^+^ CD3^+^ population and each of the five individual gates for each cytokine/chemokine was set based on the negative controls (NS_, cells loaded with DMSO-containing medium). CD4^+^CD3^+^ cells were pre-gated on lived cells and doublets were excluded. Right panel, the left pie charts illustrate the percentage of activated (yellow) and non-activated (blue) T cells following YSL- or Env-peptide stimulation, upper and lower lines, respectively. The polyfunctional profile of activated YSL- or Env-specific T cells are shown in the right pie charts. Boolean analysis was conducted to determine whether T cells secreted 1, 2, 3, 4 or 5 cytokines simultaneously (red, green, purple, blue, and orange, respectively). Numbers indicate the percentage of activated cells with one to five cytokines secreted simultaneously. **Polyfunctional profiles of CD4^+^ T cell responses against YSL-I (d) and YSL-II (e) peptides.** The left pie charts show the percentage of activated (yellow) and non-activated (blue) T cells following peptide stimulation. The polyfunctionality of activated T cells, for all responding donors, is shown as pie charts on the right. The donor names are indicated on the top. Boolean analysis was conducted to determine whether T cells secreted 1, 2, 3, 4, or 5 cytokines simultaneously (red, green, purple, blue, and orange, respectively). Numbers indicate the percentage of activated cells with one to five cytokines secreted simultaneously **(f) Heatmap summarizing YSL-I and YSL-II peptide-specific CD4^+^ T cell responses in all tested HIV-infected donors.** NA = Not applicable, NS = tested but Not significant.

We then tested the capacity of the ILINGQYSL peptide (YSL-I) and an elongated form of YSL-I, LHSFGWVMNSILING (YSL-II), which has a high predictive score for binding to HLA-DR molecules (e.g., HLA-DRß1*03:01, *04:01, *07:01 according to the SYFPEITHI server), to activate T cell responses in PBMCs of HIV-infected donors. Using our cultured IFNγ-ELISPOT assay, we observed in PBMCs of EC-1 and EC-3 donors, intermediate to very strong T cell responses targeting the YSL-I peptide (Fig. 5b). The intracellular cytokine assay revealed that the YSL-I peptide is recognized by CD4^+^ T cells with a polyfunctional profile equivalent to responses specific to immunodominant CDS-derived peptides (Fig. 5c, pie charts). Responses to YLS-I peptide were particularly strong in CD4^+^ T cells from the EC-3 donor (up to 30% of CD4^+^ T cells) and were also detected, but to a weaker extend, in the cells from donors EC-1, EC-2 and EC-4 (Fig. 5c). Two donors also exhibited weak specific CD4^+^ T cell responses to the YLS-II peptide (Fig. 5e). Overall, in our cultured ELISPOT assay, 5 donors reacted to either YSL-I or YSL-II peptides (Fig. 5f). In addition, YSL-I-specific T cell responses with polyfunctional profiles were readily detected during *ex vivo* peptide stimulations in PBMCs of HIV-infected individuals (Supplementary Fig. 11), clearly demonstrating that the YSL-ARF readily encodes viral polypeptides.

Therefore, using two complementary and independent approaches, detection of ARF-derived peptide-specific T cells in the PBMCs of HIV-infected individuals and direct isolation of one HLA-bound ARF-derived peptide using mass spectrometry-based immunopeptidomics, we readily demonstrate that the HIV ARFs that we identified by ribosome profiling encode viral polypeptides capable of inducing broad T cell responses.

## Discussion

We provide here the first characterization of the HIV translatome in CD4^+^ T cells. We revealed the existence of at least 98 ARFs distributed across the HIV genome, including the 5’UTR, that are actively translated in infected CD4^+^ T cells. Most ARFs are likely conserved among HIV clade B and C clinical isolates. We demonstrated that at least 43 ARFs can be translated in viral polypeptides, since ARF-derived polypeptides were targeted by CD4^+^ and CD8^+^ T cells from HIV-infected individuals. Remarkably, ARF-derived peptides induced a polyfunctional T cell memory response that is reminiscent of the ones targeting CDS-derived immunodominant epitopes. Finally, we identified a conserved ARF-derived epitope, ILINGQYSL, naturally presented on HIV-infected primary CD4^+^ T cells by the HLA-A*02:01 molecule. ILINGQYSL is translated from a highly conserved ARF and induces *ex vivo* a polyfunctional T cell activation. Using two complementary approaches, we demonstrated that the ARFs identified are actively translated into viral polypeptides with the capacity to induce potent T cell immunity.

In human cells, recent advances have revealed that sORFs are widely spread throughout the human genome, some overlapping different frames of classical ORFs or locating within the 5’ UTR of known genes ^3^. Remarkably, these sORFs encode microproteins or polypeptides smaller than 100 aa ^2,4^. In fact, it has been estimated that 85% of the translation products originate from non-annotated regions of the human genome and mostly from out-of-frame sequences of CDS ^9,43^. We identified 98 ARFs within the HIV-1 genome, ranging from the 5’ UTR region to the *nef* gene. Nine ARFs are located in the 5’ UTR region. Several studies have recently demonstrated that polypeptides can be encoded in the 5’ UTR regions of viral genomes ^8,9,32–34,45^. These ARFs located at the 5’ UTR are probably translation regulatory elements of downstream CDS as observed in the human genome ^46^.

In our dataset, in addition to the 5’ UTR, most ARFs were located within *gag, pol* and *env* genes, which correlate with the size of the respective CDS. Previous studies, using HLA footprint or predictive ORF approaches, also found ARFs overlapping *gag, pol*, and *nef* sequences ^18–20,25,26^. Only 16 HIV ARFs identified in our ribosomal profiling were in common with ARFs already described in the literature ^19,25,26^. This relatively low number might be due to an over-estimation of the number of ARFs by the various prediction methods or the lack of thorough annotations of some ARFs in the previous studies. It can be also related to the rigorous criteria that we applied in our study for the selection of ARFs. This is illustrated by the ARF overlapping *gag* CDS and encoding the Q9VF epitope that we previously characterized in HIV-infected cells ^20,24^ whose coverage (5.9 RPF/codon) here is just below our selection criteria. However, we have previously shown that HIV_NL4-3_-infected cells are not recognized by Q9VF-specific CTL due to a proteasomal cleavage of the Q9VF epitope ^20,24^. Remarkably, we also did not identify RPFs aligning in frame on the minus strand of HIV. In particular, we did not identify the CDS encoding the antisense protein of HIV-1 (ASP) which overlaps the *env* gene, although several lines of evidence suggest that it is expressed *in vivo* ^47,48^. The lack of ASP CDS detection might be due to a specific regulation of translation from the 3’ UTR that could be down modulated in lymphoid cells such as CD4^+^ T cells used in our study ^49^. Nonetheless, we report here the identification of novel ARFs translated from the *vif, vpr, vpu, tat* and *env* sequences as well as ARF overlapping *tat* and *rev* exons.

Combining ribosome profiling with LTM and PMY, we intended to define the translation initiations sites of the newly described ARFs. We obtained heterogenous results: for 26 ARFs a single start site was identified while for the others one or two initiation sites were delineated by patches of RPF (Table 2). Interestingly, among all potential initiation codons, 18 % were closely related to the classical ATG methionine start codon. However, these results strongly suggest that a variety of mechanisms might be responsible for ARF expression. Indeed, a large part of eukaryotic and viral CDSs or ARFs initiate translation by non-canonical mechanisms such as leaky scanning, stop codon readthrough, ribosome shunting, re-initiation, IRES, frame shifting ^13^. Due to the diversity of potential translation initiation sites for most ARFs, we defined the ARF using flanking stop codons which also probably has an influence on the threshold of ARF RFP/codon and thus ARF identification.

Based on these criteria, the length of the identified ARF ranged from 10 to 101 aa and can therefore be considered as sORF ^50^. Using an in-house HIV database of clade B and C strains, we propose that the ARF-encoded aa sequences might be conserved among both clades. Overall, we observed that the overlapping CDSs are significantly more conserved than the ARF and the third frame, suggesting that ARF might be under a selective pressure weaker than classical CDSs. Remarkably, the ARF, encoding the ILINGQYSL peptide is highly conserved (above 90% of aa conservation) among HIV clade B and C which is probably explained by the fact that it overlaps the sequence encoding the catalytic site of the HIV reverse transcriptase (RT). This sequence of the RT is crucial for HIV replication and is among the most conserved region of the virus ^51^. Nonetheless, 8 ARFs appeared to be more conserved than their overlapping CDS sequence. We cannot exclude, thus far, that these 8 ARFs might encode polypeptides with functional viral proprieties.

uORF in the 5’UTR of HIV could influence Gag expression. This hypothesis has been addressed in the accompanying manuscript submitted together with Emiliano Ricci. Indeed, using mutagenesis and reporter systems, we show that extensive uORF translation from HIV-1 transcripts conditions the expression of Gag CDS. Other ARFs in HIV genome might also regulate the expression of the main HIV CDSs. They might also correspond to irrelevant by-standard translation events imposed by the tertiary structure and sequences of HIV mRNAs. To this regard, we observed a strong accumulation of reads, in the 3 frames, 30 nucleotides upstream of SD1, which is known to fold into a stable stem loop structure (Supplementary Fig. 3). Interestingly, this region is also translated from various frames since we could detect T cell responses targeting peptides derived from two different ARFs that overlay this patch of RPF (Fig. 3 and Supplementary Fig. 3). Overall, we believe that the strong patch of reads in this region might be due to several factors including, the presence of SD1 in all viral transcripts, the formation of stable hairpin loops (containing SD1 and the RNA dimerization motif (DIS) upstream of SD1) but also to translation of ARFs.

As mentioned, ARFs might also encode polypeptides with biological functions. Indeed, it is becoming evident that short polypeptides (below 100 aa), that were largely ignored so far, can exert various functions in development or viral infection. New identified ARFs might as well be seen as a genomic reservoir of unselected sequences allowing the emergence of *de novo* genes. Indeed, pervasive translation of short peptides derived from presumed non-coding regions might expose these ARF-encoded polypeptides to selection, allowing from time to time, the passage of ARFs into to the world of established, regulated, and selected products ^52^, as observed for ASP ^53^.

In infected cells, MHC-I epitopes originate from viral proteins but also from truncated or misfolded viral polypeptides, also called DRiPs (defective ribosomal products) ^54^. DRiPs are labile products degraded shortly after translation, allowing rapid loading of MHC-I molecules and thus CTL recognition within minutes after viral infections ^55^. Remarkably, DRiPs were initially identified in HIV-infected cells ^56^. As discussed above, it is likely that ARF produced viral polypeptides belong to DRiPs. In the present work, we readily demonstrate that at least 43 ARFs, identified by Riboseq, produce viral polypeptides which can induce T cell responses in HIV-infected individuals. Some of these ARF-derived peptides are targeted by CTLs. Highlighting the importance of ARFP-specific T cells in the course of HIV infection, others and we previously demonstrated that ARFP-specific CTL recognize infected cells ^18–20^ and exert a selection pressure on the virus *in vivo* ^18,19,24^.

To identify naturally presented ARF-derived HLA ligands in HIV-infected cells, we analyzed the immunopeptidome of two infected CD4^+^ T cell lines. Unfortunately, we could not identify any ARF-derived HLA-presented peptide. These might be due to several factors including the sensitivity of shotgun mass spectrometric discovery approaches. Indeed, even in the context of the immense technical improvements in the last decades, it remains for a challenge to identified low abundance peptides with high turnover rates. In addition, the immunopeptidome is a highly dynamic, rich, and complex assembly of peptides, which is shaped by several factors including accessibility to the antigen processing and presentation machinery *e.g.* specificities of proteasomes and cellular protases or of individual HLA allotypes. Nonetheless, our analysis of the immunopeptidomics data, published in our previous work ^58^, coming from infected primary CD4^+^ T cells allowed us to identify one ARF-derived peptide, the HLA-A*02:01-restricted peptide ILINGQYSL, thus providing a direct demonstration that ARF-derived peptides are naturally presented by HLA molecules. Probably due to the low number of HIV-infected samples that we used in our study, we did not detect CTL responses targeting this peptide. However, CD4^+^ T cell responses against this peptide or an elongated version of it was observed in five HIV-infected individuals. Therefore, our results suggest that alternative translation events may give rise to epitopes recognized independently or concomitantly by CD4^+^ and CD8^+^ T cells and presented on different HLA class I and HLA class II alleles. Overall, we observed that the majority of T cell responses targeting ARF-derived peptides were mediated by CD4^+^ T cells.

As a matter of fact, HIV-specific CD4^+^ T cells play an important role in HIV-infection. The breadth and specificity of HIV-specific CD4^+^ T cell responses are associated with improved viral control and low viremia during acute and chronic infection, respectively ^59,60^. In particular, the generation of Gag-specific CD4^+^ T cells and their maintenance correlates with a better disease outcome and viral control, respectively ^59,60^. There is also evidence of an association between certain HLA-DRß1 alleles and Gag-specific CD4^+^ T cell activity and delayed disease progression ^61^. HIV-specific CD4^+^ T cells can also exert direct antiviral functions by eliminating infected cells and/or inhibiting viral replication ^62^. Remarkably others and we have shown that CD4^+^ T cell can recognize peptides derived from newly-synthetized, so-called endogenous antigens in virus-infected cells, including HIV-infected cells ^63,64^ or in tumour cells ^65^. In fact, depending on the subcellular localization, the trafficking and the nature of the antigen itself, different pathways are involved in the degradation and presentation of endogenous antigens by MHC-II molecules. This includes components of the MHC-I processing pathway, such as proteasome and TAP ^66,67^, autophagy ^68^ and receptors regulating vesicular trafficking ^69^. In the field of cancer, it was recently reported that CD4^+^ T cells also recognize peptides derived from unannotated CDSs and neoantigens ^70–73^.

To get some insights on the quality of ARFP-specific T cell responses, we analyzed the polyfunctional profiles and the pattern of cytokine secretions of these CTL and CD4^+^ T cell responses in PBMCs of PLWH. We show here that CD4^+^ and CD8^+^ T cell responses against ARFP exhibit a polyfunctional profile with 80% of cells secreting at least 3 cytokines simultaneously. The pattern of secretion does not differ between previously described T cells responses targeting immunodominant epitopes and ARFP-specific T cells responses, suggesting that the latter are as potent as classical T cell response in controlling viral replication.

Although using a small cohort of PLWH with suppressed viral replication, we were intrigued by the large heterogeneity of ARF-specific T cells responses. Several parameters might influence the magnitude and the breadth of HIV-specific T cell responses including the individual HLA allotype and T cell repertoire, the senescence of T cells, the antiretroviral treatment and the size of the active viral reservoir, defined as cells carrying a transcriptionally active viral genome ^74^. Remarkably, the later has been recently linked to the magnitude and functions of HIV-specific CD4^+^ and CD8^+^ T cells ^75,76^. Interestingly, this active reservoir is mainly composed of defective viral genomes ^75^ which might favour the expression of alternative reading frames and thus the activation of ARF-derived peptide specific T cells. In the future it will be of interest to study ARF-specific T cells responses with regards to the active reservoir and the emergence of new dominant T cell clonotypes after prolonged ART treatment ^77^.

In conclusion, using ribosome profiling, we defined here the translatome of HIV in infected CD4^+^ T cells. We demonstrated that the HIV genome harbours ARFs that can be translated in viral polypeptides used by the immune system to initiate potent T cell responses. Both in cancer and in HIV-infected cells, ARF-derived polypeptides represent promising targets to promote the control of tumor growth or viral replication. Understanding how the translation of non-canonical genomic regions is regulated and its involvement in various cellular or viral processes will certainly help in fighting cancer and infections.

## METHODS

### Virus and infection

VSV-G pseudotyped HIV_NL4-3 XCS_ and HIV_NL4-3ΔNef_ were produced by transfection of 293T cells using CAPHOS kit (Sigma-Aldrich) as in ^63^. Viruses were pseudotyped to favor viral entry and to achieve a high infection rate within a short time. SupT1 CD4^+^ T cells were cultured with RPMI GlutaMax 1640 (Gibco) complemented with 10% FBS (Dutscher) and 1% penicillin/streptomycin. 1 x 10^9^ SupT1 CD4^+^ T cells were infected with 4 055 ng of VSV-G-HIV_NL4-3 XCS_ HIV-Gagp24 for 3h in IMDM plus 10mM HEPES (Gibco/ThermoFisher Scientific) supplemented with 2 µg/ml of DEAE-dextran (Sigma). Three biological replicates were performed corresponding to 300 million cells infected at day 0. Twenty hours post-infection a fraction of the cells (2 x 10^5^) was harvested, stained with CD4-Vio770 antibody (Miltenyi) and a Live/Dead (LVD) fixable violet dye (Invitrogen), fixed with 4% paraformaldehyde (ChemCruz) and permeabilized (PBS1X, BSA 0,5%, saponin 0,05%). Cells were then stained with an HIV-1 Gag-p24 specific antibody (KC57-PE, Beckman Coulter). Sample acquisition and analysis were performed using a BD LSRFortessa and FlowJo v10.8 Software, respectively (both from BD Life Sciences). Mock-infected cells were used as negative control. Twenty-four hours post-infection, cycloheximide (CHX) was then added to the remaining cell cultures (100 µg/ml, 10 min, 37°C, Sigma). Cells were washed in cold PBS containing CHX and lysed at 4°C (10 mM Tris-HCl pH7.5, 100 mM KCl, 10 mM Mg2+ acetate, 1% Triton X100 and 2 mM DTT). Glass beads were then added to the cell supernatant and incubated on an orbital shaker (5 min, 4°C). Beads were removed by centrifugation and the supernatant was quickly frozen in liquid nitrogen and stored at -80°C. To identify the start codons, prior harvest, cells were treated with Lactimidomycin (LTM; 5 μM, 30 min, 37°C) together with Puromycin (PMY, 25 μM) for the last 20 min. Cells were lysed in above mentioned lysis buffer containing LTM and PMY.

### Ribosome profiling

Cell lysats were gently thawed on ice in presence of EDTA-free complete protease inhibitor cocktail (Roche) and 200 U of RNase Murine Inhibitor (NEB) were added. The absorbance of the crude extract obtained was measured at 260 nm. The extracts were digested using 5U/UA_260nm_ RNase I (Ambion) for 1h at 25°C the digestion was stopped adding SUPErasin RNase inhibitor (500U, Invitrogen). Monosomes were then loaded on a 24% sucrose cushion and ultracentrifugated at 100 krpm on a TLA110 rotor at +4°C. The concentrated monosomes were resuspended in 600 µl of lysis buffer. RNA were extracted by acid phenol at 65°C, chloroform and precipitated by ethanol with 0.3 M sodium acetate pH5.2. Resuspended RNAs were loaded on 17% polyacrylamide (19:1) gel with 7M urea and run in 1xTAE buffer for 6h at 100V. RNA fragments corresponding to 28-34nt were retrieved from gel and precipitated in ethanol with 0.3 M sodium acetate pH5.2 in presence of 100mg glycogen. rRNA were depleted using Ribo-Zero Gold rRNA Removal kit (Illumina). The supernatants containing the ribosome footprints were recovered and RNA were precipitated in ethanol in the presence of glycogen overnight at -20°C. The RNA concentration was measured by Quant-iT microRNA assay kit (Invitrogen) and the RNA integrity and quality was verified using Bioanalyzer small RNA Analysis kit (Agilent). cDNA library from 100 ng RNA was prepared by the High-throughput sequencing facility of I2BC, using the NebNext Small RNA Sample Prep kit with 3’ sRNA Adapter (Illumina) according to the manufacturer’s protocol with 12 cycles of PCR amplification in the last step followed by DNA purification with AMPpure XP beads cleanup. Library molarity was analyzed using Bioanalyzer DNA Analysis kit (Agilent) and an equimolar pool of libraries was sequenced with NexTSeq 500/550 High output kit v2 (75 cycles) (Illumina) with 10% PhiX.

### Bioinformatical analysis

The ribosome profiling analysis was made using the RiboDoc tool (v0.9.0) ^29^. The different main steps with corresponding programs, versions and command lines used in its analysis are described below. The reference genomes are Homo_sapiens.GRCh38.104 for Human (from the Ensembl database ^78^) and HIV1-pNL4-3 XCS for HIV.

The sequencing adapters were trimmed by cutadapt v4.3 ^79^ and the lengths of the RPFs was filter to keep reads from 25 to 35 nucleotides long as there expected length is around 30 nucleotides with the following parameters :

cutadapt -e 0.125 --max-n=1 -m 25 -M 35 -a ${adapter_sequence} -o ${output.fastq} ${input.fastq}

The removal of the rRNA reads was made by an alignment on the rRNA sequences by bowtie2 v2.5.1 ^80^ :

bowtie2 -x ${index.rRNA} -U ${input.fastq} --un-gz ${output.fastq}

The alignment on the genome was made with both hisat2 v2.2.1 ^81^ and bowtie2 v2.5.1 :

hisat2 -x ${hisat2_index.genome} --no-softclip -U ${input.fastq} --un-gz ${output.fastq} -S ${output.sam_hisat2}

bowtie2 -x ${bowtie2_index.genome} --end-to-end -U ${output.fastq} -S ${output.sam_bowtie2}

The selection of the reads uniquely mapped on the genome was made with samtools v1.14 ^82^ : samtools view -F 3844 -q 1 -h -o ${output.bam}

The counting of the reads corresponding to each transcript was done by htseqcount v2.0.2 ^83^ : htseq-count -f bam -t CDS -i Parent --additional-attr Name -m intersection-strict --nonunique fraction ${input.bam} ${input.gff} > ${output.txt}

The qualitative analysis was made on a transcriptome made from the genome with a selection of transcripts annotated as having a 5’UTR region only (for the human genome). This analysis and the determination of the P-site offset for each read length of every sample was made by the riboWaltz v1.2.0 package ^84^.

To study the reading frame of the Ribosome Protected Fragments (RPF), each read was represented by the coordinate corresponding to the first base of the associated ribosome’s P-site. To determine where the P-site is, a P-site offset has to be defined for every read length. This step was done with the riboWaltz program ^84^ which looks for the first base of each read beginning of the signal.

### Data sets and multiple alignments

Sequences were downloaded from the Los Alamos HIV Sequence Database (HIV-1 clades B and C) (https://www.hiv.lanl.gov/content/index). Sequences were filtered based on the presence of only one sequence per patient and excluding sequences carrying premature stop codons within CDS. Very similar sequences and incomplete sequences (more than 10 gaps/unknowns) were additionally discarded. We obtained 1609 and 412 sequences of HIV-1 clade B and clade C, respectively. The aa sequence conservation of the overlapping CDS and third frame were analyzed using Unipro UGENE toolkit. Multiple alignments were performed using ClustalO. Statistical analysis was performed with R (https://www.R-project.org/, version 4.2.2) using Rstudio IDE (2023.03.01.554). Comparison of population distributions were performed using Kruskal-Wallis test, Wilcoxon and Dunn tests. Plots were generated using ggplot2 package (v3.4.2). For each aa sequence conservation analysis of ARFs within HIV-1 clade B and C, the number of sequences used ranged from 168 to 1609 and from 47 to 411 sequences, respectively. Taking in consideration that in the UTR regions, the sequences may contain microdeletion and/or truncation, another pipeline was used for the ARF overlapping the UTRs. For each nucleotide sequence, the 6 phases where translated. Then, ARF aa sequences where align recursively to the 6 translated sequence. The best alignments were kept and alignments too far (+/- 150 nt) form the putative position where eliminated. Multiple alignments were performed using Clustal Omega.

### Immunopeptidome

#### Isolation of HLA ligands

5 x 10^8^ C8166 cells were infected with HIV_NL4-3ΔNef_ at a MOI of 0,05 and snap frozen 22 h post infection. HLA class I and HLA class II molecules were isolated using standard immunoaffinity purification ^85^, using the pan-HLA class I-specific W6/32, the pan-HLA class II-specific Tü-39, and the HLA-DR-specific L243 monoclonal antibodies (produced in-house) covalently linked to CNBr-activated Sepharose (Sigma-Aldrich). Cells were lysed in lysis buffer (CHAPS (Panreac AppliChem), complete protease inhibitor cocktail tablet (Roche) in PBS) for 1 h on a shaker at 4°C, sonicated, and centrifuged (45 min, 4000 rpm) and incubated again for 1 h. Lysates were cleared by sterile filtration (5 µm filter unit (Merck Millipore)) and cyclically passed through a column-based setup overnight at 4°C. Columns were washed with PBS (30 min) and ddH_2_O (1 h). Peptides were eluted by 0.2% trifluoroacetic acid (TFA), isolated by ultrafiltration (Amicon filter units (Merck Millipore)), lyophilized and desalted using ZipTip pipette tips with C18 resin (Merck).

#### Mass spectrometric data acquisition

For the mass spectrometric analysis peptides were loaded on a 75 µm x 2 cm PepMap nanotrap column (Thermo Fisher) at a flow rate of 4 µl/min for 10 min ^85^. Subsequent separation was performed by nanoflow high-performance liquid chromatography (RSLCnano, Thermo Fisher) using a 50 µm x 25 cm PepMap rapid separation column (Thermo Fisher, particle size of 2 µm) and a linear gradient ranging from 2.4 to 32.0% acetonitrile at a flow rate of 0.3 µl/min over the course of 90 min. Eluting peptides were analysed in technical replicates in an online-coupled Orbitrap Fusion Lumos mass spectrometer (Thermo Fisher) equipped with a nanoelectron spray ion source using a data dependent acquisition mode employing a top speed collisional-induced dissociation (CID, normalized collision energy 35%, HLA class I peptides) or higher-energy collisional dissociation (HCD, normalized collision energy 30%, HLA class II peptides) fragmentation method. MS1 and MS2 spectra were detected in the Orbitrap with a resolution of 120,000 and 30,000, respectively. The maximum injection time was set to 50 ms and 150 ms for MS1 and MS2, respectively. The dynamic exclusion was set to 7 s and 10 s for HLA class I and HLA class II, respectively. Mass range for HLA class I peptide analysis was set to 400 - 650 m/z with charge states 2+ and 3+ selected for fragmentation. For HLA class II peptide analysis mass range was limited to 400 1,000 m/z with charge states 2+ to 5+ selected for fragmentation.

#### Data processing

For data processing, the SEQUEST HT search engine (University of Washington ^86^) was used to search the human proteome as comprised in the Swiss-Prot database (20,394 reviewed protein sequences, December 4th 2020) supplemented with the six-frame translated genome of HIV_NL4-3_, taking into account all potential peptides ≥ 8 aa encompassed between 2 stop codons without enzymatic restriction. Precursor mass tolerance was set to 5 ppm, and fragment mass tolerance to 0.02 Da. Oxidized methionine was allowed as a dynamic modification. The peptide spectrum matches (PSM) false discovery rate (FDR) was estimated using the Percolator algorithm ^87^ and limited to 5% for HLA class I and 1% for HLA class II. Peptide lengths were limited to 8 - 12 aa for HLA class I and to 8 - 25 aa for HLA class II. Protein inference was disabled, allowing for multiple protein annotations of peptides. HLA class I annotation was performed using NetMHCpan 4.1 ^88^ and SYFPEITHI ^89^ annotating peptides with percentile rank below 2% and ≥ 60% of the maximal score, respectively.

#### Spectrum validation

Spectrum validation of the experimentally eluted peptide was performed by computing the similarity of the spectra with the corresponding isotope-labeled synthetic peptides measured in a complex matrix. The spectral correlation was calculated between the MS/MS spectra of the eluted and the synthetic peptide ^90^.

### PLWH and Samples

EC (n=4) were recruited from the CO21 CODEX cohort implemented by the ANRS|MIE (Agence Nationale de recherches sur le SIDA, les hépatites virales|Maladies infectieuses émergentes). PBMC were cryopreserved at enrollment. ECs were defined as HIV-infected individuals maintaining viral loads (VL) under 400 copies of HIV RNA/mL without treament for more than 5 years. HIV-infected efficiently treated PLWH (ART) (n=3) were recruited at AP-HP Bicêtre Hospital (Kremlin Bicêtre, France). They were treated for at least 1 year (mean of 10 years) and have an undetectable viral load using standard assays. HIV-negative individuals (n=3) were anonymous blood donors from EFS (Établissement Français du sang). A detailed description of the donors is provided in Table 4, including the median and interquartile range for age (at the time of the study), CD4 T cell count and RNA load for each group. HIV-RNA loads were measured on site with different real-time PCR-based assays; depending on the date of enrollment in the cohort and the assay routinely used on each site, the VL detection limit varied from 50 to 10 copies/mL.

### Ethic statement

All the subjects provided their written informed consent to participate in the study. The CO21 CODEX cohort and this sub-study were funded and sponsored by ANRS|MIE and approved by the Ile de France VII Ethics Committee. The study was conducted according to the principles expressed in the Declaration of Helsinki.

### Peptides

ARF-derived peptides were chosen based on the aa conservation among the clade B isolates and the predictions to bind frequent HLA alleles. The binding affinity of ARF-derived peptides was evaluated using the NetMHCPan-4.1 and NetMHCIIPan-4.1 ^88^. For the longest ARF sequences, some overlapping peptides were also designed. The synthetic peptides were then randomly distributed in 9 different pools to obtain 11 or less peptides per pool to limit peptide competition for HLA-binding (Table 3). HIV-classical epitopes were chosen based on the literature ^37^ (Supplementary Table 1). ARF-derived peptides and HIV-classical peptides were synthetized by Vivitide company and in-house at the Department of Immunology, University of Tübingen, Germany. The in-house production was performed with the peptide synthesizer Liberty Blue (CEM) using the 9 fluorenylmethyl-oxycarbonyl/tert-butyl strategy ^91^. Non-HIV common peptides (from HCMV, EBV and influenza virus) were obtained from Mabtech (PepPool: CEF (CD8), human, 3616-1).

### 12-day in vitro T cell amplification prior ELISPOT assay

PBMCs were thawed and rested 2-3h in IMDM (Gibco/ThermoFisher Scientific) containing 5% human AB serum (SAB, Institut Jacques Boy), supplemented with recombinant human IL-2 (rhIL2, 10 U/ml, Miltenyi) and Dnase I (1 U/ml, New England Biolabs). Cells were washed and 5-9 x 10^6^ PBMCs were then seeded/well in a 24-well plate in IMDM supplemented with 10 % SAB, Pen/Strep, nonessential aa and sodium pyruvate (all from Gibco/ThermoFisher Scientific). Nevirapine (NVP, 1.2 µM, HIV reagent program) was added to inhibit potential viral replication and poly I:C (2 µg/ml, InvivoGen) to facilitate the presentation of long peptides by antigen presenting cells ^92^. PBMCs were loaded with peptide pools (each peptide at 10 µg/ml), except for the CEF pool (Mabtech) used at 5 µg/ml, and cultured overnight at 37°C, 5% CO2. HIV-classical peptides and/or CEF peptide pools were used as positive control for the expansion of HIV-specific and non-HIV common anti-virus-specific T cells, respectively. On day 1 and 3, T cell media were then complemented with rhIL2 (10 U/ml) and rhIL-7 (20 ng/ml, Miltenyi) with NVP (1.2 µM), respectively, and maintained though out the culture. On days 3, 5, 7, and 9, when the cell layer was > 70% confluent, cells were split in two wells before the addition of rhIL-2, rhIL-7 and NVP. On day 7, cells were harvested, counted and a fraction submitted to IFN-γ ELISPOT assay using the ARF-derived peptide pools and positive controls: HIV classical peptide and/or CEF peptide pools. The remaining cells were maintained in culture and submitted, on day 12, to IFN-γ ELISPOT assay using individual ARF-derived peptides and the controls.

### Enzyme-linked immunospot assay - cultured IFN-γ ELISPOT

1-3 x 10^5^ cells/well were seeded in an ELISPOT plates (MSIPN4550, Millipore) in IMDM supplemented with 10 % SAB, Pen/Strep, nonessential aa, sodium pyruvate 10 %, and HEPES 10mM (Gibco/ThermoFisher Scientific), loaded with either 50, 10 or 2 µg/ml of ARF-derived peptides, HIV classical peptides and CEF peptides, respectively and incubated at 37°C for 16h. ELISPOT plates were pre-coated with anti-IFNγ primary antibody and after cell-incubation IFNγ revealed using anti-IFNγ secondary antibody conjugated with biotin (both from Mabtech) as described ^24^. The ELISPOT analysis was performed in technical triplicates or duplicates. Spots were counted with the AID ELISPOT Reader according standard protocols. Responses were considered positive when IFNγ production was superior to 20 spots/10^6^ PBMCs and at least twofold higher than background (cells loaded with the DMSO containing medium used to solubilize the peptides).

### Intracellular Cytokine Staining (ICS) assay

Cells were either cultured as described for the 12-day in vitro T cell amplification or treated directly after thawing (for the *ex vivo* assays). On day 12, cells were harvested, washed and counted. 2-10 x 10^5^ PBMCs were then seeded/well of a 96-well U-bottom plate in IMDM supplemented with 10 % SAB, Pen/Strep, nonessential aa, sodium pyruvate 10% and HEPES 10mM. PBMCs were immediately loaded with individual ARF-derived peptides (50 µg/ml), individual HIV-classical peptides (10 µg/ml) or CEF peptide pool (2ug/ml). Additionally, the CD107a-BV786 antibody (BD Biosciences) was included in each condition for degranulation detection. For the *ex vivo* assessment of T cell activation, 2-4 x 10^6^ PBMCs were seeded per well of a 96-well U-bottom plate using the same IMDM supplemented medium. The same peptide loading and CD107a-BV786 antibody addition steps were performed. After 1h at 37°C, brefeldin A was added (5 µg/ml) and the incubation carried out for additional 5h. PBMCs were washed in PBS and incubated with LVD and human Fc Block (BD Biosciences). Cells were then stained 30 min on ice with a mix of cell surface antibodies: CD3-PE, CD8-SuperBright600 (Invitrogen) and CD4-AF750 (Beckman Coulter). Next, cells were washed, fixed with PFA 4% and permeabilized (PBS + BSA 0.5%, saponin 0,05%). Cytokine production were detected by intracellular staining using IFN-γ-PerCP-Cy-5.5, MIP1β-FITC, TNF-PE-Cy7 and IL2-APC antibodies (all from BD Biosciences).

### Flow cytometry and data analysis

Sample acquisition and analysis were performed using Cytoflex with CytExpert software (Beckman Coulter). The compensation matrix was calculated automatically by the CytExpert software after measuring negative and single-color controls. Sample analysis was performed using FlowJo v10.8 Software (BD Life Sciences). After elimination of doublets, lymphocytes were gated on side scatter area versus forward scatter area pseudo-color dot plot and dead cells were removed according to LVD staining. CD4^+^CD3^+^ events were gated versus individual gates made for the detection of MIP1β, IFNγ, IL2, CD107a and TNF secretion. All gates were set based on the unstimulated control (cells loaded with the DMSO containing medium used to solubilized the peptides). Individual cytokine secretions were then combined together using Boolean gating to create a full array of possible combinations of response patterns from the CD4^+^CD3^+^-cell subsets. The same procedure was used to CD8^+^CD3^+^ events. Positive responses were reported after background subtraction. Samples were considered positive when the % of activated cells was at least 2 times higher than negative controls.

### Statistical analysis

Statistical significances (p-values) were calculated using Prism Software (GraphPad). Statistical tests used for each individual figure are indicated in the figure legends.

## Code Availability

The code package for this study is available and fully described in the above section “Bioinformatical analysis”.

## Data Availability

The mass spectrometry immunopeptidomics data have been deposited to the ProteomeXchange Consortium (http://proteomecentral.proteomexchange.org) via the PRIDE (Perez-Riverol *et al*, 2019) partner repository with the dataset identifier:

Project accession: PXD043984

Username

Password: PL8zE01i

The Riboseq data have been deposited to the public functional genomics data repository GEO: https://www.ncbi.nlm.nih.gov/geo/query/acc.cgi?acc=GSE239818

Reviewer token: orqbsuwsznwdnmz

## Acknowledgements

This work was granted by the Agence Nationale de Recherche sur le SIDA (ANRS-Maladies infectieuses émergentes) and Sidaction for fundings. L.B, A.B and E.L were supported by ANRS. L.B. and M.P. were supported by Sidaction. G.B. was supported by ANR-20-IDEES-0002. We thank Remi Villette for his expertise using R, Bernard Maillere and his team for the access to the ELISPOT reader, Anne Lopes, Paul Roginski, for discussions, Frederic Suba and Clemence Richetta for access to the L3 facility of ENS-Paris-Saclay. We thank all participants of the ANRS CODEX cohort and the NIH AIDS Research and Reference Reagent Program for providing drugs. This work was supported by the Deutsche Forschungsgemeinschaft (DFG, German Research Foundation, WA 4608/1-2, J.S.W.), the Deutsche Forschungsgemeinschaft under Germany’s Excellence Strategy (EXC2180 390900677, J.S.W. and H.-G.R.), the German Cancer Consortium (DKTK, H.-G.R. and J.S.W.), the Ernst Jung Prize for Medicine (H.-G.R.), the Landesforschungspreis of Baden-Württemberg (H.-G.R.), the Wilhelm Sander Stiftung (2016.177.3, J.S.W.), the Deutsche Krebshilfe (German Cancer Aid, 70114948, J.S.W.), and the Fortüne Program of the University of Tübingen (2451-0-0, J.S.W.). The collaboration between Tübingen University and I2BC was supported by the French-German Partnership Hubert Curien Procope 2021 program.

## Author contributions

A.M. conceived and designed the project together with O.N. for the Riboseq experiments. Design and performed the experiments: L.B., A.N., B.C.R., I.H., S.G., D.D., L.G., A.B., Y.V. Analysed the data: H.A., P.F., L.B., G.B., A.N.,, S.D., B.C.R., I.H., S.G., Y.V., J.V, M.P., S.G.-D., J.W. O.N. and A.M. Contributed reagents/materials/analysis tools: H.A., P.F., G.B., I.H., A.G., M.C.Z., C.G., M.A., L.H., E.R., E.L., R.J-M., S.C., A.E., A.S., B.A., O.L., H.-G.R., J.W., O.N. and A.M.. Wrote the paper: L.B. and A.M. with contributions from H.A., A.N. and O.N, and all authors.

## Materials and Correspondence

Arnaud Moris

## SUPPLEMENTARY FIGURE & TABLE LEGENDS

**Supplementary Table 1: List of synthesized classical HIV peptides (included in the HIV pool) derived from viral CDSs** and used as positive response for HIV-specific T cell responses in IFNγ-ELISPOT assays.

**Supplementary Table 2: List of ARF-derived peptides tested in ICS.** Peptides were named according to the ARF from which there are translated from (left column). The short name of the peptides is the peptide name used in the Figures. Pools where the peptides were included, peptide sequences and lengths are indicated.

**Supplementary Table 3: List of classical HIV peptides used in ICS.** These HIV CDS-derived peptides were previously described as immunodominant CD8^+^ and CD4^+^ T cell epitopes (https://www.hiv.lanl.gov/content/immunology). Additional CD4^+^ T cell epitopes used as positive controls in IFNγ-ELISPOT assays (included in the HIV pool) are listed. The sequences, CDSs, HLA-restrictions and names of each peptide are indicated.

**Supplementary Fig. 1-related to Fig. 1. (a) Outline of ribosome profiling protocol.** Twenty hours post-infection a fraction of HIV_NL4-3_-infected CD4^+^ SupT1 cells was collected. The viability and percentage of HIV-Gag^+^ and CD4^+^ cells were assessed using flow cytometry. Four hours later (24h post-infection) infected cells were treated with CHX and lysed. Polysomes were then extracted, treated with RNase, ribosomal RNAs (rRNA) depleted and ribosomes protected fragments (RPFs) isolated and sequenced. **(b) Analysis of the infection rate of HIV_NL4-3_-infected CD4^+^ SupT1 cells.** The gating strategy and the percentage of positive cells in each quadrant is shown. Three biological replicates were used. Infected cells were labeled with a viable dye and stained with antibodies specific to CD4 and HIV-Gag. Left panels: the percentages of viable cells. Middle and right panels: percentage of infected CD4^-^HIV-Gag^+^ and CD4^+^HIV-Gag^+^ cells within HIV_NL4-3_-infected and mock treated CD4^+^ SupT1 cells, respectively. The same negative control was used for replicate 1 and 2 since they were performed side by side. **(c) Trinucleotide periodicity on cellular transcripts/metagene of 29 nt.** Data were analysed using an inhouse bioinformatics pipeline. RFPs were aligned to annotated start codons of human CDSs to analyse the periodicity of the reads. This allows us to calculate an offset between the 5’ extremity of the reads and the first nucleotide at the ribosomal P-site position (+12) for the 28- and 29-mer. The plot corresponds to position of the +12 nucleotide of each ribosome footprints around the start codon (-14/+29). Each reading frame is visualised by a different colour.

**Supplementary Fig. 2-related to Fig. 1**.: **Distribution of ribosome protected fragments (RPFs) across HIV-1 genome obtained from HIV_NL4-3_-infected SupT1 cells.** The number of RPFs (Nb of reads) are represented according to the position in HIV_NL4-3_ genome. For clarity, the numbers of RPFs aligned on the three reading frames are totally presented in (a) and truncated above 5000 reads in (b). In (b), the data are also shown with RPF occupancy histograms separated by reading frame. As in Fig. 1, RPFs translated in frames 0, +1, and +2 are represented in green, red, and blue, respectively. HIV genome is represented below; CDSs are indicated and colored accordingly to their reading frames from the first position of the genome. The positioning of ARFs (purple lines) are indicated. **(c) ARF selection criteria.** Sequenced RPFs were analysed using an inhouse bioinformatics pipeline and RPFs were aligned on HIV_NL4-3_ genome according to their reading frame. CDSs and ARFs translated in frame 0, +1, and +2 are represented in green, red, and blue, respectively. The ARF inclusion criteria were: 1) not corresponding to CDS sequences; 2) amino-acid length ≥ 10; 3) translated between 2 stops codons and 4) reads per codon ≥ 8. Green rectangles indicate the canonical *gag* CDS. Left panel: the blue rectangle indicates the ARF-GAG16 that passed the inclusion criteria. right panel: the untagged red rectangle shows a potential ARF that failed the inclusion criteria.

**Supplementary Fig. 3-related to Fig. 1: Distribution of ribosome protected fragments (RPFs) in a zoom plot of the HIV genomic region containing the major splice donor site (SD1) obtained from HIV_NL4-3_-infected SupT1 cells (see Fig. 1).** The number of RPFs (Nb of reads) are represented according to the position in HIV_NL4-3_ genome. For clarity, the numbers of RPFs are totally presented and truncated above 5000 reads in the top panel and lower panel, respectively. As in Fig. 1, RPFs translated in frames 0, +1, and +2 are represented in green, red, and blue, respectively. For each frame, the HIV genome is represented below. At the bottom, ARF identified in this region are indicated with rectangles and colored accordingly to their reading frames. Colored crosses and triangles indicate stop codons and putative start codons. The position of ARF-derived peptides that induce potent T cell responses in PBMCs of PLWH (see Fig. 3) are also shown for each ARF using the same color coding. The position of SD1 and its hairpin loop is also indicated.

**Supplementary Fig. 4-related to Table 2: Identification of ARF start codons using translation inhibitors lactimidomycin (LTM) and puromycin (PMY) followed by Riboseq.** In a second set of Riboseq experiments, LTM+PMY-treatments were used to define the start codons of the ARFs. **Trinucleotide periodicity on cellular transcripts/metagene of 30 mers.** Data were analysed using an inhouse bioinformatics pipeline. RFPs were aligned to annotated start and stop codons of human CDSs to analyse the periodicity of the reads and the accumulation of reads at the initiation and termination sites. Each reading frame is visualised by a different colour. The three plots correspond to the metagene of the three biological replicates.

**Supplementary Fig. 5-related to Fig. 2: Features of ARF amino-acid (aa) conservation among HIV-1 clade B and C**. **(a) Percentage of aa conservation of each identified ARFs among HIV-1 clade B, depending on the reading frame.** ARFs are represented in relation to the reading frame of the CDS (x-axis). Kruskal-Wallis test was performed, and the p-value is reported on the graphic. **(b) Percentage of aa conservation of each identified ARFs depending on the overlapping CDS.** Kruskal-Wallis test was conducted followed by a post-hoc Dunn test (Holm correction), and the p-values (<0.001 and <0.0001) are reported on the graphic. **(c) Comparison of ARFs aa sequence conservation between clade B and C.** A pairwise Wilcoxon test was applied and p-values are shown on the graph. **(d) Correlation between ARF aa sequence conservation among clade B (blue) and C (red) and ARFs relative expression (expressed as reads/nucleotide). (e) CDS aa sequences are slightly more conserved than the aa sequences of ARF and of the third frame.** The percentage of aa conservation of the overlapping CDSs, ARFs, and third frames are shown. Kruskal-Wallis test followed by a pairwise Wilcoxon test was achieved, and p-values are shown on the graph.

**Supplementary Fig. 6-related to Fig. 3: Detection of ARF-derived peptide (ARFP)-specific T cell responses using cultured IFNγ-ELISPOT assay. (a) Experimental procedure of the 12-day T cell amplification.** PBMCs from HIV-infected individuals were seeded in wells of 24-well plates and loaded with the different peptide pools in the presence of the indicated cytokines and polyI:C to allow a better presentation of elongated peptides and Nevirapine to block potential viral replication. Seven days later, a fraction of the cells was counted and loaded with or without the peptide pool used for the initial culture and T cell reactivity was monitored using an IFNγ-ELISPOT assay. The remaining cells were then allowed to expand further by renewing the cytokine cocktail and tested on day 12, after the initial culture, for their capacity to react to individual peptides of the pools tested positive at day 7 using IFNγ-ELISPOT assay. **(b) Day 7 IFNγ-ELISPOT responses against all peptide pools for every HIV-infected individual** presented as SFU/10^6^ PBMCs (y-axis). A response was considered positive when the number of rough IFNγ^+^ spots per well was >20 and 2 times higher than the negative control (POOL (-) white bar, cells loaded with DMSO-containing medium). Positive responses are indicated by colored bars depending on the overlapping genomic region. HIV-CDS-derived peptides pools were used as a positive control. SAT: saturated well for which the IFNγ signal was so high that it could not be quantify.

**Supplementary Fig. 7-related to Fig. 3: Detection of T cell responses specific to unique ARF-derived peptides (ARFP) in the PBMCs of HIV-infected individuals.** At day 12 after the initial culture, the capacity of cultured PBMCs from HIV-infected donors to react to individual peptides of the pools, tested positive at day 7, was assessed using IFNγ-ELISPOT assay. The results are presented as SFU/10^6^ PBMCs. A response was considered positive when the number of spots was >20 and 2 times higher than the negative control (POOL (-) white bar, cells loaded with DMSO-containing medium). Shaded lines represent the threshold of positivity. HIV-classical peptide pool and CEF peptides were used as positive controls. **(a), (b), (c), (d), (e), (f) and (g) results from the cultured IFNγ-ELISPOT assay of cells from ART-1, ART-2, ART-3, EC-1, EC-2, EC-3 and EC-4 HIV-infected individuals, respectively.** Only deconvolutions for which a peptide pool elicited a significant T response are shown. Pictures of the original rough data from the ELISPOT plates are presented below or on the sides of the corresponding graphic (ARFP and controls were tested in duplicates or triplicates). SAT: saturated well for which the IFNγ signal was so high that it could not be quantify.

**Supplementary Fig. 8-related to Fig. 3**: **Analysis of ARF-derived peptide (ARFP)-specific T cells responses. (a) ARF-derived peptide stimulation levels are significantly higher than negative controls.** Responses to individual ARFP were plotted side-by-side to their respective negative controls. Saturated responses were assigned an arbitrary score of 1250 SFU/10⁶ PBMCs. Wilcoxon matched-pairs signed rank test was applied. p-value: (**\*\*\*\***<0.0001). **(b) Comparison of the magnitude of T cell responses between ART and EC.** Responses to unique ARFP were plotted for each PLWH individual as SFU/10^6^ PBMCs. Negative controls were subtracted and saturated responses attributed on arbitrary score of 1250 SFU/10^6^ PBMCs. Kruskal-Wallis test with Dunn’s correction was applied. No significant differences were observed. **(c) uORF-derived peptide stimulation levels are significantly higher than negative controls.** As in (a) focusing on uORF-derived peptide. p-value: (**\*\***<0.002). **(d) Comparison of the magnitude of T cell responses induced by ARFP encoded from different HIV genomic regions.** Responses to individual ARFP were plotted and grouped according to the overlapping HIV CDS or UTR. Negative controls (POOL -) were subtracted and saturated responses were assigned an arbitrary score of 1250 SFU/10⁶ PBMCs. One-way ANOVA test was applied. No significant differences were observed.

**Supplementary Fig. 9-related to Fig. 3: Detection of T cell responses to ARF-derived peptides in the PBMCs of non-infected individuals.** PBMCs from non-infected individuals (EFS 0685, EFS 1619, and EFS 5174) were seeded and loaded with the different peptide pools as in Supplementary Fig. 4. T-cell reactivity to the peptide pools used for the initial culture (a) and to individual peptides of the pools (b) was monitored using an IFNγ-ELISPOT, at day 7 and day 12, respectively. IFNγ-ELISPOT are presented as SFU/10^6^ PBMCs (y-axis). A response was considered positive when the number of spots was >20 and 2 times higher than the negative control (POOL (-) white bar, cells loaded with DMSO-containing medium). (a) CEF peptide pool was used as a positive control. SAT: saturated well for which the IFNγ signal was so high that it could not be quantify. (a) Positive responses are indicated by colored bars depending on the overlapping genomic region. (b) Shaded lines represent the threshold of positivity. Only deconvolutions for which a peptide pool elicited a significant T response are shown. SAT: saturated well for which the IFNγ signal was so high that it could not be quantify.

**Supplementary Fig. 10-related to Fig. 4: Analysis of the polyfunctional profiles of ARFP-, CDS- and CEF-specific T cell responses in PBMCs of HIV-infected individuals. Percentages of ARF-(blue dots) or CDS-(grey dots) specific CD4^+^CD3^+^ (a) and CD8^+^CD3^+^ (b) T cell responses secreting the indicated effector molecules. (b)** CEF-specific CD8^+^ T cell responses are also presented (red dots). Negative controls were subtracted. Mann-Whitney U test and p values are shown. **(c, d) Heatmaps summarizing the frequency of simultaneously secreted cytokine/chemokine (x-axis, from 1 to 5) upon activation of CD4^+^CD3^+^ (c) and CD8^+^CD3^+^ (d) T cells using individual ARFP-(left panels), CDS- or CEF-derived (right panels) peptides for each HIV-infected donor.** The frequency of secreting T cells is represented by the filling gradient. Only ARFP that stimulated T cell responses are shown, **(e) Effector molecules secretion profiles of CD4^+^CD3^+^ (top panel) and CD8^+^CD3^+^ (bottom panel) T cell responses against ARF-(top row) and CDS-derived (bottom row) peptides.** The 32 possible patterns of effector molecules secretions are indicated by the grey line below the heatmap and defined in a listed format on the right. (+) and (-) indicated produced and not produced molecules respectively.

**Supplementary Fig. 11-related to Fig. 4: ARF-derived peptides (ARFP) elicit specific T cell responses *ex vivo* in PBMCs of HIV-infected individuals. ARFP-specific T cell responses from EC-1 (a), and EC-3 (b) donors.** Intracellular cytokine staining was performed for MIP-1β, IFNγ, IL-2, CD107a, and TNF after PBMC stimulation with individual ARFP. Positive controls included HIV-classical peptides and non-HIV common viral peptides. Table on the left provide an overview of peptide stimulation and associated T-cell responses. Cells were gated on CD4^+^CD3^+^ or CD8^+^CD3^+^ T cell populations, and individual gates for each cytokine/chemokine were determined based on negative controls (cells loaded with DMSO-containing medium). CD4^+^/CD8^+^CD3^+^ cells were pre-gated on living cells, excluding doublets, and backup gating was applied to limit non-specific signals. The upper pie charts summarize the % of activated (white) and non-activated (blue) T cells following peptide stimulation. The lower pie charts represent the polyfunctionality of activated T cells. Boolean analysis was conducted to determine whether T cells secreted 1, 2, 3, 4, or 5 cytokines simultaneously (red, green, purple, blue, and orange, respectively). Numbers indicate the percentage of activated cells secreting one to five cytokines simultaneously. **(b)** on the right, the gating and responses of YSL-I-specific CD4+ T cell responses are shown. Gates for each cytokine/chemokine were set based on the negative control (cells loaded with DMSO containing medium, NS: Not stimulated) NS_YSL-I. CD4^+^CD8^-^ T cell populations were pre-gated on CD4^+^CD3^+^ cells, lived cells and doublets were excluded.

**Supplementary Fig. 12-related to Fig. 5: Immunopeptidome analysis of HIV-infected cells. Defining the immunopeptidome of HIV-derived HLA class II associated peptides in HIV_NL4-3ΔNef_-infected C8166 cells by mass spectrometery. (a)** Briefly, C8166 cells were infected or not (mock) with HIV_NL**4-3ΔNef**_ at 0.05 MOI. 24 hours post-infection, a fraction of cells was stained with an HIVGag-p24 antibody to estimate the fraction of HIV-Gag^+^ infected cells. Mock infected cells were used to delineate the HIV-Gag^+^ population (21% of HIV-Gag^+^ cells). **(b)** Cells were then lysed, HLA class-II complexes immunoprecipitated using Tü39 antibody and HLA-ligands sequenced using LC-MS/MS. **(c) Table summarizing HIV-derived HLA class II-presented peptides isolated from HIV_NL4-3ΔNef_-infected C8166 cells.** All peptides are derived from classical HIV CDSs. **(d) Validation of the experimentally eluted ARF-derived peptide ILINGQYSL.** Comparison of fragment spectra (m/z on the x-axis) of the HLA-restricted peptide eluted from the HIV_NL4-3_-infected primary CD4^+^ T cells (identification) to its corresponding synthetic peptide (validation, mirrored on the x-axis) with the calculated spectral correlation coefficient (R2). Identified b- and y-ions are marked in red and blue, respectively. (e) **Characteristics of YSL-ARF.** The frame and the positions in HIV_NL4-3_ genome of the beginning of YSL-ARF (stop-to-stop), of the start (identified in the Riboseq+LTM+PMY experiments) and stop codons, as well as the nucleotide and aa sequences and size are indicated. The ILINGQYSL (YSL-I) and the LHSFGWVMNSILING (YSL-II) aa sequences are highlighted in red or underlined, respectively.

