## Supplementary Fig_1_to_13 & Table 1_to_3 for "Unveiling novel conserved HIV-1 open reading frames encoding T cell antigens using ribosome profiling"

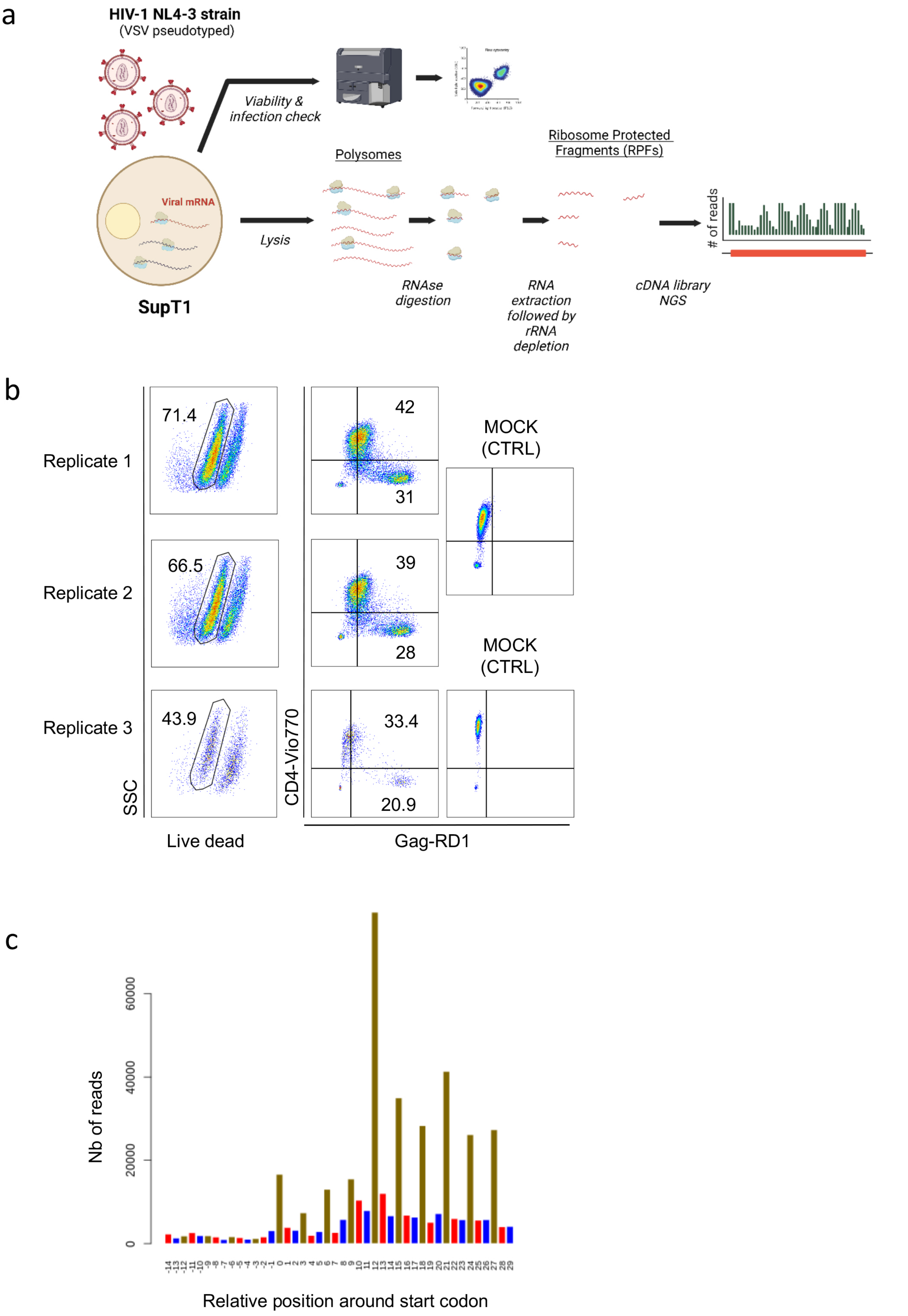


Supplementary Fig. 1 related to Fig.1

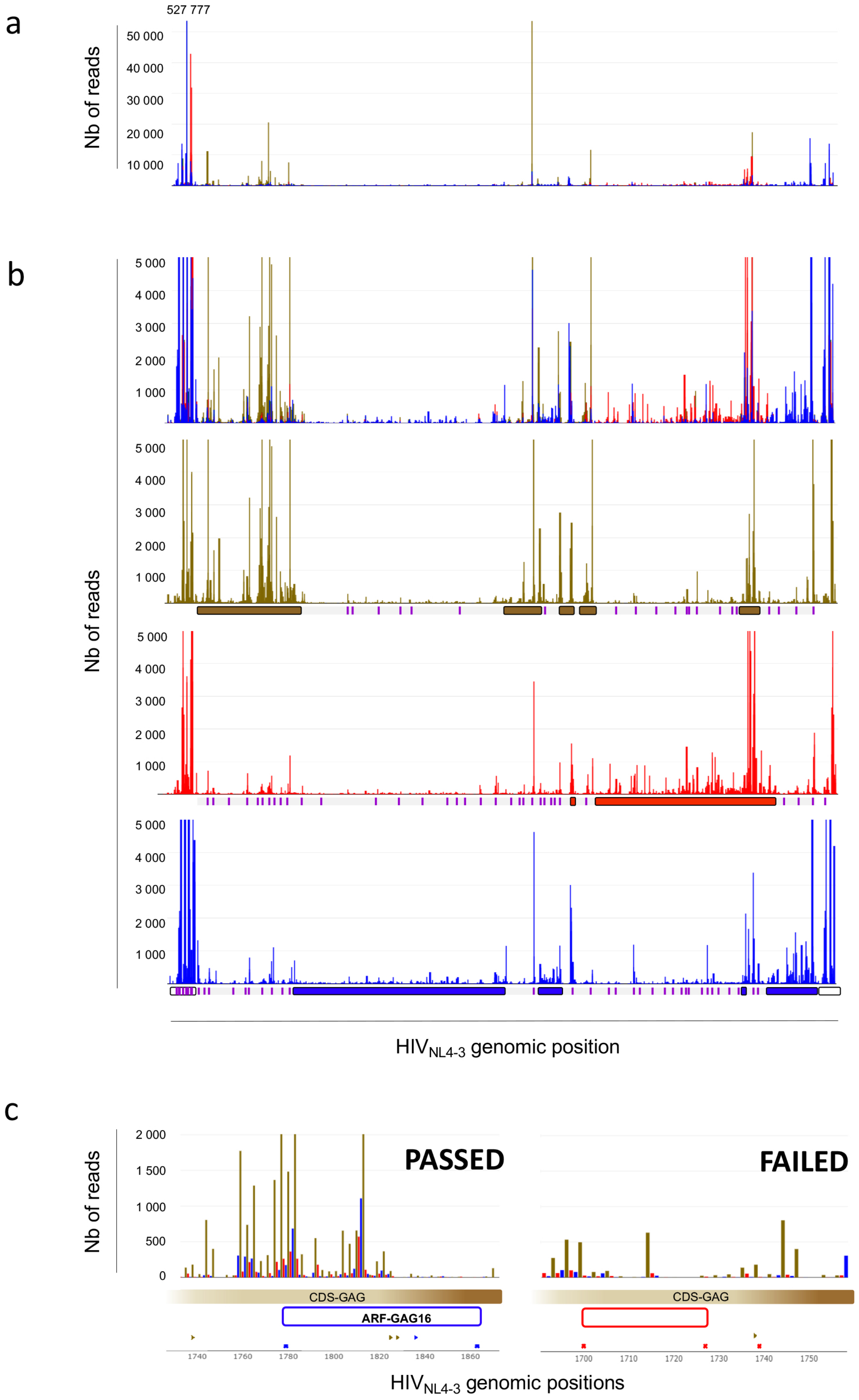


Supplementary Fig. 2 related to Fig. 1

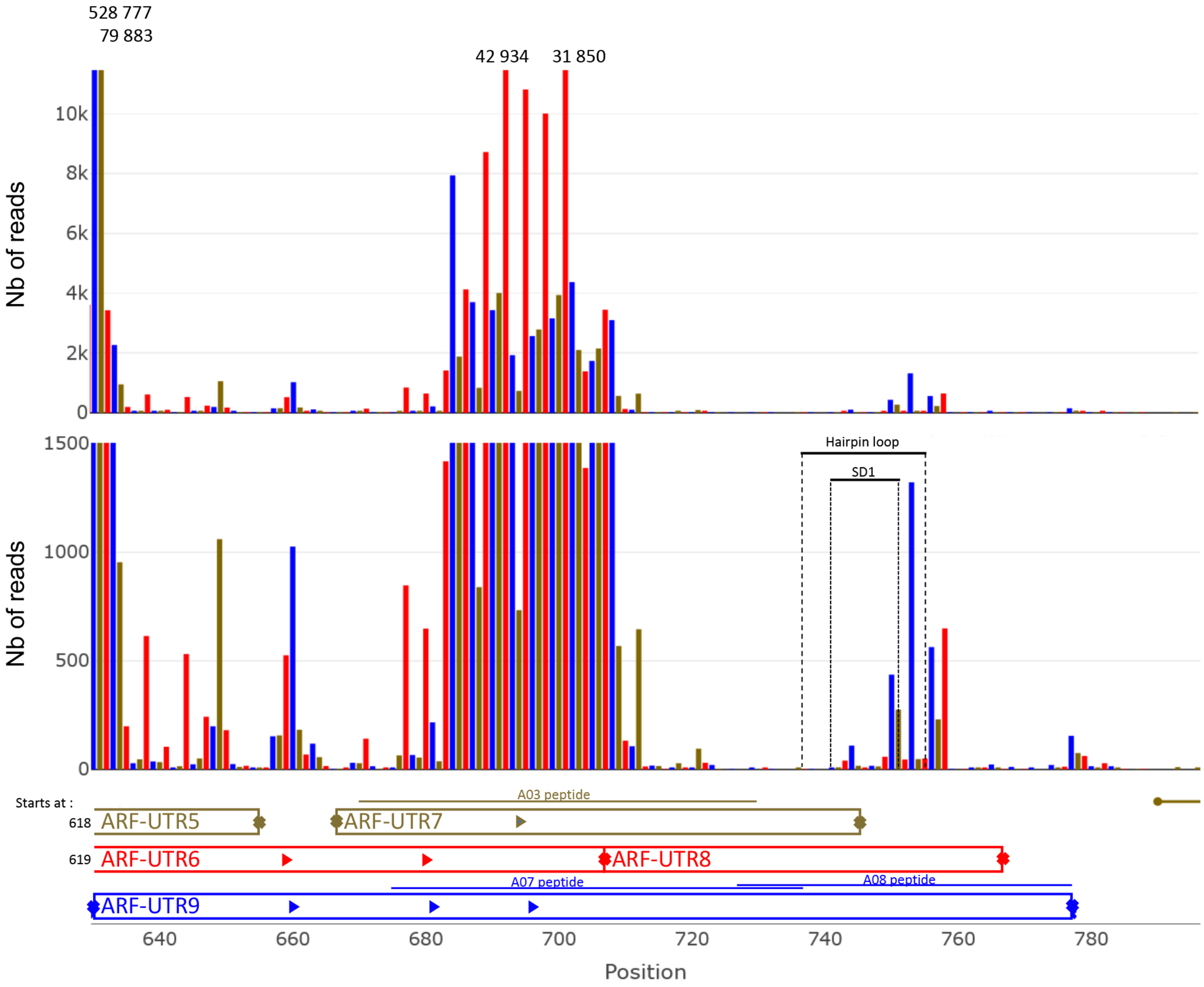


Supplementary Fig. 3 related to Fig. 1

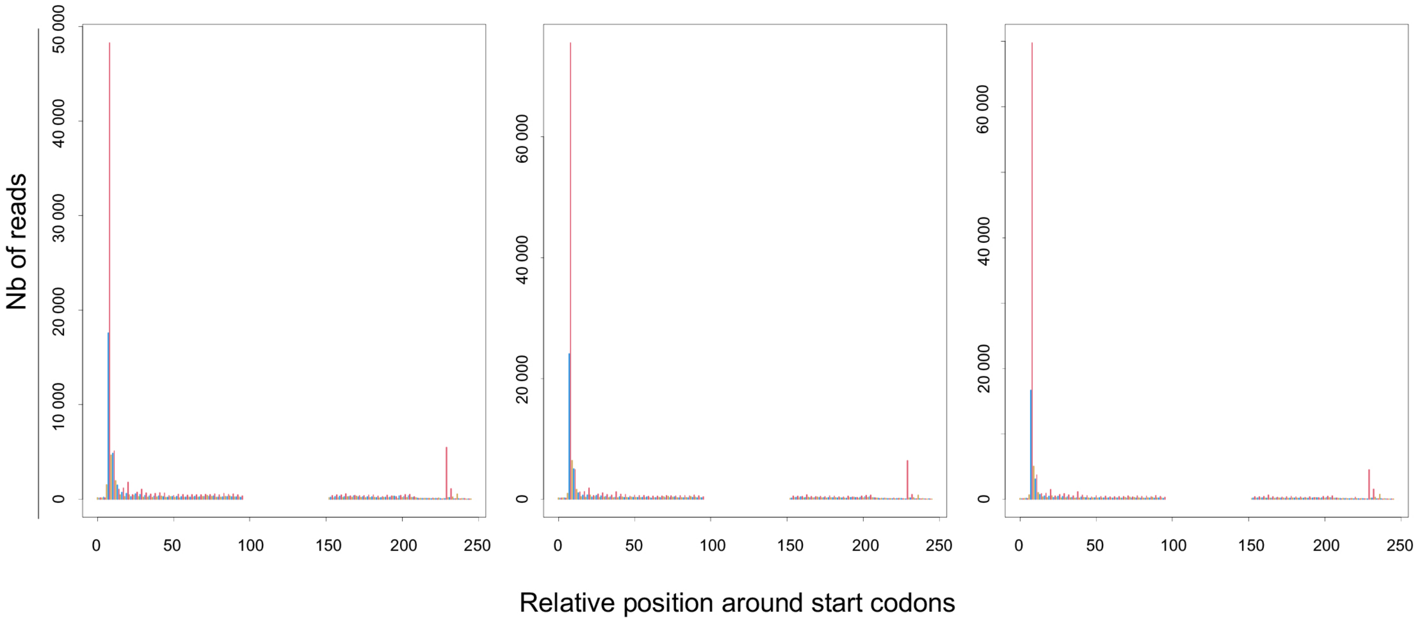


Supplementary Fig. 4 related to Table 2

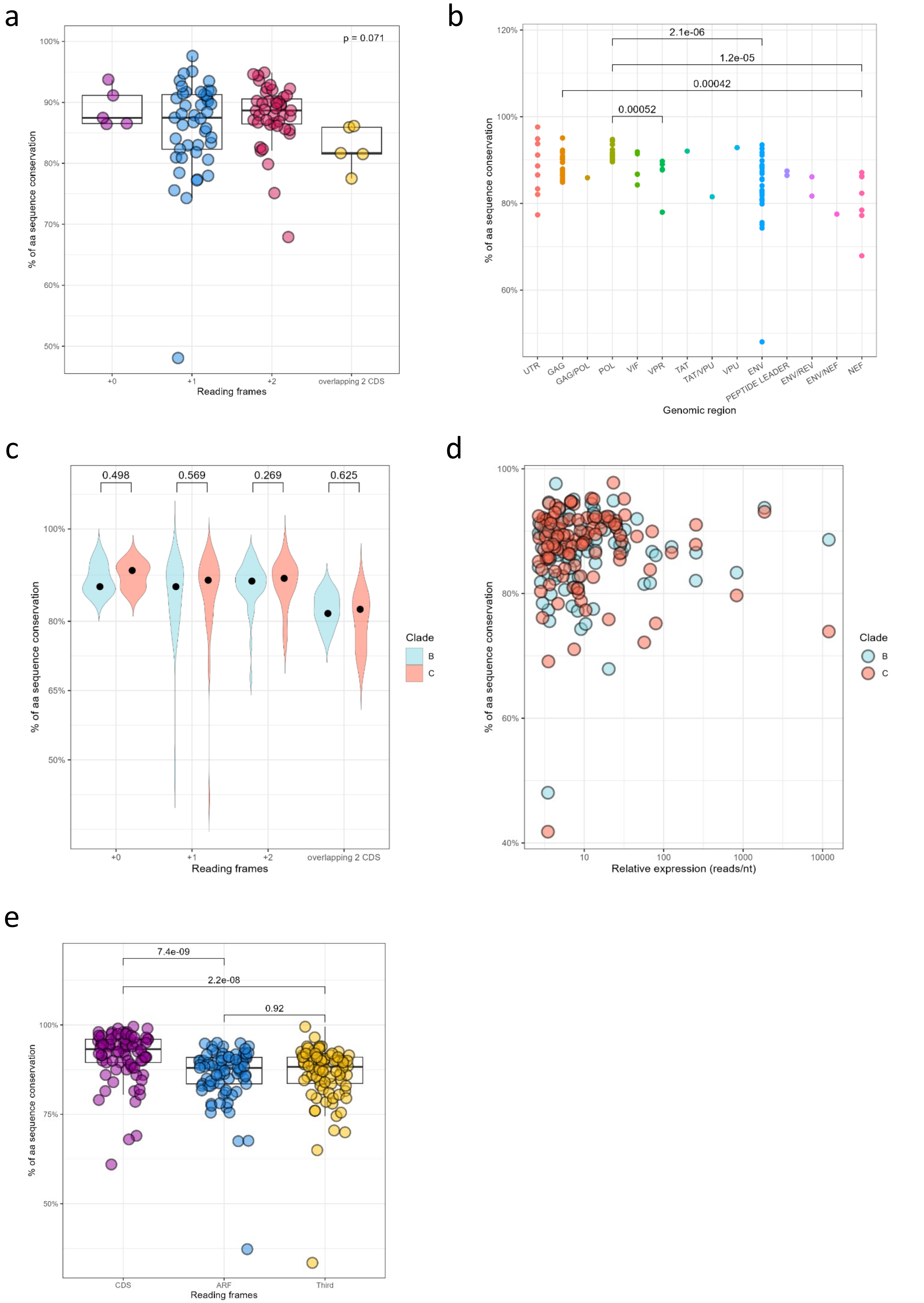


Supplementary Fig. 5 related to Fig.2

a

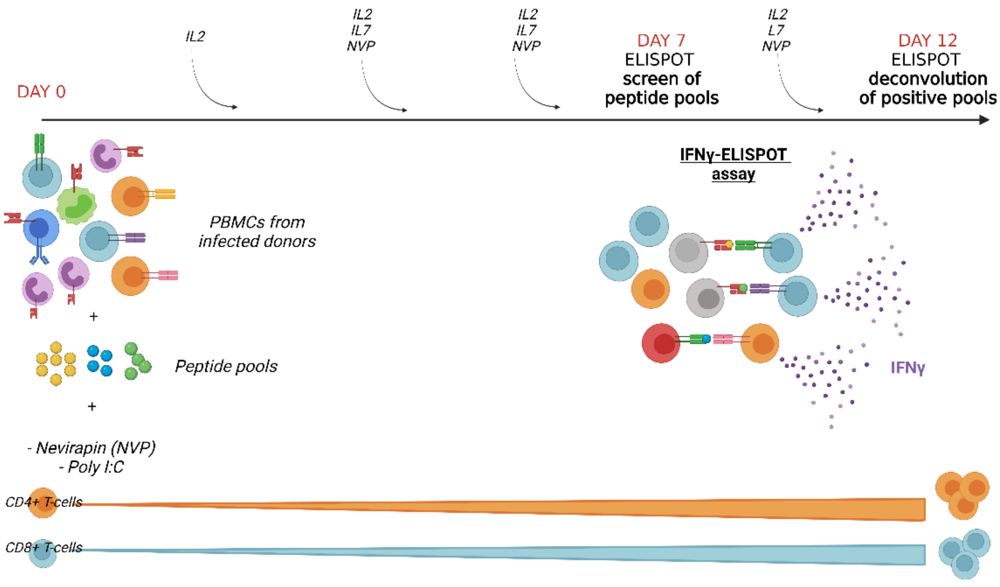


b

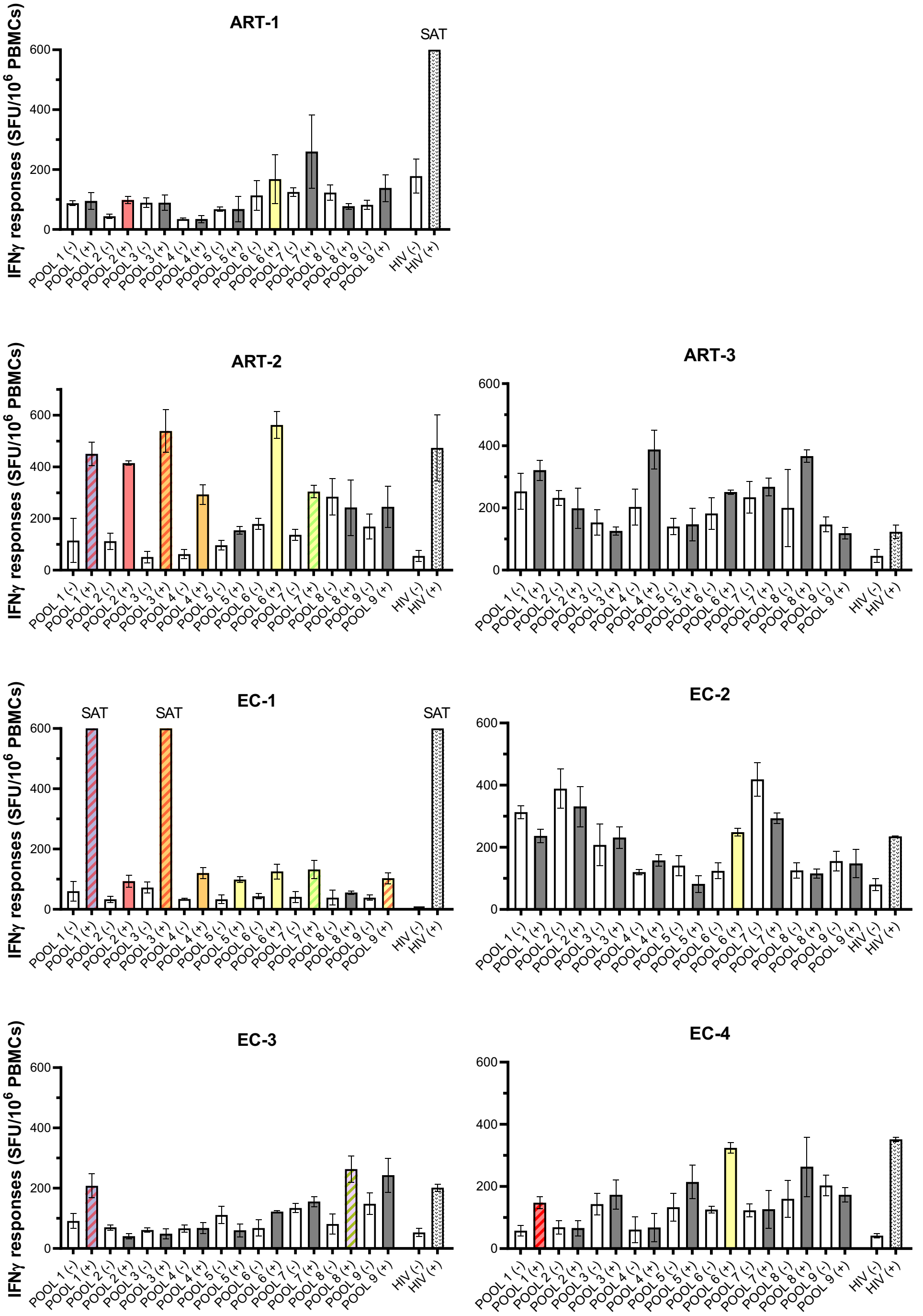


Supplementary Fig. 6 related to Fig. 3

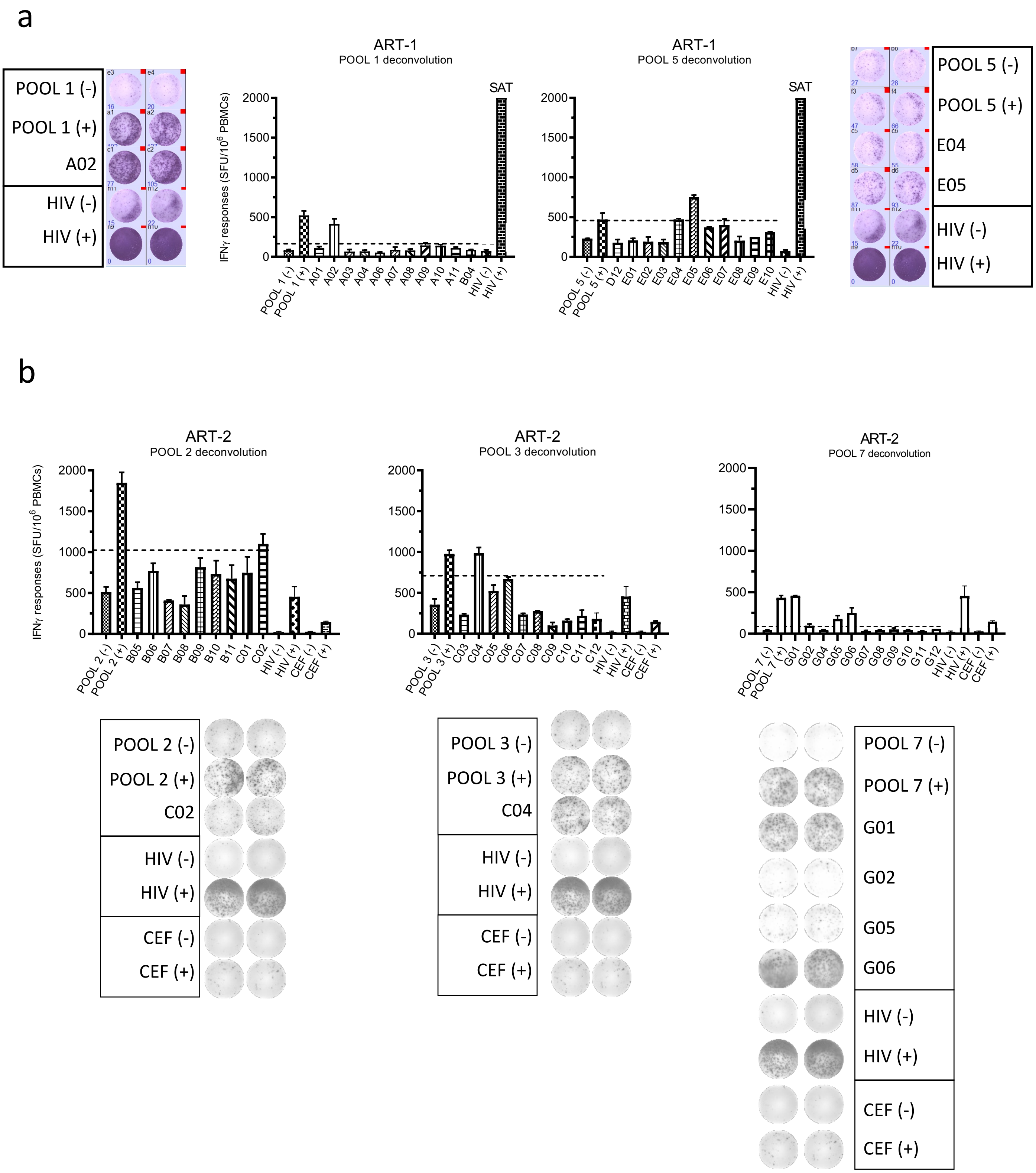


Supplementary Fig. 7 related to Fig. 3 1/4

c

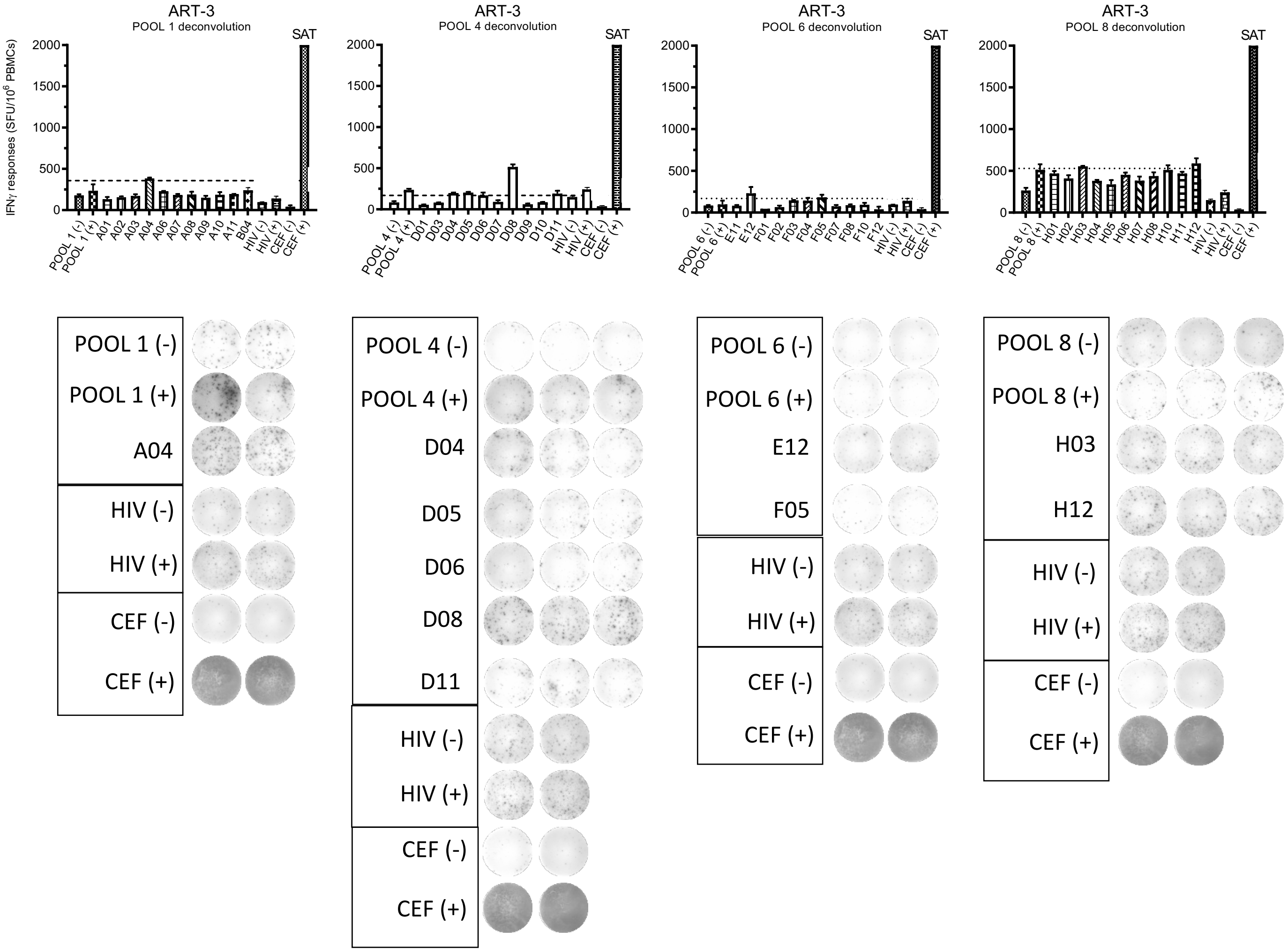

ART-3
POOL 1 deconvolution
ART-3
POOL 4 deconvolution
ART-3
POOL 6 deconvolution
ART-3
POOL 8 deconvolution
SAT
IFNγ responses (SFU/10^6 PBMCs)
POOL 1 (-)
POOL 1 (+)
A04
HIV (-)
HIV (+)
CEF (-)
CEF (+)
POOL 4 (-)
POOL 4 (+)
D04
D05
D06
D08
D11
POOL 6 (-)
POOL 6 (+)
E12
F05
POOL 8 (-)
POOL 8 (+)
H03
H12


d

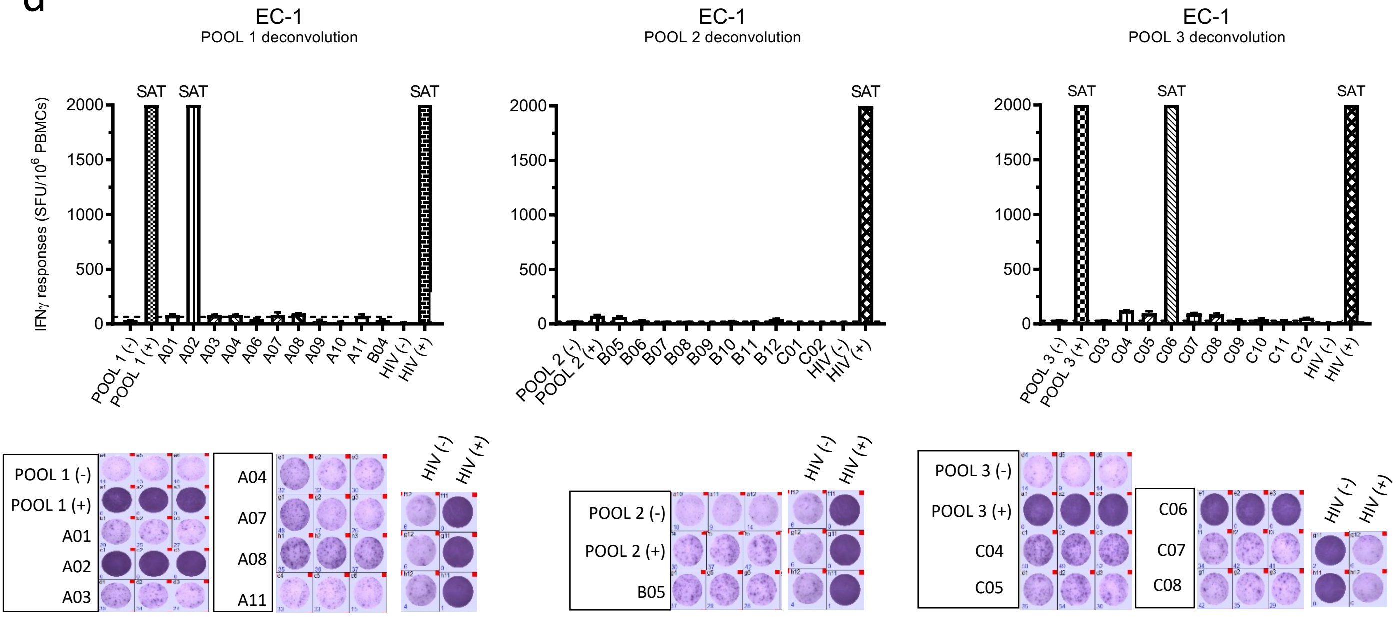

EC-1
POOL 1 deconvolution
EC-1
POOL 2 deconvolution
EC-1
POOL 3 deconvolution
SAT
IFNγ responses (SFU/10^6 PBMCs)
POOL 1 (-)
POOL 1 (+)
A01
A02
A03
A04
A07
A08
A11
HIV (-)
HIV (+)
POOL 2 (-)
POOL 2 (+)
B05
POOL 3 (-)
POOL 3 (+)
C04
C05
C06
C07
C08

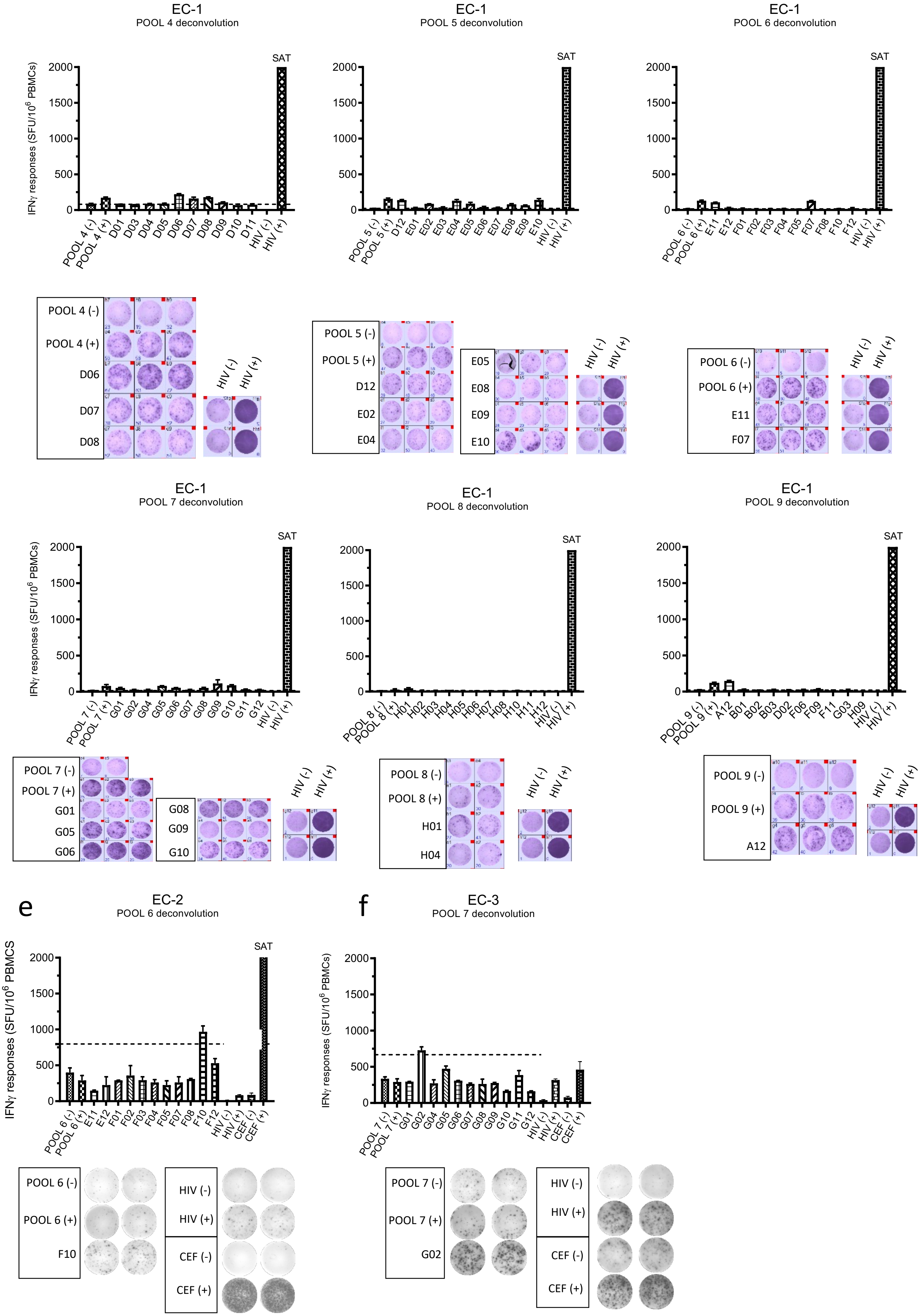


Supplementary Fig. 7 related to Fig. 3 3/4

g

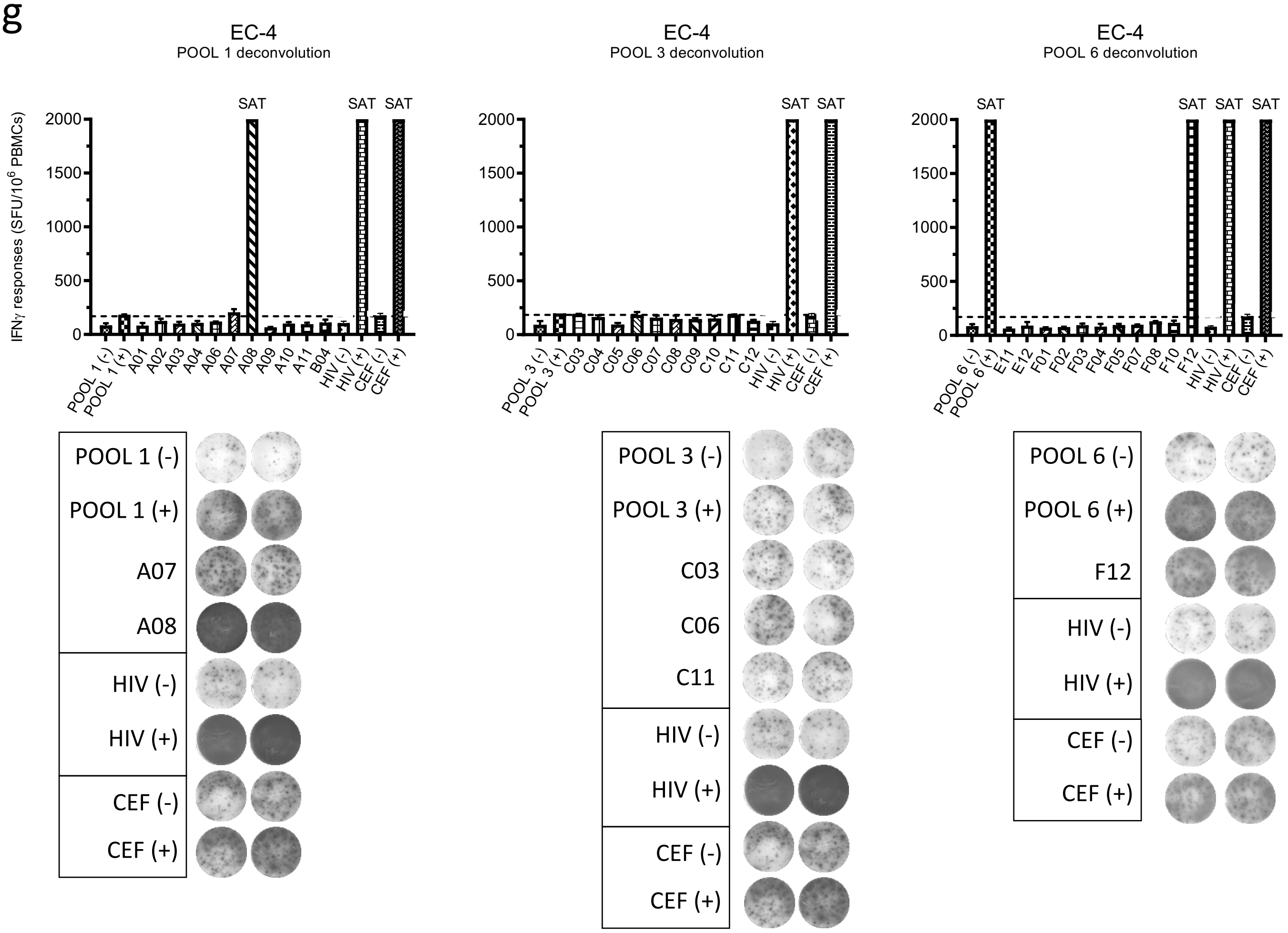


Supplementary Fig. 7 related to Fig. 3 4/4

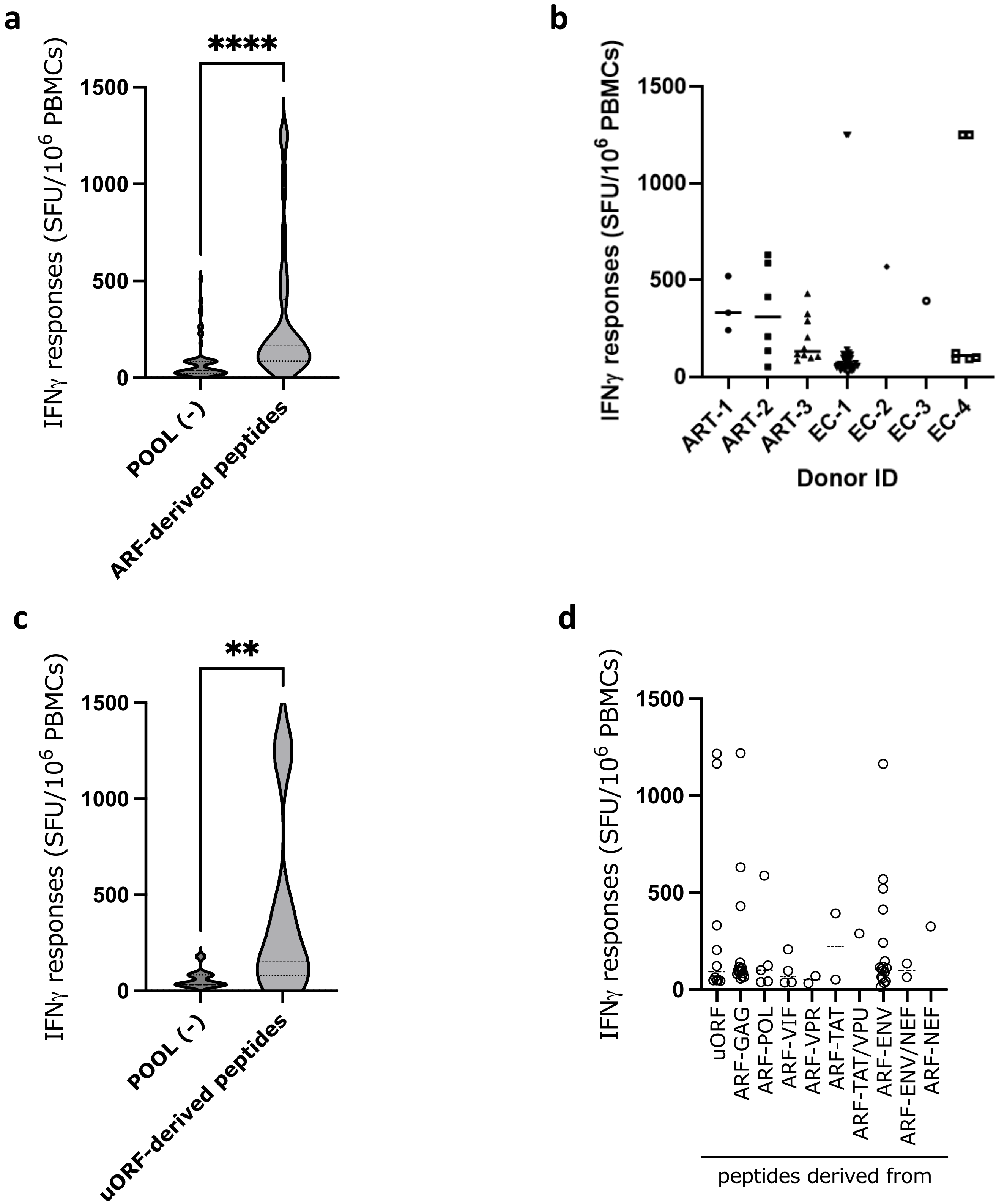


Supplementary Fig. 8 related to Fig. 3

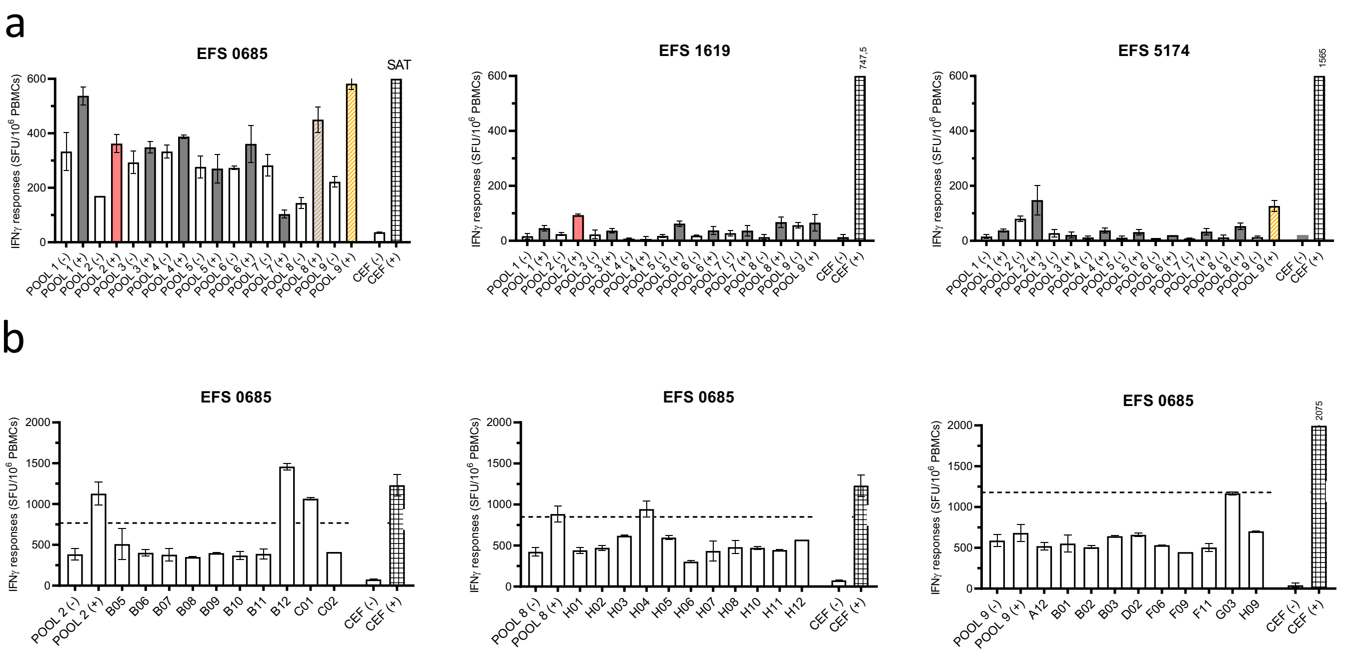


Supplementary Fig. 9 related to Fig. 3

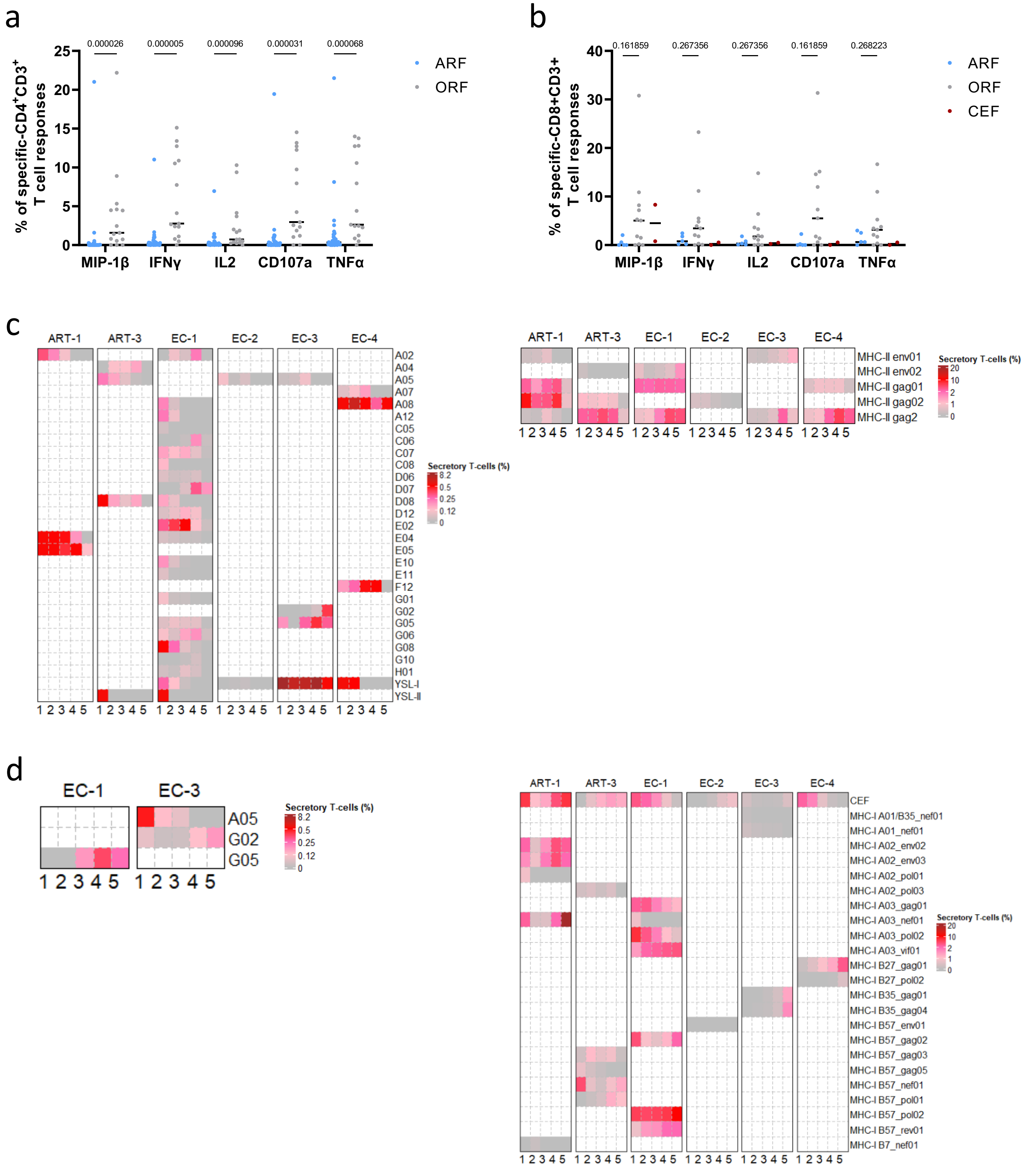


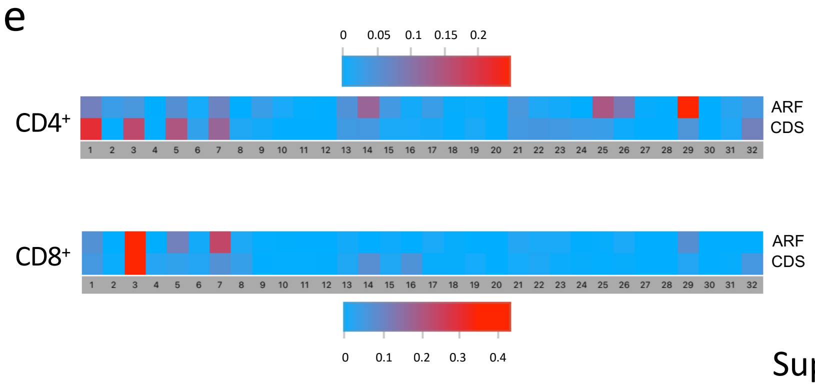


Category

| # | TNFα | CD107a | IFNγ | IL2 | MIP-1β | # | TNFα | CD107a | IFNγ | IL2 | MIP-1β |
|---|---|---|---|---|---|---|---|---|---|---|---|
| 1 | + | + | + | + | - | 17 | - | + | + | + | - |
| 2 | + | + | + | + | + | 18 | - | + | + | + | + |
| 3 | + | + | + | - | - | 19 | - | + | + | - | - |
| 4 | + | + | + | - | + | 20 | - | + | + | - | + |
| 5 | + | + | - | + | - | 21 | - | + | - | + | - |
| 6 | + | + | - | + | + | 22 | - | + | - | + | + |
| 7 | + | + | - | - | - | 23 | - | + | - | - | - |
| 8 | + | + | - | - | + | 24 | - | + | - | - | + |
| 9 | + | - | + | + | - | 25 | - | - | + | + | - |
| 10 | + | - | + | + | + | 26 | - | - | + | + | + |
| 11 | + | - | + | - | - | 27 | - | - | + | - | - |
| 12 | + | - | + | - | + | 28 | - | - | + | - | + |
| 13 | + | - | - | + | - | 29 | - | - | - | + | - |
| 14 | + | - | - | + | + | 30 | - | - | - | + | + |
| 15 | + | - | - | - | - | 31 | - | - | - | - | - |
| 16 | + | - | - | - | + | 32 | - | - | - | - | + |

Supplementary Fig. 10 related to Fig. 4

a

| Sample | Name of peptides | Reading frame | Specific-CD4$^{+}$ T cell response | Specific-CD8$^{+}$ T cell response |
|---|---|---|---|---|
| EC-1 | A02 | ARF | YES | NO |
| | C06 | ARF | YES | NO |
| | D07 | ARF | YES | NO |
| | G05 | ARF | NO | YES |
| | G06 | ARF | NO | NO |
| | YSL-I | ARF | YES | NO |
| | MHC-I A03_vif01 | CDS | YES | NO |
| | MHC-I B57_rev01 | CDS | YES | NO |
| | MHC-II gag2 | CDS | YES | NO |
| | CEF | non-HIV common viral peptides | YES | NO |

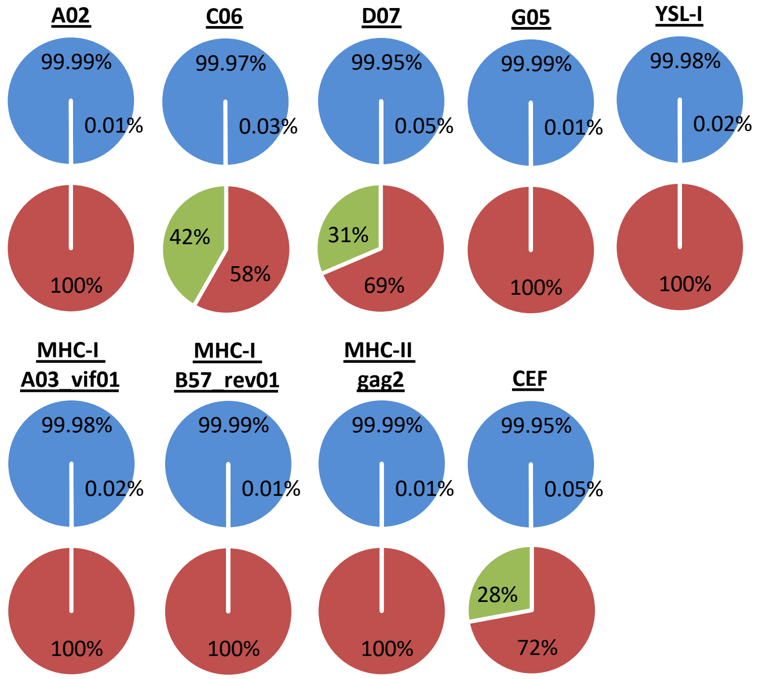


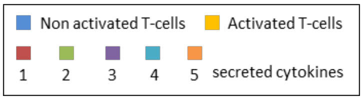


b

| Sample | Name of peptides | Reading frame | Specific-CD4$^{+}$ T cell response | Specific-CD8$^{+}$ T cell response |
|---|---|---|---|---|
| EC-3 | A05 | ARF | YES | NO |
| | G02 | ARF | YES | NO |
| | G05 | ARF | YES | NO |
| | YSL-I | ARF | YES | NO |
| | MHC-I B35_gag01 | CDS | NO | NO |
| | MHC-I B353gag04 | CDS | NO | NO |
| | MHC-II env01 | CDS | NO | YES |
| | MHC-II gag2 | CDS | YES | NO |
| | CEF | non-HIV common viral peptides | NO | NO |

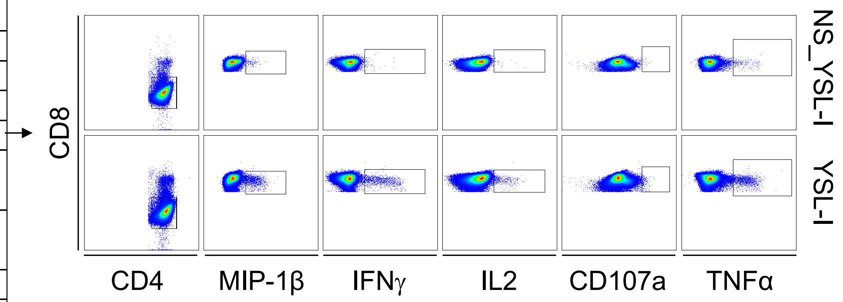


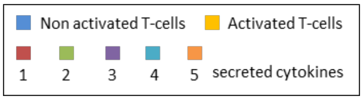


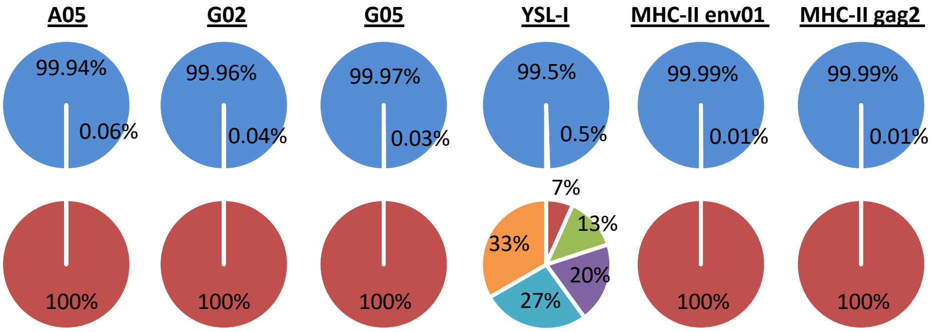


Supplementary Fig. 11

a

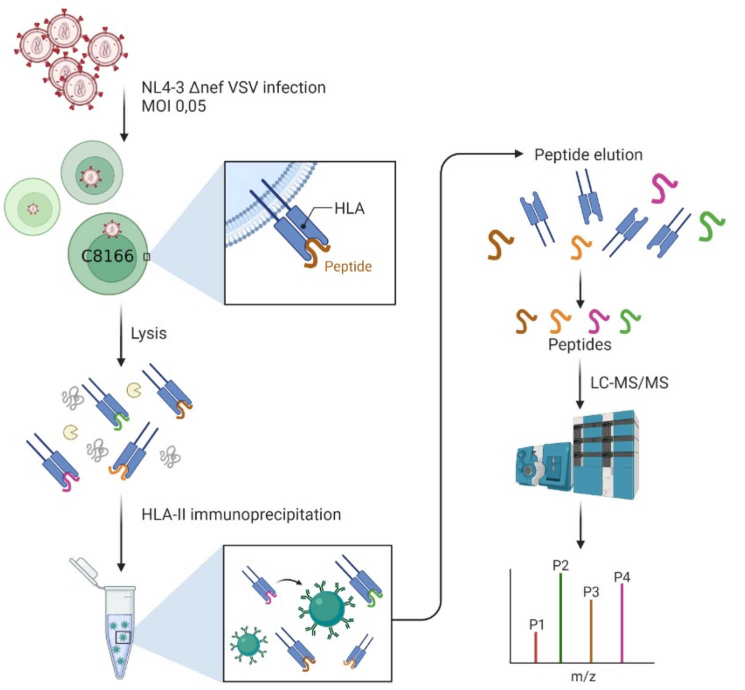


b

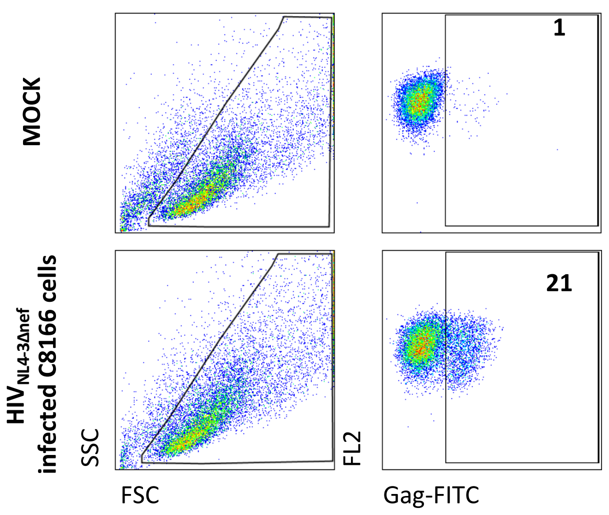


c

| Sample | HIV strain | % Infection | HLA type | HIV ligands | 5' Position (nt) | CDS |
|---|---|---|---|---|---|---|
| C8166 | NL4-3 ΔNEF | 20% | DRB1*03:01; DRB1*04:01; DQB1*02;DQB1*03:01 | RPEPTAPPEESFRFGEETTTPSQK | 2143 | *gag* |
| | | | | RPEPTAPPEESFRFGEETTTPSQKQ | 2143 | *gag* |
| | | | | SRPEPTAPPEESFRFGEETTTPSQK | 2140 | *gag* |
| | | | | ESFRFGEETTTPSQK | 2170 | *gag* |
| | | | | ESFRFGEETTTPSQKQEP | 2170 | *gag* |
| | | | | LQSRPEPTAPPEESFRFGEETTTPS | 2134 | *gag* |
| | | | | APPEESFRFGEETTTPSQK | 2158 | *gag* |
| | | | | APPEESFRFGEETTTPSQKQ | 2158 | *gag* |
| | | | | APPEESFRFGEETTTPSQKQEP | 2158 | *gag* |
| | | | | APPEESFRFGEETTTPSQKQE | 2158 | *gag* |
| | | | | SPEVIPMFSALSEGATP | 1282 | *gag* |
| | | | | VDRFYKTLRAEQASQEVK | 1678 | *gag* |
| | | | | DYVDRFYKTLRAEQASQEVK | 1672 | *gag* |

d

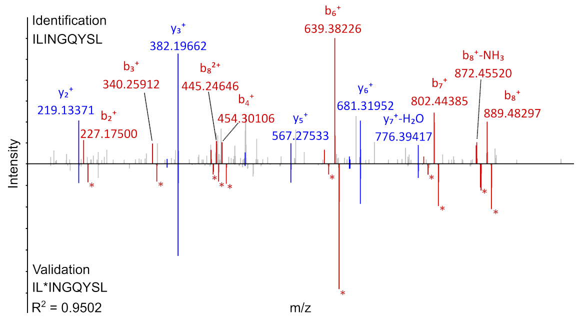


e

| position in HIV genome | | YSL-ARF | | | |
|---|---|---|---|---|---|
| Beginning | End | Phase | Nucletide-sequence | Size (aa) | Amino acid-sequence |
| 5' Stop 3181 | 3282 | 1 | ggtggggatttaccacaccagacaaaaaacatcagaaagaacctccattcctttggatgggttatgaactccatcctgataaatggacagtacagcctatag | 33 | GGDLPHQTKNIRKNLHSFGWVMNSILINGQYSL* |
| start codon 3222 | 3282 | 1 | ctccattcctttggatgggttatgaactccatcctgataaatggacagtacagcctatag | 19 | LHSFGWVMNSILINGQYSL* |

Supplementary Fig. 12 related to Fig. 5

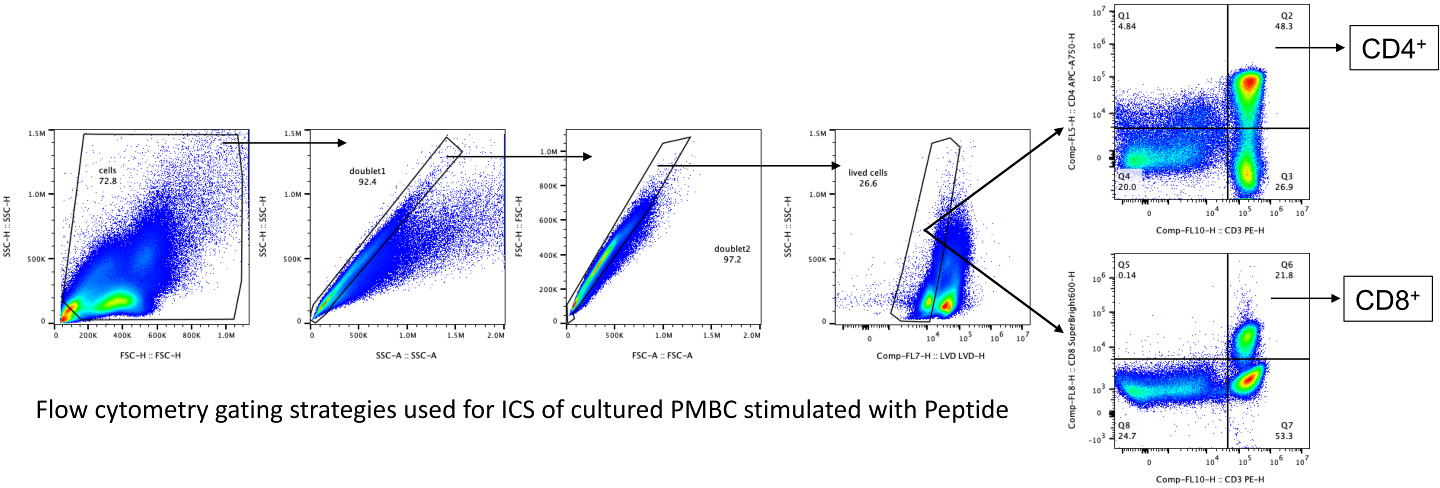


Flow cytometry gating strategies used for ICS of cultured PMBC stimulated with Peptide

Supplementary Fig. 13 related to Fig. 4

Supplementary Table 1

| Peptide name | CDS | Synthetised peptide sequence | Pool |
|---|---|---|---|
| MHC-II env01 | *env* | FEPIPIHYCAPAGFAILKC | HIV |
| MHC-II env02 | *env* | VSTVQCTHGIRPVVSTQLLL | HIV |
| B06 | *env* | KVSFEPIPIHYCAPAGFAILKC | HIV |
| B08 | *env* | KVSFEPIPIHYCAPAGFAILK | HIV |
| MHC-II env04 | *env* | TGEIIGDIRQAHCNISktqW | HIV |
| MHC-II gag01 | *gag* | EKAFSPEVIPMFSALSEGAT | HIV |
| D09 | *gag* | YVDRFYKTLRAEQASQEV | HIV |
| MHC-II gag02 | *gag* | CKTILKALGPAATLEEMMTA | HIV |

Supplementary Table 2

| | Peptide name | Short name of the peptide | Synthetised peptide sequence | Length (aa) | Pool |
|---|---|---|---|---|---|
| 1 | ARF-UTR2-A01 | A01 | LASIKLALSALSVCPSVV | 18 | 1 |
| 2 | ARF-UTR4-A02 | A02 | LWVLEIPQTLLVSVENL | 17 | 1 |
| 3 | ARF-UTR7-A03 | A03 | SQRRSLDAGLGLLKRARQEA | 20 | 1 |
| 4 | ARF-UTR8-A04 | A04 | TRERLSLDAGLGLLKSARQKRREL | 24 | 1 |
| 5 | ARF-UTR8-A05 | A05 | RQKRRELSRRRTRLAEGLGLLKC | 23 | 9 |
| 6 | ARF-UTR9-A07 | A07 | EEISRRRTRLAEARTARGE | 19 | 1 |
| 7 | ARF-UTR9-A08 | A08 | GEGRRLVSTPKILTSGG | 17 | 1 |
| 8 | ARF-POL1-A11 | A11 | QYWMWVMHIFQFP | 13 | 1 |
| 9 | ARF-POL2-A12 | A12 | SRLLGSSIRNTTSRRVKKEKI | 21 | 9 |
| 10 | ARF-POL8-B05 | B05 | LTQQIRRLSYKQFI | 14 | 2 |
| 11 | ARF-POL15-C03 | C03 | HKSSAKKKSKDHQGLWKTDGR | 21 | 3 |
| 12 | ARF-GAG2-C04 | C04 | SVKARGKEKISIKTYSMGK | 19 | 3 |
| 13 | ARF-GAG2-C05 | C05 | TYSMGKQGARTI | 12 | 3 |
| 14 | ARF-GAG2-C06 | C06 | GARTIRSQSWPVRNIRRL | 18 | 3 |
| 15 | ARF-GAG5-C07 | C07 | TNTGTATTIPSDRIRRT | 17 | 3 |
| 16 | ARF-GAG6-C08 | C08 | KHQKAVDKYWDSYNHPF | 17 | 3 |
| 17 | ARF-GAG8-C11 | C11 | IASSACRAYCTRPDERTKGK | 20 | 3 |
| 18 | ARF-GAG9-C12 | C12 | QELLVPFRNK | 10 | 3 |
| 19 | ARF-GAG13-D04 | D04 | DKDQRNPLETM | 11 | 4 |
| 20 | ARF-GAG14-D05 | D05 | SKSRASFTGGKKLDDRNLVGPK | 22 | 4 |
| 21 | ARF-GAG15-D06 | D06 | QKPCWSRMRTQIVRLF | 16 | 4 |
| 22 | ARF-GAG16-D07 | D07 | YFKSIRTSSYTRRNDDSM | 18 | 4 |
| 23 | ARF-GAG17-D08 | D08 | CRKAILGTKERLLSVSIVAKK | 21 | 4 |
| 24 | ARF-GAG20-D11 | D11 | EKGLLEMWKRRTPNERLY | 18 | 4 |
| 25 | ARF-ENV2-D12 | D12 | SVVLQKNCGSQSIMGYLCGK | 20 | 5 |
| 26 | ARF-ENV2-E02 | E02 | KHMIQRYIMFGPHMPVYPQTPT | 22 | 5 |
| 27 | ARF-ENV4-E04 | E04 | VQRYPLSQFPYIIVPRLVLRF | 21 | 5 |
| 28 | ARF-ENV5-E05 | E05 | ANSHTLLYPGWFCDSKV | 17 | 5 |
| 29 | ARF-ENV7-E08 | E08 | ASSYNSTAVKWQSSRRRGSN | 20 | 5 |
| 30 | ARF-ENV8-E09 | E09 | SLINPQEGTQKL | 12 | 5 |
| 31 | ARF-ENV9-E10 | E10 | KQCMPLPSVEKLDVHQILQGYY | 22 | 5 |
| 32 | ARF-ENV10-E11 | E11 | MFIKYYRAAINKRWW | 15 | 6 |
| 33 | ARF-ENV11-E12 | E12 | QEMVVITRPRSSDLEEEI | 18 | 6 |
| 34 | ARF-ENV18-F05 | F05 | ELLRRNSICCN | 11 | 6 |

Supplementary Table 2

| | | | | | |
|---|---|---|---|---|---|
| 35 | ARF-ENV19-F07 | F07 | GATASVATHSLGHQTAPGKSP | 21 | 6 |
| 36 | ARF-ENV20-F10 | F10 | RINSSWEFGVALENSFAPL | 19 | 6 |
| 37 | ARF-ENV23-F12 | F12 | YTPYLKNRRTNKKRMNKNYWNW | 22 | 6 |
| 38 | ARF-ENV25-G01 | G01 | NDSRRLDRFKNSFYCTFYSK | 20 | 7 |
| 39 | ARF-peptide leader TAT-G02 | G02 | IELGRDTHHYRFRPASQPRGDP | 22 | 7 |
| 40 | ARF-ENV/NEF1-G05 | G05 | KNKTGLRKGFAIRWVASGQKVV | 22 | 7 |
| 41 | ARF-VIF1-G06 | G06 | QHIGVYIQEKETGIWVRESP | 20 | 7 |
| 42 | ARF-VIF3-G08 | G08 | QGRISTVLGTSSINSTKKDKAT | 22 | 7 |
| 43 | ARF-VIF5-G09 | G09 | QRIDGTSPRRPRATEGAIQ | 19 | 7 |
| 44 | ARF-VPR2-G10 | G10 | RSLRMKLLDIFLGHGFMA | 18 | 7 |
| 45 | ARF-VPR4-H01 | H01 | EFCNNCCLFISELGVNIAE | 19 | 8 |
| 46 | ARF-TAT/VPU1-H03 | H03 | KVLLSLPSLFHNKRLRHLLWQ | 21 | 8 |
| 47 | ARF-NEF7 | H12 | FTPRKDKISLICGSTTHK | 18 | 8 |

Supplementary Table 3

| | HLA-restriction | CDS | Sequence (aa) | Peptide name |
|---|---|---|---|---|
| 1 | A*01/B*35 | Nef | YFPDWQNYT | MHC-I A01/B35_NEF01 |
| 2 | A*01 | Env | RRGWEVLKY | MHC-I A01_ENV01 |
| 3 | A*01 | Rev | ISERILSTY | MHC-I A01_REV01 |
| 4 | A*01 | Gag | GSEELRSLY | MHC-I A01_GAG01 |
| 5 | A*01 | Nef | WRFDSRLAFH | MHC-I A01_NEF01 |
| 6 | B*35 | Gag | PPIPVGDIY | MHC-I B35_GAG01 |
| 4 | B*35 | Pol | HPDIVIYQY | MHC-I B35_POL01 |
| 7 | B*35 | Nef | VPLRPMTY | MHC-I B35_NEF01 |
| 8 | B*35 | Env | DPNPQEVVL | MHC-I B35_ENV01 |
| 9 | B*35/B*51 | Pol | IPLTEEAEL | MHC-I B35/B51_POL01 |
| 10 | B*35 | Gag | NSSKVSQNY | MHC-I B35_GAG02 |
| 10 | B*35 | Env | TAVPWNASW | MHC-I B35_ENV02 |
| 11 | B*35 | Pol | TVLDVGDAY | MHC-I B35_POL02 |
| 12 | B*35 | Pol | VPLDEDFRKY | MHC-I B35_POL03 |
| 13 | B*35 | Env | VPVWKEATTTL | MHC-I B35_ENV03 |
| 14 | B*35 | Gag | WASRELERF | MHC-I B35_GAG03 |
| 15 | B*35 | Gag | HPVHAGPIA | MHC-I B35_GAG04 |
| 16 | B*35 | Gag | NPPIPVGDIY | MHC-I B35_GAG05 |
| 16 | A*02 | Gag | SLYNTVATL | MHC-I A02_GAG01 |
| 17 | A*02 | Pol | ILKEPVHGV | MHC-I A02_POL01 |
| 18 | A*02 | Nef | VLEWRFDSRL | MHC-I A02_NEF01 |
| 19 | A*02 | Nef | PLTFGWCYKL | MHC-I A02_NEF02 |
| 20 | A*02 | Vpr | AIIRILQQL | MHC-I A02_VPR01 |
| 21 | A*02 | Pol | ALVEICTEM | MHC-I A02_POL02 |
| 21 | A*02 | Gag | FLGKIWPSYK | MHC-I A02_GAG02 |
| 22 | A*02 | Nef | GAFDLSFFL | MHC-I A02_NEF03 |
| 23 | A*02 | Pol | LVGPTPVNI | MHC-I A02_POL03 |
| 24 | A*02 | Env | RGPGRAFVTI | MHC-I A02_ENV01 |
| 25 | A*02 | Env | RIRQGLERA | MHC-I A02_ENV02 |
| 26 | A*02 | Env | SLLNATDIAV | MHC-I A02_ENV03 |
| 26 | A*02 | Pol | VIYQYMDDL | MHC-I A02_POL04 |
| 27 | A*02 | Gag | YVDRFYKTL | MHC-I A02_GAG03 |
| 28 | A*03 | Gag | RLRPGGKKK | MHC-I A03_GAG01 |
| 29 | A*03 | Gag | KIRLRPGGK | MHC-I A03_GAG02 |
| 30 | A*03 | Pol | AIFQSSMTK | MHC-I A03_POL01 |
| 31 | A*03 | Env | RLRDLLLIVTR | MHC-I A03_ENV01 |
| 32 | A*03 | Nef | QVPLRPMTYK | MHC-I A03_NEF01 |
| 32 | A*03 | Nef | AVDLSHFLK | MHC-I A03_NEF02 |
| 33 | A*03 | Pol | AVFIHNFKRK | MHC-I A03_POL02 |
| 34 | A*03 | Vif | RIRTWKSLVK | MHC-I A03_VIF01 |
| 35 | A*03 | Gag | RLRPGGKKKY | MHC-I A03_GAG03 |
| 36 | A*03 | Pol | RMRGAHTNDVK | MHC-I A03_POL03 |

Supplementary Table 3

| | | | | |
|---|---|---|---|---|
| 37 | A*03 | Vif | RIRTWKSLVK | MHC-I A03_VIF02 |
| 37 | A*24 | Gag | KYKLKHIVW | MHC-I A24_GAG01 |
| 38 | A*24 | Nef | RYPLTFGW | MHC-I A24_NEF01 |
| 39 | B*57 | Vpr | AVRHFPRIW | MHC-I B57_VPR01 |
| 40 | B*57 | Gag | FSPEVIPMF | MHC-I B57_GAG01 |
| 41 | B*57 | Nef | HTQGYFPDW | MHC-I B57_NEF01 |
| 42 | B*57 | Vif | ISKKAKGWF | MHC-I B57_VIF01 |
| 43 | B*57 | Gag | ISPRTLNAW | MHC-I B57_GAG02 |
| 43 | B*57 | Pol | IVLPEKDSW | MHC-I B57_POL01 |
| 44 | B*57 | Gag | KAFSPEVI | MHC-I B57_GAG02 |
| 45 | B*57 | Gag | KAFSPEVIPMF | MHC-I B57_GAG03 |
| 46 | B*57 | Gag | QASQEVKNW | MHC-I B57_GAG04 |
| 47 | B*57 | Pol | STTVKAACWW | MHC-I B57_POL02 |
| 48 | B*57 | Gag | TSTLQEQIGW | MHC-I B57_GAG05 |
| 48 | B*57 | Rev | KAVRLIKFLY | MHC-I B57_REV01 |
| 49 | B*27 | Pol | KTAVQMAVF | MHC-I B27_POL01 |
| 50 | B*27 | Gag | KRWIILGLNK | MHC-I B27_GAG01 |
| 51 | B*27 | Gag | FRNQRKTVK | MHC-I B27_GAG02 |
| 52 | B*27 | Gag | IRLRPGGKK | MHC-I B27_GAG03 |
| 53 | B*27 | Pol | KRKGGIGGY | MHC-I B27_POL02 |
| 54 | B*27 | Nef | RRQDILDLWI | MHC-I B27_NEF01 |
| 54 | B*27 | Vpr | VRHFPRIWL | MHC-I B27_VPR01 |
| 55 | B*27 | Nef | LRPMTYKAA | MHC-I B27_NEF02 |
| 56 | B*27 | Gag | QRGNFRNQRK | MHC-I B27_GAG04 |
| 57 | B*27 | Gag | RRWIQLGLQK | MHC-I B27_GAG05 |
| 58 | B*51 | Gag | GRRGWEALKY | MHC-I B51_GAG01 |
| 59 | B*51 | Pol | TAFTIPSI | MHC-I B51_POL01 |
| 59 | B*51 | Env | RAIEAQQHL | MHC-I B51_ENV01 |
| 60 | B*51 | Vpr | EAVRHFPRI | MHC-I B51_VPR01 |
| 61 | B*51 | Pol | EKEGKISKI | MHC-I B51_POL01 |
| 62 | B*51 | Vif | IPLGDAKLII | MHC-I B51_VIF01 |
| 63 | B*51 | Env | LPCRIKQII | MHC-I B51_ENV02 |
| 64 | B*51 | Pol | RPLVTIKI | MHC-I B51_POL03 |
| 65 | B*51 | Gag | RPGGKKKYKL | MHC-I B51_GAG02 |
| 65 | B*44 | Pol | EEMNLPGRW | MHC-I B44_POL01 |
| 66 | B*44 | Env | AENLWVTVY | MHC-I B44_ENV01 |
| 67 | B*44 | Gag | AEQASQDVKNW | MHC-I B44_GAG01 |
| 68 | B*44 | Gag | LYNTVATLY | MHC-I B44_GAG02 |
| 69 | B*44 | Gag | RDYVDRFYKTL | MHC-I B44_GAG03 |
| 70 | B*07 | Gag | SPRTLNAWV | MHC-I B07_GAG01 |
| 70 | B*07 | Env | RPNNNTRKSI | MHC-I B07_ENV01 |
| 71 | B*07 | Env | IPRRIRQGL | MHC-I B07_ENV02 |
| 72 | B*07 | Nef | TPGPGVRYPL | MHC-I B07_NEF01 |
| 73 | B*07 | Gag | GPGHKARVL | MHC-I B07_GAG02 |
| 74 | B*07 | Pol | SPAIFQSSM | MHC-I B07_POL01 |

Supplementary Table 3

| | | | | |
|---|---|---|---|---|
| 75 | B*07 | Vpr | FPRIWLHGL | MHC-I B07_VPR01 |
| 76 | DRβ1*04 | Gag | EKAFSPEVIPMFSALSEGAT | MHC-II gag01 |
| 76 | DRβ1*04 | Gag | CKTILKALGPAATLEEMMTA | MHC-II gag02 |
| 77 | DRβ1*04 | Gag | NKIVRMYSPTSILDIRQGPK | MHC-II gag2 |
| 78 | DRβ1*01 | Env | FEPIPIHYCAPAGFAILKC | MHC-II env01 |
| 79 | DRβ1*01 | Env | VSTVQCTHGIRPVVSTQLLL | MHC-II env02 |
| 80 | DRβ1*01 | Env | RPNNNTRKSINIGPGRALYT | MHC-II env03 |
| 81 | DRβ1*01 | Env | TGEIIGDIRQAHCNISKTQW | MHC-II env04 |
